# Local adaptation, plasticity, and evolved resistance to hypoxic cold stress in high-altitude deer mice

**DOI:** 10.1101/2024.06.21.600120

**Authors:** Naim M. Bautista, Nathanael D. Herrera, Ellen Shadowitz, Oliver H. Wearing, Zachary A. Cheviron, Graham R. Scott, Jay F. Storz

## Abstract

A fundamental question in evolutionary biology concerns the relative contributions of phenotypic plasticity vs. local adaptation (genotypic specialization) in enabling wide-ranging species to inhabit diverse environmental conditions. Here we conduct a long-term hypoxia acclimation experiment to assess the relative roles of local adaptation and plasticity in enabling highland and lowland deer mice (*Peromyscus maniculatus*) to sustain aerobic thermogenesis at progressively increasing elevations. We assessed the relative physiological performance capacities of highland and lowland natives as they were exposed to progressive, stepwise increases in hypoxia, simulating the gradual ascent from sea level to an elevation of 6000 m. The final elevation of 6000 m far exceeds the highest attainable elevations within the species’ range, and therefore tests the animals’ ability to tolerate levels of hypoxia that surpass the prevailing conditions within their current distributional limits. Our results demonstrate that highland natives exhibit superior thermogenic capacities at the most severe levels of hypoxia, suggesting that the species’ broad fundamental niche and its ability to inhabit such a broad range of elevational zones is attributable to a combination of genetically based local adaptation and plasticity. Transcriptomic and physiological measurements identify evolved changes in the acclimation response to hypoxia that contribute to the enhanced thermogenic capacity of highland natives.

**SIGNIFICANCE STATEMENT:** In species that are distributed across steep environmental gradients, the ability to inhabit a broad range of conditions may be attributable to local adaptation and/or a generalized acclimatization capacity (phenotypic plasticity). By experimentally acclimating highland and lowland deer mice (*Peromyscus maniculatus*) to progressively increasing levels of hypoxia during a simulated ascent to 6000 m, we assessed the relative roles of evolved and plastic changes in thermogenic capacity. At especially severe levels of hypoxia, the superior thermogenic performance of highland natives relative to lowland conspecifics suggests that the broad fundamental niche of deer mice is largely attributable to local adaptation to different elevational zones, including evolved plasticity in gene expression and respiratory traits.

## INTRODUCTION

A fundamental question in evolutionary biology concerns the relative contributions of phenotypic plasticity vs. local adaptation (genotypic specialization) in enabling wide-ranging species to inhabit diverse environmental conditions. In species that are distributed across environmental gradients, the interplay of plasticity and local adaptation can influence geographic range limits, range overlap with other species, and population persistence in the face of changing environmental conditions (Case & Taper, 2000; Chevin & Lande, 2011; Goldberg & Price, 2022; Holt & Gaines, 1992; Kirkpatrick & Barton, 1997; Thibert-Plante & Hendry, 2011). In species that are distributed across elevational gradients, plasticity in environmental tolerance can be expected to influence the ability of species to track upward shifts in temperature isotherms as a result of climate change. Since upward range shifts entail exposure to a progressively lower partial pressure of O_2_ (*P*O_2_), air-breathing animals must contend with associated constraints on aerobic metabolism when colonizing especially high elevations.

The deer mouse, *Peromyscus maniculatus*, is an excellent study species for investigating questions about plasticity and the evolution of environmental tolerance, as it has the broadest elevational distribution of any North American mammal. This quintessential generalist species is continuously distributed across habitats ranging from lowland desert and prairie to the high alpine and can be locally abundant on the summits of ∼4300 m peaks in the Southern Rockies and other mountain ranges in western North America. For small endotherms living at such elevations, a capacity for sustained aerobic thermogenesis may often be critical for survival during periods of extreme cold (Hayes & O’Connor, 1999). When challenged with the combined stressors of cold and hypoxia, mice that can attain an especially high thermogenic capacity (the maximum rate of O_2_-consumption [*V*O_2max_] elicited by cold exposure) can maintain constant body temperature at lower ambient temperatures than others. Mice with higher thermogenic capacities can therefore remain active in the cold for longer periods of time, providing increased opportunities for foraging, finding mates, and other fitness-relevant activities (Sears et al., 2006). Common-garden experiments have revealed that high-altitude deer mice exhibit greater thermogenic capacities in hypoxia than lowland conspecifics, and these population differences in aerobic performance are at least partly genetically based (Cheviron et al., 2012; Cheviron et al., 2013; Cheviron et al., 2014; Lui et al., 2015; Tate et al., 2017; Tate et al., 2020).

Here we conduct a long-term hypoxia-acclimation experiment to assess the relative roles of local adaptation and plasticity in enabling highland deer mice to sustain aerobic thermogenesis at simulated elevations that span the species’ current range and exceed the upper limits. We compare the relative physiological performance capacities of highland and lowland natives as they are exposed to progressive, stepwise increases in hypoxia, simulating a gradual ascent from sea level to 6000 m. The final elevation of 6000 m far exceeds the highest attainable elevations within the geographical range of deer mice and therefore tests the animals’ ability to tolerate levels of hypoxia that surpass the prevailing conditions within their current distributional limits. If tolerance to hypoxic cold stress is primarily plastic, then long-term acclimation to gradually increasing hypoxia should enable highlanders and lowlanders to sustain similar performance capacities at the most extreme elevations. By contrast, if such tolerance primarily reflects an outcome of genetically based local adaptation, then highlanders would be expected to outperform equally well-acclimated lowlanders at elevations above some specific threshold.

## RESULTS AND DISCUSSION

### Highland mice more effectively mitigate hypoxia-induced decrements in thermogenic performance

We subjected highland and lowland deer mice to a common-garden acclimation trial involving discrete reductions in *P*O_2_ that simulated a gradual ascent from 0 to 6000 m above sea level over an 7-wk period (**Fig. 1A**). At each 1000 m increment in simulated elevation, we measured thermogenic *V̇*O_2max_ in conjunction with numerous subordinate traits, and we contrasted the measurements in acclimated mice to measurements in unacclimated controls from each population that were maintained in normoxia for the duration of the experiment. The general expectation is that measures of whole-animal aerobic performance such as *V̇*O_2max_ will progressively decline as a function of increasing elevation, but these hypoxia-induced decrements in performance may be partially mitigated by plasticity (as revealed by a difference between acclimated mice and unacclimated controls from the same population). Decrements in performance may also be partially mitigated by genetic adaptation to hypoxia in highland mice, which would be evidenced by higher *V̇*O_2max_ in highlanders relative to lowlanders (McClelland & Scott, 2019; Storz & Scott, 2019). Results revealed that acclimated highlanders sustained higher *V̇*O_2max_ at the most severe levels of hypoxia (4000-6000 m) in comparison with acclimated and unacclimated lowlanders (**Fig. 1B; Table S1)**. The fact that even unacclimated highlanders sustained higher *V̇*O_2max_ on average than fully acclimated lowlanders at 6000 m provides compelling evidence of local adaptation. At elevations >4000 m, acclimated highlanders maintained a significantly higher *V̇*O_2max_ than unacclimated controls, demonstrating that plasticity further mitigates the performance decrement in hypoxia, consistent with predictions of the ‘beneficial acclimation/adaptive plasticity’ hypothesis. In lowlanders, however, the fact that acclimated and unacclimated mice exhibited essentially identical *V̇*O_2max_ at 6000 m reveals the limits of compensatory plasticity under the most extreme levels of hypoxia.

**Figure 1.**
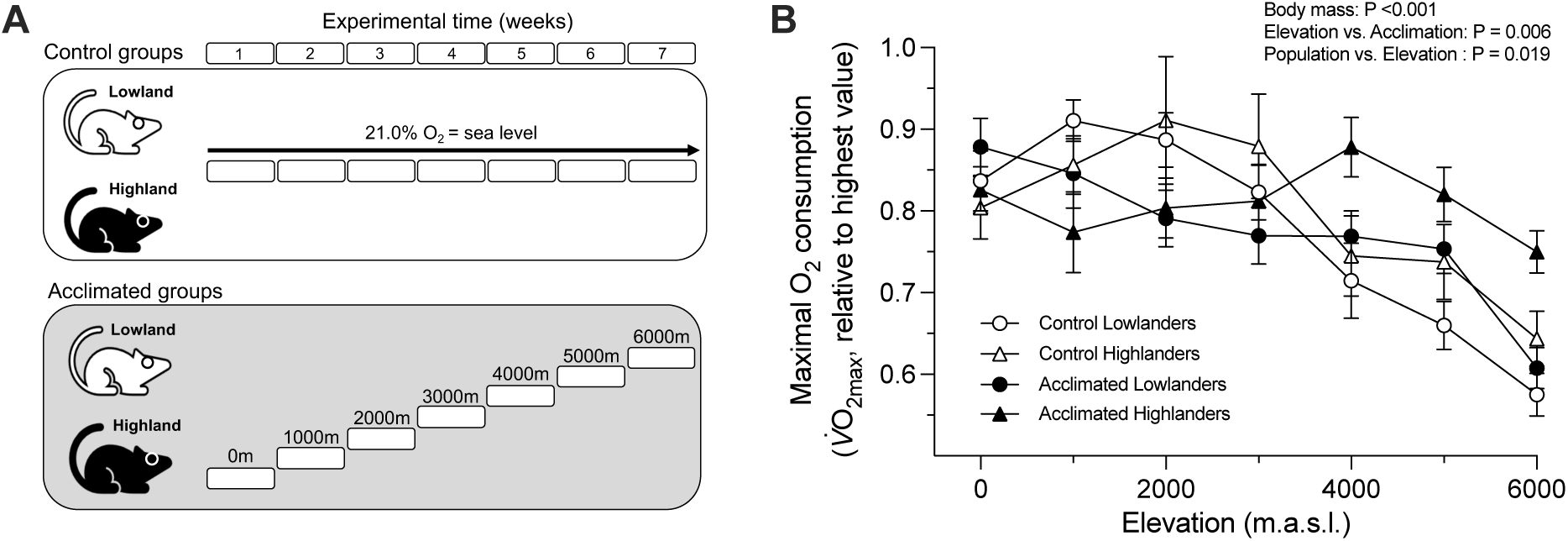
Thermogenic *V̇*O_2max_. Thermogenic *V̇*O_2max_ in acclimated and nonacclimated (control) deer mice under progressively increasing levels of hypoxia, from sea level to the equivalent of 6000 m. **A**) Experimental design. Top: highland and lowland mice in control groups were maintained in normobaric normoxia (equivalent to sea level) for the duration of the experimental period. Bottom: Highlanders and lowlanders in the treatment groups were acclimated to increasing levels of hypoxia that simulated a gradual ascent from sea level up to 6000m. **B**) At the highest elevations (4000-6000 m above sea level), highland natives sustain a higher thermogenic *V̇*O_2max_ than lowlanders in both acclimation and control groups, reflecting evolved differences in performance capacities in hypoxia. *V̇*O_2max_ was determined by acute exposure to cold (-5°C) heliox at *P*O_2_’s that progressively diminished with each increment of 1000 m during the simulated ascent. Data are expressed as values relative to the highest *V̇*O_2max_ attained by the mice. Mean ± s.e.m., N = 14-16 mice per data point. Statistical significance was considered with an *α* ≤ 0.05.

These findings suggest that the broad fundamental niche of deer mice and the species’ ability to inhabit such a broad range of elevational zones is largely attributable to local adaptation. The results also reveal evidence for the evolution of plasticity, as highland natives exhibited an enhanced acclimation response relative to lowlanders, enabling them to more effectively mitigate the decline in *V̇*O_2max_ under extreme levels of hypoxia corresponding to elevations that exceed the upper limits of the species’ range (5000-6000 m). The increased magnitude of hypoxia-induced plasticity in *V̇*O_2max_ of highlanders relative to lowlanders is consistent with results of previous common-garden experiments in which deer mice were acclimated to a uniform level of hypoxia over 6-8 weeks (Tate et al., 2017; Tate et al., 2020).

### Variation in subordinate traits that potentially contribute to the evolved enhancement of *V̇*O_2max_ in highland mice

Since *V̇*o_2max_ is an integrated measure of whole-animal physiological performance that reflects the flux capacity of the O_2_-transport pathway, population differences in *V̇*o_2max_ could potentially reflect evolved and/or plastic changes in any number of respiratory or circulatory steps in the pathway (McClelland & Scott, 2019; Scott & Dalziel, 2021; Storz & Scott, 2019; Storz, Scott, et al., 2010). At elevations >4000 m, acclimated and unacclimated highlanders exhibited consistently higher breathing frequencies (*ƒ*_R_) at *V̇*o_2max_ in comparison with lowlanders in the same experimental groups (**Fig. 2A; Table S1**). This was associated with an increase in total ventilation (*V̇*_E_) in highlanders, which could help maintain higher alveolar *P*O_2_ (thereby increasing O_2_ diffusion into the blood), but the variation in *V̇*_E_ was not significant. Increased *ƒ*_R_ also increases the respiratory excretion of metabolically produced CO_2_, consistent with the higher *V̇*CO_2_ observed in acclimated highland mice (**Fig. 2B**). None of the other measured respiratory traits exhibited consistent differences between highlanders and lowlanders (**Fig. 2C-H**). The lack of variation in air convection requirements among groups suggests that the variation in ventilation contributed to the observed variation in *V̇*o_2max_ and *V̇*CO_2_.

**Figure 2.**
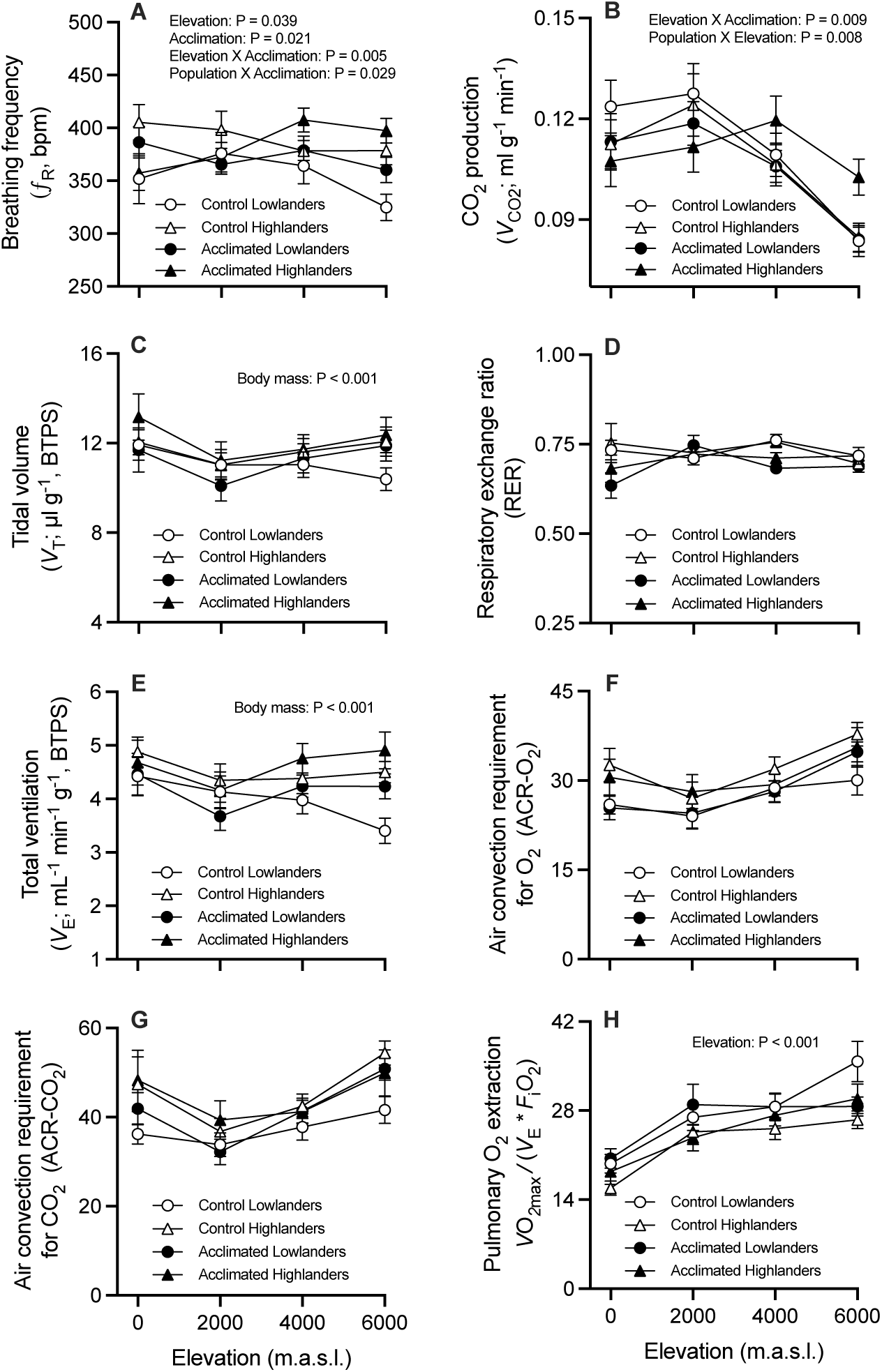
Ventilatory variables. Variation in ventilatory traits measured at *V̇*O_2max_ in acclimated and control mice. **A)** Breathing frequency; **B)** Carbon dioxide production; **C)** Tidal volume; **D)** Respiratory exchange ratio (*V̇*CO_2_/*V̇*O_2_); **E)** Total ventilation; **F)** Air convection requirement for oxygen (V_E_/*V̇*O_2max_); **G)** Air convection requirement for carbon dioxide (V_E_/*V̇*CO_2_); and **H)** Pulmonary oxygen extraction *V̇*O_2max_/(VE*F_i_O_2_). Statistical significance was considered with an *α* ≤ 0.05. N = 14-16 mice per data point.

In the same set of experimental mice, we also measured traits related to circulatory O_2_ transport that could potentially contribute to the observed population differences in *V̇*O_2max_ (**Fig. 3; Table S1**). As expected, exposure to gradually increasing levels of hypoxia (lower inspired *P*O_2_), resulted in corresponding reductions in arterial O_2_ saturation (*S*_a_O_2_) in both highlanders and lowlanders (**Fig. 3A**). Within both groups, the hypoxia-induced decrements in *S*_a_O_2_ were less severe in acclimated mice relative to unacclimated controls, especially at the highest elevations, which presumably reflects plasticity in respiratory and/or circulatory traits that partially compensate the lower inspired *P*O_2_. At 6000 m, acclimated lowlanders maintained a higher *S*_a_O_2_ relative to unacclimated lowlanders (80.6% vs. 73.9%, respectively), and acclimated highlanders maintained a higher *S*_a_O_2_ relative to unacclimated highlanders (88.1% vs. 78.6%). Acclimated highland mice mitigated the decline in *S*_a_O_2_ much more effectively than did acclimated lowlanders, as *S*_a_O_2_ values at 6000 m were 88.1% vs. 80.6%, respectively.

**Figure 3.**
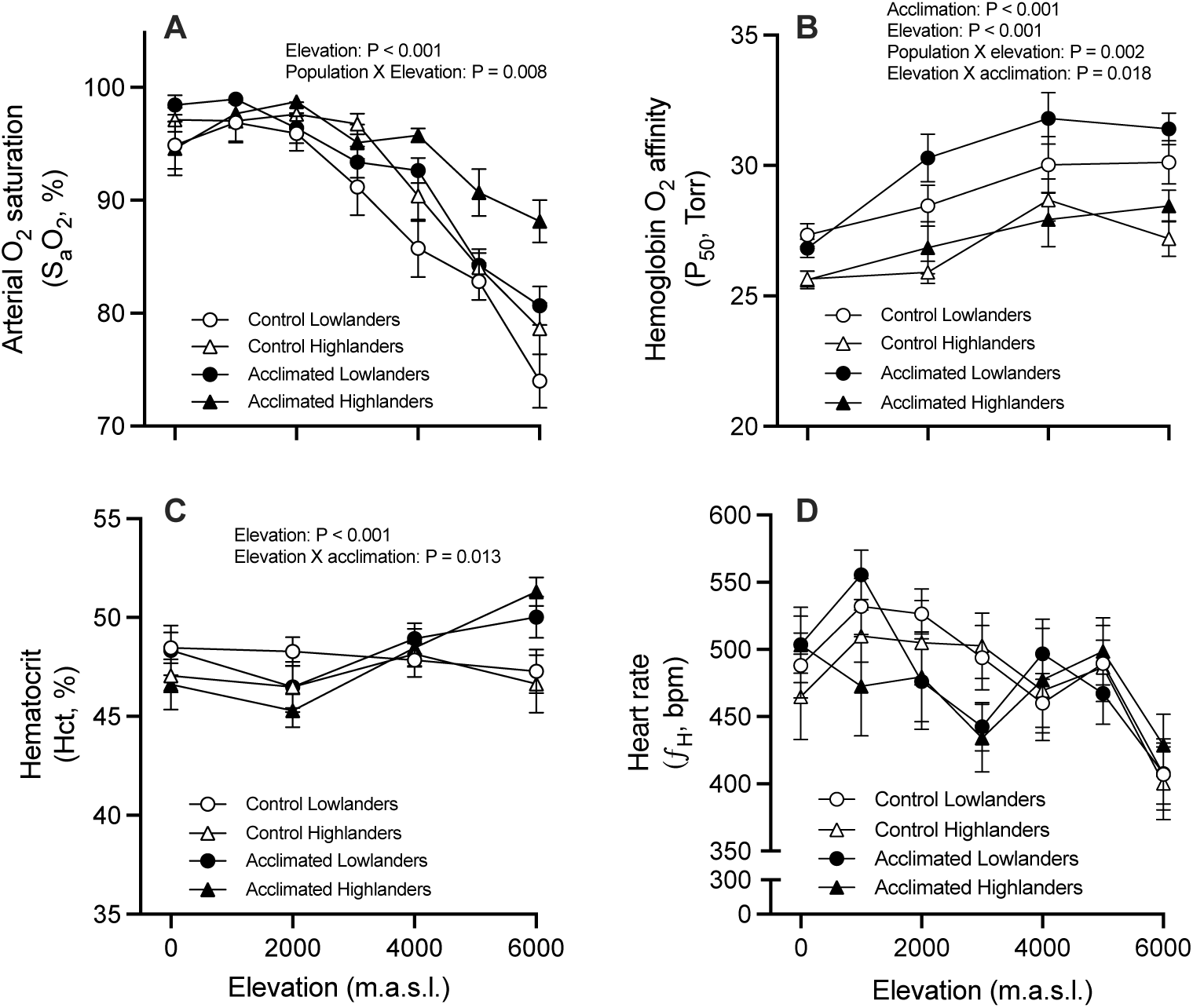
Circulatory variables. Variation in circulatory traits (measured at *V̇*O_2max_) in acclimated and control mice. **A)** Arterial O_2_ saturation and **C)** heart rate (beats per minute) estimated at the moment that the mice reached *V̇*O_2max_ during experimentation. Acclimated highlander mice exhibited higher S_a_O_2_ values at the highest elevations in comparison with the rest of the groups. **B)** Hemoglobin-O_2_ affinity (measured in intact erythrocytes at pH 7.4) and **D)** hematocrit. Statistical significance was considered with an *α* ≤ 0.05. N = 14-16 mice per data point, with exception of N = 7-11 mice per data points at 0m in B.

At a given elevation, the higher *S*_a_O_2_ of highlanders relative to lowlanders, and of acclimated mice relative to non-acclimated mice, may reflect an augmentation in blood *P*O_2_ that stems from increased ventilation (leading to increased alveolar *P*O_2_, as described above), increased pulmonary O_2_ diffusing capacity, and/or increased hemoglobin (Hb)-O_2_ affinity (by increasing Hb-O_2_ saturation at a given blood *P*O_2_ in pulmonary capillaries). Population differences in Hb-O_2_ affinity are indicated by the fact that highlanders maintained consistently lower values of *P*_50_ (the *P*O_2_ at which Hb is 50% saturated with O_2_) at each acclimation step (**Fig. 3B**). Under severe hypoxia, an increased Hb-O_2_ affinity can help safeguard *S*_a_O_2_, but this only confers a physiological benefit if tissue O_2_ diffusion capacity is sufficiently high that the augmented arterial O_2_ content translates into a greater tissue O_2_ extraction (Storz & Bautista, 2022; Wearing et al., 2021; Wearing & Scott, 2021). High-altitude vertebrates often have higher Hb-O_2_ affinities in comparison with lowland relatives (Storz, 2016, 2019), and this is also true in deer mice, as highland natives from the Rockies have evolved a higher Hb-O_2_ affinity than lowland conspecifics from the Great Plains (Jensen et al., 2016; Natarajan et al., 2015; Storz, Runck, et al., 2010; Storz et al., 2009). Moreover, genetic differences in Hb-O_2_ affinity between highland and lowland deer mice translate into consistent differences in *S*_a_O_2_ under hypoxia (Tate et al., 2017; Tate et al., 2020; Wearing et al., 2021). The higher *S*_a_O_2_ of highland mice should contribute to an increase in O_2_ transport to systemic tissues, given that highland mice have evolved higher capacities for tissue O_2_ diffusion and utilization owing to increased muscle capillary density, increased volume density of total and subsarcolemmal mitochondria, increased density of oxidative fibers, and increased mitochondrial oxidative capacity (Dawson & Scott, 2022; Lui et al., 2015; Mahalingam et al., 2020; Mahalingam et al., 2017; Scott et al., 2015; Tate et al., 2017; Tate et al., 2020). Hypoxia-induced increases in blood *P*_50_ in both highland and lowland mice (**Fig. 3B**) is almost certainly attributable to increases in the intraerythrocytic concentration of 2,3-diphosphoglycerate (DPG), an organic phosphate that is a metabolite of glycolysis, and which allosterically inhibits Hb-O_2_ binding (Storz, 2019). Upon exposure to hypoxia, increased ventilation produces a decrease in blood *P*CO_2_ (respiratory hypocapnia), which in turn leads to an increase in the intraerythrocytic concentration of DPG, thereby reducing Hb-O_2_ affinity. Thus, the effect of acclimation on *S*_a_O_2_, which is particularly strong in highlanders, cannot be explained by plasticity in Hb-O_2_ affinity, since the hypoxia-induced change in red cell DPG is in the wrong direction. Instead, the ability of acclimated highlanders to more effectively mitigate the decline in *S*_a_O_2_ at the highest elevations likely reflects hypoxia-induced improvements in pulmonary function.

In addition to minimizing the hypoxia-induced reduction in *S*_a_O_2_, other compensatory changes in convective delivery of O_2_ to metabolizing tissues are critical for sustaining aerobic performance at high elevation (Ivy & Scott, 2015; Storz & Scott, 2019). As expected, both highland and lowland mice increased hematocrit in response to hypoxia exposure (**Fig. 3C**). By the time the acclimated mice ‘ascended’ to 6000 m at the end of 7 weeks, the observed increases in hematocrit relative to unacclimated control mice is expected to reflect a hypoxia-induced increase in erythropoiesis, mediated by renal release of erythropoietin (Soliz et al., 2005). For both highlanders and lowlanders, the hematocrits of acclimated mice at 6000 m were 51±0.7% and 50±1.0%, respectively, well below the level at which increased vascular resistance would be expected to impair O_2_ transport and limit *V̇*O_2_max (Çınar et al., 1999; Hoffman, 2011; Schuler et al., 2010). For a given blood volume, an increased hematocrit increases blood O_2_ content and therefore likely benefits tissue O_2_ delivery over the range of increase observed during the 8-wk acclimation to progressively increasing hypoxia.

At any given *P*O_2_, systemic O_2_ delivery can be enhanced by increasing cardiac output (heart rate × stroke volume). Thus, if an increased cardiac output contributes to the superior thermogenic capacity of highland mice, it is likely attributable to an increased stroke volume, as previously observed (Tate et al., 2020), since heart rate at *V̇*O_2max_ exhibited no significant differences between highlanders and lowlanders in the acclimation or control groups (**Fig. 3D**).

During the 7-wk acclimation period, there were no significant changes in organ or tissue masses (**Fig. S1; Table S2**) except that lowland mice exhibited significant increases in the wet mass (but not volume) of the lungs and in the mass of the right ventricle of the heart (**Fig. S1B-C**). Hypoxia exposure can lead to pulmonary vasoconstriction and vascular remodeling, resulting in a thickening of arterial smooth muscle (Sylvester et al., 2012; West et al., 2021a). These changes reduce vessel distensibility, increase pulmonary arterial blood pressure, and can lead to pulmonary edema (West et al., 2021a; Young et al., 2019). Indeed, pulmonary edema may explain why wet lung mass increased in acclimated lowlanders without any corresponding increase in lung volume. Furthermore, pulmonary hypertension can increase the workload of the right ventricle and induce hypertrophic growth (West et al., 2021a), as observed in acclimated lowlanders. The fact that right ventricle hypertrophy and pulmonary edema did not seem to occur in highland mice subjected to the same hypoxia challenge supports previous findings suggesting that they have evolved a means of attenuating hypoxic pulmonary hypertension (West et al., 2021a). The suppression of right ventricle hypertrophy in highlander *P. maniculatus* has been associated with shifts in the expression of genes associated with the interferon regulatory factor (*IRF*) signaling pathway (Velotta et al., 2018), which plays a role in cell growth, cell differentiation and immunity, vascular hyperplasia, stroke, and cardiac hypertrophy (Jiang et al., 2014; Zhang et al., 2015).

Skeletal muscles support ventilation and contribute to thermogenesis via shivering, so changes in the oxidative capacity of skeletal muscles could contribute to variation in whole-animal thermogenic performance in multiple ways (Lui et al., 2015; Scott et al., 2015; Storz & Scott, 2019). To investigate possible changes in muscle metabolism during hypoxia acclimation we tested for changes in the activity of key enzymes that serve as markers of metabolic/oxidative capacity (HOAD, COX, CS, LDH) in the left and right ventricles, gastrocnemius, and the diaphragm of mice (**Fig. S2**; **Table S3)**. In highland mice, COX activity in the diaphragm was significantly higher than that of lowlanders irrespective of acclimation history (**Fig. S2H**), consistent with results of previous studies (Dawson et al., 2018; Lui et al., 2015). COX is the site of mitochondrial O_2_ consumption (Timón-Gómez et al., 2019), so an increased activity could facilitate greater capacity for oxidative phosphorylation and/or augment aerobic ATP production at low *P*O_2_ (Dawson & Scott, 2022; Gnaiger et al., 1998), and could thus contribute to increasing ventilation. There was no other significant variation in tissue-specific enzyme activities among groups (**Fig. S2H**), but the pattern of variation was consistent with previous findings showing that highlanders also have greater oxidative capacity in the gastrocnemius and in several other skeletal muscles (Garrett et al., 2024; Lui et al., 2015; Mahalingam et al., 2017).

In summary, genetic variation in Hb-O_2_ affinity and breathing at least partly accounts for the observed population differences in *S*_a_O_2_ in hypoxia (**Fig. 3A**), which in turn may contribute to observed differences in thermogenic *V̇*O_2max_ (**Fig. 1B**). In addition to the combination of plastic and evolved changes that enhance circulatory O_2_ transport in highlanders relative to lowlanders, lowland mice suffer characteristic symptoms of hypoxic pulmonary hypertension (right-ventricle hypertrophy) that are expected to reduce aerobic performance by impairing pulmonary function, whereas highland mice appear to have evolved a means of suppressing these maladaptive side-effects of hypoxia acclimation.

### Transcriptomic changes underlying hypoxia-induced physiological plasticity

We next used high-throughput RNA sequencing of lung and right ventricle tissues to examine how phenotypic plasticity and genotypic specialization in gene expression contribute to the evolved differences in physiological plasticity in highland mice. We sampled a total of 60 individuals across both populations (n= 30 lowland, 30 highland) at the end of the 7-week stepped acclimation experiment. Information on sequence quality metrics, as well as mapping and assignment rates are presented in **Table S4**. We used weighted gene co-expression network analysis (WGCNA – Langfleder and Horvath 2008) to identify groups of co-expressed genes within each tissue type. The WGNCA analyses revealed that both the lung and right ventricle transcriptomes were highly structured, with most genes grouping into co-expression modules. For the lung, we measured the expression of 17,552 genes, 88% of which were grouped into one of 39 co-expression modules that ranged in size from 42 to 2,193 genes (**Table S5)**. The right ventricle transcriptome was similarly structured: 82% of the 18,269 measured genes were grouped into one of 37 modules that ranged in size from 49 to 2,001 genes (**Table S5**).

Each of these co-expression modules comprise genes that exhibit correlated expression patterns across samples. As a result, module expression can be summarized using principal components analysis (PCA), with the first principal component (PC1) representing overall module expression (hereafter known as the gene eigenvalue). We tested for effects of population (highland vs. lowland), acclimation treatment (hypoxia vs. normoxia), and their interaction on module expression using an analysis-of-variance (ANOVA) on rank-transformed module eigengene values following Plachetzki et al. (2014). In the lung, 14 modules (36%) exhibited significant population effects, indicating constitutive differences in module expression between highlanders and lowlanders (**Table 1**). This may partly reflect population differences in cell-type composition arising from differences in lung structure (West et al., 2021b). Treatment effects that are independent of population indicate transcriptional responses to hypoxia that are shared between highlanders and lowlanders; six modules exhibited this pattern in the lung transcriptome (**Table 1; Table S6**). Population differences in the transcriptional response to hypoxia would be indicated by significant interactions between population and treatment. Here, only a single module (L26) exhibited this pattern in the lung transcriptome (**Table 1; Table S6**). We recovered qualitatively similar patterns in the right ventricle transcriptome, where the 10 modules (27%) exhibited population effects on expression, and only a single module exhibited treatment effects and an interaction between population and treatment (**Table 1; Table S10**). Overall, this analysis revealed that population of origin had much stronger effects on module expression than acclimation treatment, suggesting that genotypic specialization may play an important role in shaping the transcriptome of high-altitude mice.

**Table 1.** Module expression. Population and treatment effects on module expression for the lung and right ventricle. A significant population effect is indicative of constitutive differences in module expression between highlanders and lowlanders while treatment effects that are independent of population indicate transcriptional responses to hypoxia that are shared between highlanders and lowlanders. Population differences in the transcriptional response to hypoxia would be indicated by significant interactions between population and treatment.

| Tissue | Population | Treatment | Population*Treatment |
| --- | --- | --- | --- |
| Lung | 14 (36%) | 6 (15%) | 1 (3%) |
| Right Ventricle | 10 (27%) | 1 (3%) | 1 (3%) |

To gain further insight into the mechanistic basis of performance differences between highlanders and lowlanders, we tested whether transcriptional module expression was correlated with variation in two key subordinate traits that exhibited population-specific hypoxia responses: arterial oxygen (*S*_a_O_2_), because variation in this trait may be due in part to variation in pulmonary structure and function, and relative right ventricle mass. Both traits exhibited distinct responsiveness to hypoxia in lowlanders and highlanders (**Fig. 3 and Fig. S2B**). To test whether changes in the expression of transcriptional modules contribute to population-specific physiological responses in pulmonary function and right ventricle hypertrophy, we tested for associations of *S*_a_O_2_ values with module expression in the lung transcriptome, and for associations of relative right ventricle masses with module expression in the right ventricle transcriptome. We identified one module in the lung transcriptome that was negatively associated with *S*_a_O_2_ (L38 *r* = -0.59, FDR adjusted *p* = 0.0023, **Table S7**), but was not differentially expressed between highlanders and lowlanders. Module L38 is a small module comprised of 44 genes and was enriched for genes involved in the assembly and arrangement of constituent parts of the extracellular matrix and granulocytosis. The hub gene associated with this module is Basonuclin Zinc Finger Protein, Bnc1 (**Table S7**), a gene that encodes a zinc finger protein found in the basal cell layer of the epidermis and in hair follicles and is thought to play a regulatory role in keratinocyte proliferation and rRNA transcription as a regulator.

None of the modules in the right ventricle transcriptome were associated with relative right ventricle mass after multiple test correction (**Table S10**). However, the modules with the strongest associations (RV03 and RV32) had opposing associations with right ventricle mass. Module RV03 was negatively associated with mass (r = -0.363, nominal p-value = 0.018, FDR corrected *p* = 0.2265, **Table S11**) (**Fig. 4C**), and was enriched for genes involved in lipid and other small molecule metabolic processes (GO:0006629, GO:0044281, **Table S12; Fig. 4**). Expression of module RV03 was not responsive to hypoxia, but was constitutively expressed at a significantly higher level in highlanders relative to lowlanders (**Fig. 4D**). Module RV32 was positively associated with mass (r = 0.36, nominal p-value = 0.019, FDR corrected *p* = 0.2265: **Table S11**; **Fig. 4E**), and was enriched for genes involved in transmembrane tyrosine kinase signaling (GO:0007169, **Table S12; Fig 4**). This module was upregulated in response to hypoxia in lowlanders, but was not responsive to hypoxia and was constitutively expressed at a lower level in highlanders (**Fig. 4F**). The regulation of tyrosine kinases has been implicated in cardiac hypertrophy, and several cardiac myopathies in mammals, including rats and house mice (Sala et al., 2012).

**Figure 4.**
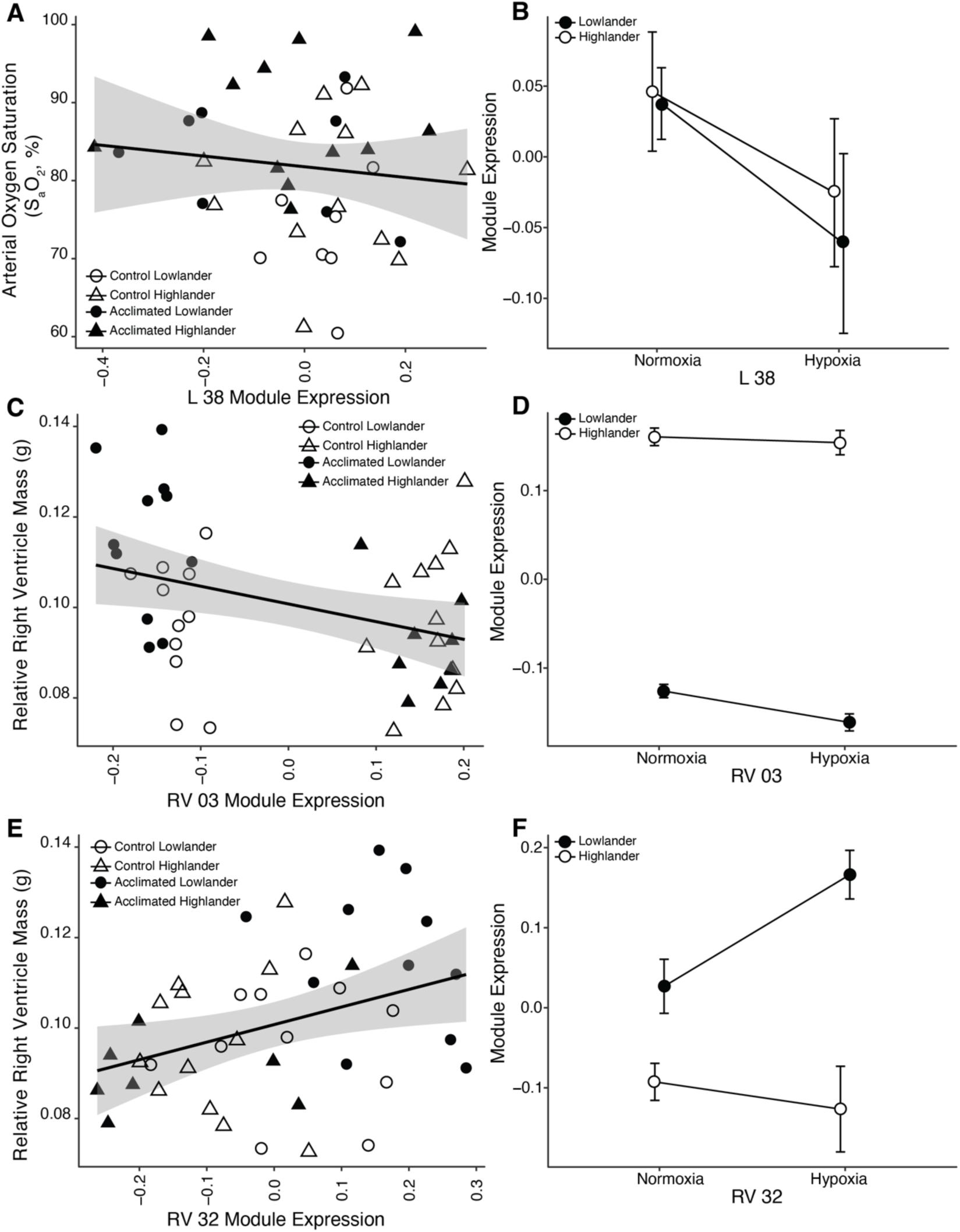
Module expression. Lung and right ventricle module expression and trait associations **A)** L 38 expression relative to S_a_O_2_ **B)** module L 38 expression **C)** RV 03 expression and relative right ventricle mass **D)** RV 03 expression **E)** RV 32 expression relative to right ventricle mass **F)** RV 32 expression.

Previous acclimation experiments have demonstrated that the suppression of hypoxia-induced right ventricle hypertrophy in highland deer mice is associated with differential regulation of interferon regulatory factors (*IRF1*, *IRF7*, and *IRF9*) and other inflammatory signaling genes (Velotta et al. 2018). These experiments compared highland deer mice to a closely-related lowland congener, *P. leucopus*, at a simulated elevation of 4350 m, whereas our experiments compared conspecific populations of deer mice acclimated to a much more extreme simulated elevation (6000 m). Velotta et al. (2018) identified two regulatory modules associated with right ventricle mass, and we tested for overlap in gene content between those two modules and the modules that exhibited the strongest associations with right ventricle mass in the present study (RV03 and RV32). Of the 1035 genes that comprise modules RV03 and RV32, 20 overlapped with genes in the modules identified by Velotta et al. (2018) (**Table S13**), representing a significant enrichment (Fisher’s Exact Test p = 3.45 x 10^-5^). All 20 of the overlapping genes were members of module RV03 and included interferon regulatory factor 7 (*IRF7*). Velotta et al. (2018) had identified *IRF7* as a hub gene in one of the regulatory modules associated with right ventricle hypertrophy, and highland deer mice exhibited constitutively elevated expression of this gene relative to the lowland congener, *P. leucopus*. We also identified *IRF7* as a hub gene in module RV03 (intermodular connectivity, KME score = 0.379, top 10% of kME scores for RV03), and this module was constitutively upregulated in highland mice (**Fig. 4D**). *IRF7* is a negative regulator of pathological cardiac hypertrophy in rodents (Jiang et al., 2014), suggesting that the upregulation of this gene (and others in module RV03) may contribute to the suppression of right ventricle hypertrophy in hypoxia-acclimated highland deer mice. The lack of more extensive overlap between the RV03/RV32 gene sets and the gene sets that were associated with right ventricle hypertrophy in Velotta et al. (2018) (**Table S13**) may be explained by differences between studies in the evolutionary scale of comparison (between species versus between conspecific populations) and/or differences in the severity hypoxia exposure (4350 m vs 6000 m). Nonetheless, the overlapping gene set (**Table S13**) provides a list of promising candidates for follow-up investigations into evolved mechanisms of suppressing right ventricle hypertrophy in hypoxia.

## CONCLUSIONS

At levels of hypoxia corresponding to the upper elevational range limits of *P. maniculatus* (and beyond), the superior thermogenic capacity of highland natives relative to lowland conspecifics indicates that the ability of the species to inhabit such a broad range of elevations is attributable to a combination of genetically based local adaptation and evolved changes in the plastic response to hypoxia. At simulated elevations >4000 m, highland natives mitigated the decline in arterial O_2_ saturation more effectively than did lowlanders due to their higher Hb-O_2_ affinity and a more responsive, hypoxia-induced increase in breathing rate. Whereas hypoxia-acclimated lowlanders developed right-ventricle hypertrophy, a symptom of hypoxic pulmonary hypertension that is expected to impair aerobic performance, highlanders appear to have evolved a means of suppressing this maladaptive side-effect of hypoxia acclimation. This population difference in physiological plasticity is associated with regulatory differences in hypoxia-induced gene expression in the right-ventricle. Our results reveal the importance of local adaptation, including evolved changes in plasticity and the suppression of maladaptive plasticity, in determining the niche breadth of species that are distributed across environmentally heterogeneous landscapes.

## MATERIALS AND METHODS

### Mouse populations and Acclimation protocol

Wild adult deer mice (*Peromyscus maniculatus rufinus* [Wagner 1985]) native to lowland (∼500 m.a.s.l., Kearney, NE, USA) and highland (Mount Blue Sky ∼4350 m.a.s.l., CO, USA) environments were live-trapped and transported to McMaster University (Hamilton, ON Canada) where captive breeding populations were established. Mice used in this study were second-generation progeny from each population that were raised and maintained in common husbandry conditions (normoxia 24–25°C, 12h:12h light:dark photoperiod) and provided with water and standard rodent chow *ad libitum*. Experimental protocols were conducted in two separated replicates with a total of 32 adult mice each (∼6 months old). The 32 mice from the first replicate belonged to five families, and the 32 mice of the second replicate belonged to seven families. Each replicate consisted of 16 highlander and 16 lowlander mice with equal numbers of females and males. Each population was divided into control and acclimation treatments. Control mice were maintained in normobaric normoxia (21% O_2_) at 25°C, 12h:12h light:dark photoperiod with *ad libitum* access to food and water throughout the length of the study. In contrast, the acclimation treatment involved exposing the mice to discrete, stepwise reductions in the partial pressure of O_2_ (*P*O_2_), simulating a mountain ascent from sea level to 6000 m.a.s.l. at increments of 1000 m. Incremental increases in isobaric hypoxia corresponded to 21%, 18.7%, 16.8%, 14.8%, 13.1%, 11.6% and 10.1% O_2_. Exposure to each *P*O_2_ occurred for 8 days and was conducted inside a glove box chamber (O_2_ Control In Vitro Glove Box. Coy Laboratory Products Inc., MI, USA.) where the temperature, photoperiod and access to food and water were identical to the control conditions. Cleaning of the holding cages was performed twice a week, which required that the ascending hypoxic group be returned to normoxia for a brief period (<15min) per cleaning event. Body mass, food and water consumption were recorded **(Fig. S3; Table S1).** All experimental protocols were approved by the Animal Research Ethics Board at McMaster University (AUP 20-01-02) following guidelines established by the Canadian Council on Animal Care.

### Measurement of thermogenic *V̇*O_2max_ and subordinate cardiorespiratory traits

Measurements of thermogenic *V̇*O_2max_ were recorded in the acclimation group after 6 days at each acclimation step (simulating ascending elevations of 0m, 1000m, 2000m, 3000m, 4000m, 5000m, and 6000m) at the PO_2_ of that step. Time-matched measurements were made in the control group at the same PO_2_. We used open-flow respirometry following established protocols that have been shown to elicit values of thermogenic *V̇*O_2max_ that equal or exceed those measured during exercise (Chappell & Hammond, 2004; Ivy et al., 2021; Wearing et al., 2021). In brief, we determined thermogenic *V̇*O_2max_ as the maximal rate of O_2_ consumption measured over 30s during acute exposure to cold heliox. *V̇*O_2_ measurements were performed on each individual mouse placed inside a 530mL whole-body plethysmography chamber. The chamber was kept inside a freezer to maintain its internal temperature at -5°C, confirmed with a thermocouple (PT-6, Physitemp, Clifton, NJ USA). The proportion of O_2_ (balance He) in the incurrent flow to the chamber (1500 ml * min^-1^) was regulated and measured with an MFC-2 mass flow controller (Sable Systems, Las Vegas, NV, USA) and precision flow control valves for O_2_ and helium (Sierra Instruments, Monterey CA, USA). The gas was cooled prior to entering the chamber by passing the flow through copper coils inside the freezer. A differential pressure transducer (Validyne DP45: Cancoppas, Mississauga, ON, Canada) was used for measuring pressure oscillations induced by the breathing of the mouse relative to a reference chamber. The differential pressure transducer was calibrated to volume prior to each individual *V̇*O_2max_ measurement by repeatedly withdrawing and injecting 300 μL from the chamber using a Hamilton syringe, after which baseline O_2_ and CO_2_ fractions of incurrent gas passing through the empty respirometry chamber were recorded. The individual mouse was then weighed in a top loading balance, its core body temperature (*T*_b_) was determined with a rectal probe (RET-3-ISO; Physitemp, Clifton, NJ, USA), and it was fitted with a MouseOx Plus pulse oximetry neck collar (Starr Life Sciences, Oakmont, PA, USA; which required prior removal of the fur of the neck). The mouse was then placed inside the chamber (with the neck collar connected through a port in the chamber lid) for simultaneous measurements of excurrent O_2_ and CO_2_ fractions, breathing, heart rate (*f*_H_), and arterial O_2_ saturation (S_a_O_2_) during exposure to cold (-5°C) heliox for ∼10 min. O_2_ and CO_2_ fractions were measured by passing a subsample of the gas stream through pre-baked drierite followed by a fuel cell O_2_ analyzer (FC-10; Sable Systems) and infrared CO_2_ analyzer (CA-10; Sable Systems). Measurements of incurrent gas flow rate, O_2_ and CO_2_ fractions, and chamber temperature were recorded using PowerLab 8/32 and LabChart 8 Pro software (ADinstruments, Colorado Springs, CO, USA). Pulse oximetry data was recorded using the Starr Life Sciences acquisition software. *V̇*O_2max_ was reached between 4-8 min of the trial and was calculated based on incurrent and excurrent O_2_ and CO_2_ gas fractions with established respirometry formulas (Lighton, 2018). Tidal volume (*V*_T_) was calculated based on the barometric method of plethysmography in flow-through respirometry conditions (Drorbaug & Fenn, 1955; Jacky, 1980; Lim et al., 2014). Total ventilation (*V̇*_E_) is the product of breathing frequency (*ƒ*_R_) and V*_T_*, air convection requirement for O_2_ is *V̇*_E_/*V̇*O_2max_, air convection requirement for CO_2_ is *V̇*_E_/*V̇*CO_2_ (measured concurrent with the measurement of *V̇*O_2max_), pulmonary O_2_ extraction was calculated as *V̇*O_2max_ divided by the product of *V̇*_E_ and [O_2_] of inspired air, and respiratory exchange ratio was calculated as *V̇*CO_2_/*V̇*O_2max_.

### Measurement of hematological traits

Hematological variables were determined two days after *V̇*O_2max_ was measured at PO_2_s corresponding to elevations of 0m, 2000m, 4000m and 6000m. Thirty microliters of blood were sampled by carefully pinching the ventral artery of the tail of individual mouse under anesthesia (1.5% isoflurane), and used for estimating hematological variables as follows: 10μL were used for calculating hemoglobin concentration ([Hb]) using Drabkin’s solution (Briggs & Bain, 2017); 10μL were sampled with two 5μL capillary tubes (Drummond Microcaps®) for determining hematocrit (Hct) by spinning the blood for 5min at 12,000g; and the last 10μL of blood were added to 5 ml of buffer (50 mM HEPES, 10 mM EDTA, 100 mM NaCl, 0.1% bovine serum albumin, 0.2% antifoaming agent, at pH of 7.4 at 37°C) and used to determine the O_2_ affinity (P_50_) and cooperativity (n_50_) of intact erythrocytes from O_2_ dissociation curves generated with a Hemox Analyzer and its Analytic Software (TCS Scientific).

### Organ mass and lung volume

Mice were euthanized (isoflurane overdose followed by cervical dislocation) after the last blood sampling event and the gastrocnemius, soleus, and diaphragm muscles, as well as the liver, kidney, spleen, heart, and interscapular brown fat were carefully excised, weighed, and immediately flash-frozen in liquid N_2_ and stored at -80°C. The trachea was then cannulated with PE50 tubing (FisherScientific, Mississauga, Canada) and secured with 2-0 gauge suture silk (Prolene, Fisher Scientific). The trachea and lungs were carefully excised, inflated with saline solution (0.9% NaCl) to a pressure of 30 cm H_2_O, and lung volume was then determined by the immersion displacement technique (Scherle, 1970). The lungs were then deflated and flash-frozen in liquid N_2_ and stored at -80°C.

### Enzyme activity

The right ventricle, combined left ventricle and septum, and gastrocnemius were powdered using a liquid N_2_-cooled mortar and pestle, and stored at -80°C. Powdered samples of ∼10mg were homogenized in 20 volumes of homogenization buffer (concentrations in mM: 100 KH_2_PO_4_, 5 ethylenediaminetetraacetic acid, and 0.1% Triton-X-100, at pH 7.2) using a glass tissue homogenizer. The maximal activities of cytochrome-c oxidase (COX) and ß-hydroxyacyl CoA dehydrogenase (HOAD) were assayed shortly after homogenization, and the maximal activities of citrate synthase (CS) and lactate dehydrogenase (LDH) were measured after storage of homogenate at -80°C. Activity was assayed at 37°C by measuring the change in absorbance over time (HOAD, 340nm; COX: 550nm; CS, 412nm; LDH, 340nm) under the following conditions: HOAD, 0.1 mM acetoacetyl-CoA (omitted in background reactions), 0.28 mM NADH, 100 mM triethylamine hydrochloride, pH 7.0; COX, 100 mM KH_2_PO_4_, 0.1 mM reduced cytochrome C (omitted in background reactions), pH 7.0; CS, 40 mM Tris, 0.5 mM oxaloacetate (omitted in background reactions), 0.22 mM acetyl-CoA, 0.1 mM 5,5’-dithio-bis-(2-nitrobenzoic acid), pH 8.0; LDH, 40 mM Tris, 0.28 mM NADH, 1 mM pyruvate (omitted in background reactions), pH 7.4. Preliminary experiments verified that substrate concentrations were saturating. All enzyme assays were run in triplicate. Enzyme activities were calculated as the reaction rates measured in the presence of all substrates minus the background rate (measured in background reactions that lacked the key substrates identified above).

### Sample preparation for RNA-seq

We used high-throughput RNA sequencing to identify gene expression differences among individual deer mice from lowland and highland populations. Libraries were prepared and sequenced by Azenta Life Technologies. Briefly, for each sample, RNA was extracted from the lung and right ventricle of the heart using a SMART-Seq HT kit for full-length cDNA synthesis and amplification. Sequencing libraries were prepared using an Illumina Nextera X kit, quality checked using a TapeStation, and then were sequenced on a shared lane of Illumina HiSeq 4000 (PE 150 bp) platform by Azenta.

### Statistical Analysis: Physiological variables

Statistical analysis and graphing were performed in R V4.4.0 (https://cran.r-project.org/), R-studio V2024.04.0+735 (https://www.rstudio.com/), and Prism V9 (GraphPad). Normality of the residuals of the statistical linear models data were analyzed visually and with Shapiro-Wilk test; homogeneity of the variance was assessed with Levene’s test (Quinn & Keough, 2002). If data did not meet the assumptions, rank transformation was performed before re-analyzing data. Post hoc analyses were performed with Tukey’s honest significant test (TukeyHSD) to determine differences among group means. Statistical linear mixed effects models were computed using the *lmer* function from the *lme4* package in R, and the best fitted model was chosen based on the lowest AIC value. Statistically significant effects of interest are shown within the figures, and the complete result from statistical analysis are given in supplemental information **(Tables S1-3**).

Body mass, *V̇*O_2max_, S_a_O_2_, *f_H_*, *V*_T_, *V̇*_E_, ACR-O_2_, ACR-CO_2_, *V̇*CO_2_, RER, *V̇*_E_/*V̇*CO_2_, Pulmonary O_2_ extraction, Hct, [Hb], P_50_, N_50_, were compared between groups considering population, elevation, acclimation, sex, and replicate as fixed effects, the family to which the mice belong was considered as a random effect, the ID number of each mouse was included as a random effect (nested within family) to account for repeated measurements, and body mass was included as a covariate in the models (with the exception of body mass itself). Replicate number was not computed in the analysis of water and food consumption because these data were only collected for replicate 2. Organ and tissue masses were compared among groups considering population, acclimation, sex, and replicate as fixed effects, the family of the mice was considered as random effect, and body mass was included as a covariate. Enzyme activities were compared among groups with a similar model as organ masses, but replicate number was not included because we only ran the assays on tissues of mice from Replicate 1.

### Analysis of transcriptomes

RNA-seq raw reads for each tissue were treated separately and were cleaned using the FastP pipeline (Chen et al., 2018) to trim adapters and low-quality bases. Quality metrics were assessed using multiQC v 1.11 (Ewels et al., 2016). Cleaned reads were then mapped to the chromosome-level assembly of the nuclear and mitochondrial *Peromyscus maniculatus* genomes (GCF_003704035.1) using HiSat2 2.2.1 (Zhang et al., 2021) with a maximum and minimum mismatch penalty (−mp 2.0). We implemented featureCounts v2.0.3 (Liao et al., 2014) to count numbers of reads aligning to annotated genes (i.e., counts were performed at the gene level; n = 32,828); featureCounts was run using default parameters, with the exception that we counted reads overlapping with more than one feature (-O flag), meaning that each overlapping feature receives a count of 1 from a read.

Count data were processed and analyzed using the R v 4.3.3 programming language (https://www.R-project.org) and all of the pipeline scripts/commands and output files are available via GitHub (https://github.com/NathanaeldHerrera/Pman_rnaseq). For each tissue, we first assessed the count data for potential technical artifacts due to sequencing using a principal components analysis (PCA) with the base prcomp function using five technical sequencing metrics: mean quality score, percent bases with quality score greater than or equal to 30, percent GC content, genome alignment rate, percent reads assigned with featureCounts. PC1 and PC2 were plotted using ggbiplot v 0.6.2 (https://github.com/friendly/ggbiplot) to identify samples located outside of the 95% data ellipse (**Fig. S4**). For the right ventricle, we removed two samples and for the lung, we removed five samples. We filtered read counts with less than an average of 20 reads per individual since genes with low read counts are subject to measurement error (Robinson & Smyth, 2007). We retained a total of 17,752 and 14,890 genes after filtering for lung and right ventricle transcriptomes, respectively. Three outlier samples (three lung, one right ventricle) were removed following visual inspection of multidimensional scaling plots using plotMDS in edgeR (Robinson et al., 2010) following Chen et al. (2019). Raw read counts were then normalized by library size and log-transformed using the calcNormFactors and cpm functions within the edgeR package.

We used weighted gene co-expression network analysis (WGCNA) (Langfelder & Horvath, 2008) to identify modules (sets of genes) that are co-expressed across each tissue type, thereby providing insights into how regulatory mechanisms are affected by treatments. Network construction and module detection was performed using the blockwiseModules function in WGCNA with networkType set to “signed”. We used a maximum block size of 18,000 (greater than the total number of genes) and setting the random seed for reproducibility. Co-expression networks were constructed by assessing pairwise Pearson correlations among gene pairs. Subsequently, an adjacency matrix was computed by applying a soft thresholding power of b = 8 and b = 12 to the correlation matrix for the lung and right ventricle, respectively. The choice of soft thresholding power, b, aims to establish a network with an approximately scale-free topology, favoring strong correlations over weak ones (Zhang & Hovarth, 2005). The value of b for each tissue dataset was selected because it marks the point at which the improvement of scale-free topology model fit begins to decline with increasing thresholding power (i.e., the inflection point; **Fig. S5**). From the resulting adjacency matrix, a topological overlap measure (a robust indicator of interconnectedness) was calculated for each gene pair. Topological based dissimilarity was then calculated and used as input for average linkage hierarchical clustering to generate cluster dendrograms. Modules, representing clusters of genes with dense interconnections and high correlation, were identified as branches of the resulting cluster tree using the dynamic tree-cutting method (Langfelder & Horvath, 2008).

After defining modules, we used a multistep approach to link individual expression with variation in SaO_2_ for the lung and relative right ventricle mass for the right ventricle. First, we summarized module expression using PCA of gene expression profiles using the blockwiseModules function in WGCNA. Given that genes within modules are inherently correlated, we utilized the first principal component axis, also known as the module eigengene, to represent module expression (Langfelder and Horvath, 2008). We then used module eigengene values to test for associations between module expression and each phenotype within each module (using Pearson correlation; cor function in WGCNA) across each network. P-values for these correlations were computed using Student’s asymptotic test (corPvalueStudent function in WGCNA). We also conducted an analysis-of-variance (ANOVA) on rank-transformed module eigengene values in order to test for the effects of population, treatment, and their interaction on module expression following Plachetzki et al. (2014). P-values from association tests and ANOVAs were corrected for multiple testing using the false-discovery rate method (Benjamini & Hochberg, 1995). For all modules exhibiting a statistically significant effect in the ANOVA, we used the R package gProfilerR (Reimand et al., 2016) to perform a functional enrichment analysis (**Fig. S6; Table S9, S12**).

## Statements

### Ethical statement

All protocols of animal handling and sampling were performed in accordance with the Canadian Council on Animal Care and were approved by the McMaster University Animal Research Ethics Board (AUP #20-01-02).

### Competing Interest Statement

None

### Funding

This work was funded by grants from the National Institutes of Health (R01 HL159061, JFS and ZAC), National Science Foundation (IOS-2114465, JFS and ZAC; OIA-1736249, JFS and ZAC), and the National Geographic Society (NGS-68495R-20, JFS).

## Acknowledgments

We thank Folasade Ologundudu for technical assistance during the development of the experimental design.

## Notes

**Competing Interest Statement:** The authors declare no conflict of interest.

### Competing Interest Statement

The authors have declared no competing interest.

## REFERENCES

Benjamini, Y., & Hochberg, Y. (1995). Contrilling the false discovery rate: a practical and powerful approach to multiple testing. J. R. Statist. Soc. B, 57(1), 289–300.

Briggs, C., & Bain, B. J. (2017). 3 - Basic Haematological Techniques. In B. J. Bain, I. Bates, & M. A. Laffan (Eds.), Dacie and Lewis Practical Haematology (Twelfth Edition) (pp. 18–49). Elsevier. 10.1016/B978-0-7020-6696-2.00003-5

Case, T. J., & Taper, M. L. (2000). Interspecific competition, environmental gradients, gene flow, and the coevolution of species’ borders. The American Naturalist, 155(5), 583–605.

Chappell, M. A., & Hammond, K. A. (2004). Maximal aerobic performance of deer mice in combined cold and exercise challenges. *Journal of Comparative Physiology B: Biochemical*, Systemic, and Environmental Physiology, 174(1), 41–48. 10.1007/s00360-003-0387-z

Chen, S., Zhou, Y., Chen, Y., & Gu, J. (2018). fastp: an ultra-fast all-in-one FASTQ preprocessor. Bioinformatics, 34(17), i884–i890. 10.1093/bioinformatics/bty560

Chen, Y., McCarthy, D., Balldoni, P., Ritchie, M., Robinson, M., & Smyth, G. (2019). edgeR: differential analysis of sequence read count data User’s Guide. *Available from:* https://bioconductor.org/packages/release/bioc/vignettes/edgeR/inst/doc/edgeRUsersGuide.pdf.

Chevin, L. M., & Lande, R. (2011). Adaptation to marginal habitats by evolution of increased phenotypic plasticity. Journal of Evolutionary Biology, 24(7), 1462–1476. 10.1111/j.1420-9101.2011.02279.x

Cheviron, Z. A., Bachman, G. C., Connaty, A. D., McClelland, G. B., & Storz, J. F. (2012). Regulatory changes contribute to the adaptive enhancement of thermogenic capacity in high-altitude deer mice. Proceedings of the National Academy of Sciences, 109(22), 8635–8640. 10.1073/pnas.1120523109

Cheviron, Z. A., Bachman, G. C., & Storz, J. F. (2013). Contributions of phenotypic plasticity to differences in thermogenic performance between highland and lowland deer mice. Journal of Experimental Biology, 216(7), 1160–1166. 10.1242/jeb.075598

Cheviron, Z. A., Connaty Ad Fau - McClelland, G. B., McClelland Gb Fau - Storz, J. F., & Storz, J. F. (2014). Functional genomics of adaptation to hypoxic cold-stress in high-altitude deer mice: transcriptomic plasticity and thermogenic performance. Evolution, 68(1558-5646 (Electronic)). 10.1111/evo.12257

Çınar, Y., G., D., M., P., & B., Ç. A. (1999). Effect of hematocrit on blood pressure via hyperviscosity. American Journal of Hypertension, 12(7), 739–743. 10.1016/s0895-7061(99)00011-4

Dawson, N. J., Lyons, S. A., Henry, D. A., & Scott, G. R. (2018). Effects of chronic hypoxia on diaphragm function in deer mice native to high altitude. Acta Physiologica, 223(1), e13030. 10.1111/apha.13030

Dawson, N. J., & Scott, G. R. (2022). Adaptive increases in respiratory capacity and O_2_ affinity of subsarcolemmal mitochondria from skeletal muscle of high-altitude deer mice. The FASEB Journal, 36(7). 10.1096/fj.202200219r

Drorbaug, J. E., & Fenn, W. O. (1955). A barometric method for measuring ventilation in newborn infants. Pediatrics, 16(1), 81–87. 10.1542/peds.16.1.81

Ewels, P., Magnusson, M., Lundin, S., & Kaller, M. (2016). MultiQC: summarize analysis results for multiple tools and samples in a single report. Bioinformatics, 32(19), 3047–3048. 10.1093/bioinformatics/btw354

Garrett, E. J., Prasad, S. K., Schweizer, R. M., McClelland, G. B., & Scott, G. R. (2024). Evolved changes in phenotype across skeletal muscles in deer mice native to high altitude. *American Journal of Physiology-Regulatory*, Integrative and Comparative Physiology, 326(4), R297–R310. 10.1152/ajpregu.00206.2023

Gnaiger, E., Lassnig, B., Kuznetsov, A., Rieger, G., & Margreiter, R. (1998). Mitochondrial Oxygen Affinity, Respiratory Flux Control And Excess Capacity Of Cytochrome c Oxidase. Journal of Experimental Biology, 201(8), 1129–1139. 10.1242/jeb.201.8.1129

Goldberg, E. E., & Price, T. D. (2022). Effects of Plasticity on Elevational Range Size and Species Richness. The American Naturalist, 200(3), 316–329.

Hayes, J. P., & O’Connor, C. S. (1999). Natural Selection on Thermogenic Capacity of High-Altitude Deer Mice. Evolution, 53(4), 1280–1287. 10.1111/j.1558-5646.1999.tb04540.x

Hoffman, J. I. E. (2011). Pulmonary Vascular Resistance and Viscosity: The Forgotten Factor. Pediatric Cardiology, 32(5), 557–561. 10.1007/s00246-011-9954-3

Holt, R. D., & Gaines, M. S. (1992). Analysis of adaptation in heterogeneous landscapes: Implications for the evolution of fundamental niches. Evolutionary ecology, 6, 433–447.

Ivy, C. M., Prest, H., West, C. M., & Scott, G. R. (2021). Distinct Mechanisms Underlie Developmental Plasticity and Adult Acclimation of Thermogenic Capacity in High-Altitude Deer Mice [Original Research]. Frontiers in Physiology, 12. https://www.frontiersin.org/article/10.3389/fphys.2021.718163

Ivy, C. M., & Scott, G. R. (2015). Control of breathing and the circulation in high-altitude mammals and birds. Comparative Biochemistry and Physiology Part A: Molecular & Integrative Physiology, 186, 66–74. 10.1016/j.cbpa.2014.10.009

Jacky, J. P. (1980). Barometric measurement of tidal volume: effects of pattern and nasal temperature. 80(0161-7567).

Jensen, B., Storz, J. F., & Fago, A. (2016). Bohr effect and temperature sensitivity of hemoglobins from highland and lowland deer mice. Comparative Biochemistry and Physiology Part A: Molecular & Integrative Physiology, 195, 10–14. 10.1016/j.cbpa.2016.01.018

Jiang, D.-S., Liu, Y., Zhou, H., Zhang, Y., Zhang, X.-D., Zhang, X.-F., Chen, K., Gao, L., Peng, J., Gong, H., Chen, Y., Yang, Q., Liu, P. P., Fan, G.-C., Zou, Y., & Li, H. (2014). Interferon Regulatory Factor 7 Functions as a Novel Negative Regulator of Pathological Cardiac Hypertrophy. Hypertension, 63(4), 713–722. 10.1161/hypertensionaha.113.02653

Kirkpatrick, M., & Barton, N. H. (1997). Evolution of a species’ range. The American Naturalist, 150(0003- 0147 (Print)), 1–23.

Langfelder, P., & Horvath, S. (2008). WGCNA: an R package for weighted correlation network analysis. BMC Bioinformatics, 9, 559. 10.1186/1471-2105-9-559

Liao, Y., Smyth, G. K., & Shi, W. (2014). featureCounts: an efficient general purpose program for assigning sequence reads to genomic features. Bioinformatics, 30(7), 923–930. 10.1093/bioinformatics/btt656

Lighton, J. R. (2018). Measuring metabolic rates: a manual for scientists. Oxford University Press.

Lim, R., Zavou, M. J., Milton, P.-L., Chan, S. T., Tan, J. L., Dickinson, H., Murphy, S. V., Jenkin, G., & Wallace, E. M. (2014). Measuring Respiratory Function in Mice Using Unrestrained Whole-body Plethysmography. Journal of Visualized Experiments(90). 10.3791/51755

Lui, M. A., Mahalingam, S., Patel, P., Connaty, A. D., Ivy, C. M., Cheviron, Z. A., Storz, J. F., McClelland, G. B., & Scott, G. R. (2015). High-altitude ancestry and hypoxia acclimation have distinct effects on exercise capacity and muscle phenotype in deer mice. *American Journal of Physiology-Regulatory*, Integrative and Comparative Physiology, 308(9), R779–R791. 10.1152/ajpregu.00362.2014

Mahalingam, S., Cheviron, Z. A., Storz, J. F., McClelland, G. B., & Scott, G. R. (2020). Chronic cold exposure induces mitochondrial plasticity in deer mice native to high altitudes. The Journal of Physiology, 598(23), 5411–5426. 10.1113/jp280298

Mahalingam, S., McClelland, G. B., & Scott, G. R. (2017). Evolved changes in the intracellular distribution and physiology of muscle mitochondria in high-altitude native deer mice. The Journal of Physiology, 595(14), 4785–4801. 10.1113/jp274130

McClelland, G. B., & Scott, G. R. (2019). Evolved Mechanisms of Aerobic Performance and Hypoxia Resistance in High-Altitude Natives. Annual Review of Physiology, 81(1), 561–583. 10.1146/annurev-physiol-021317-121527

Natarajan, C., Hoffmann, F. G., Lanier, H. C., Wolf, C. J., Cheviron, Z. A., Spangler, M. L., Weber, R. E., Fago, A., & Storz, J. F. (2015). Intraspecific Polymorphism, Interspecific Divergence, and the Origins of Function-Altering Mutations in Deer Mouse Hemoglobin. Molecular Biology and Evolution, 32(4), 978–997. 10.1093/molbev/msu403

Plachetzki, D. C., Sabrina Pankey, M., Johnson, B. R., Ronne, E. J., Kopp, A., & Grosberg, R. K. (2014). Gene co-expression modules underlying polymorphic and monomorphic zooids in the colonial hydrozoan, Hydractinia symbiolongicarpus. Integr Comp Biol, 54(2), 276–283. 10.1093/icb/icu080

Quinn, G. P., & Keough, M. J. (2002). Experimental design and data analysis for biologists. Cambridge university press.

Reimand, J., Arak, T., Adler, P., Kolberg, L., Reisberg, S., Peterson, H., & Vilo, J. (2016). g:Profiler-a web server for functional interpretation of gene lists (2016 update). Nucleic Acids Res, 44(W1), W83–89. 10.1093/nar/gkw199

Robinson, M. D., McCarthy, D. J., & Smyth, G. K. (2010). edgeR: a Bioconductor package for differential expression analysis of digital gene expression data. Bioinformatics, 26(1), 139–140. 10.1093/bioinformatics/btp616

Robinson, M. D., & Smyth, G. K. (2007). Moderated statistical tests for assessing differences in tag abundance. Bioinformatics, 23(21), 2881–2887. 10.1093/bioinformatics/btm453

Sala, V., Gallo, S., Leo, C., Gatti, S., Gelb, B. D., & Crepaldi, T. (2012). Signaling to Cardiac Hypertrophy: Insights from Human and Mouse RASopathies. Molecular Medicine, 18(6), 938–947. 10.2119/molmed.2011.00512

Scherle, W. (1970). A simple method for volumetry of organs in quantitative stereology. Mikroskopie, 26(0026-3702), 57–60.

Schuler, B., Arras, M., Keller, S., Rettich, A., Lundby, C., Vogel, J., & Gassmann, M. (2010). Optimal hematocrit for maximal exercise performance in acute and chronic erythropoietin-treated mice. Proceedings of the National Academy of Sciences, 107(1), 419–423. 10.1073/pnas.0912924107

Scott, G. R., & Dalziel, A. C. (2021). Physiological insight into the evolution of complex phenotypes: aerobic performance and the O2 transport pathway of vertebrates. Journal of Experimental Biology, 224(16). 10.1242/jeb.210849

Scott, G. R., Elogio, T. S., Lui, M. A., Storz, J. F., & Cheviron, Z. A. (2015). Adaptive Modifications of Muscle Phenotype in High-Altitude Deer Mice Are Associated with Evolved Changes in Gene Regulation. (1537–1719 (Electronic)).

Sears, M. W., Hayes, J. P., O’Connor, C. S., Geluso, K., & Sedinger, J. S. (2006). Individual variation in thermogenic capacity affects above-ground activity of high-altitude Deer Mice. Functional Ecology, 20(1), 97–104. 10.1111/j.1365-2435.2006.01067.x

Soliz, J., Joseph, V., Soulage, C., Becskei, C., Vogel, J., Pequignot, J. M., Ogunshola, O., & Gassmann, M. (2005). Erythropoietin regulates hypoxic ventilation in mice by interacting with brainstem and carotid bodies. The Journal of Physiology, 568(2), 559–571. 10.1113/jphysiol.2005.093328

Storz, J. F. (2016). Gene Duplication and Evolutionary Innovations in Hemoglobin-Oxygen Transport. Physiology, 31(3), 223–232. 10.1152/physiol.00060.2015

Storz, J. F. (2019). Hemoglobin: insights into protein structure, function, and evolution. Oxford University Press. 10.1093/oso/9780198810681.001.0001

Storz, J. F., & Bautista, N. M. (2022). Altitude acclimatization, hemoglobin-oxygen affinity, and circulatory oxygen transport in hypoxia. Molecular Aspects of Medicine, 84, 101052. 10.1016/j.mam.2021.101052

Storz, J. F., Runck, A. M., Moriyama, H., Weber, R. E., & Fago, A. (2010). Genetic differences in hemoglobin function between highland and lowland deer mice. Journal of Experimental Biology, 213(15), 2565–2574. 10.1242/jeb.042598

Storz, J. F., Runck, A. M., Sabatino, S. J., Kelly, J. K., Ferrand, N., Moriyama, H., Weber, R. E., & Fago, A. (2009). Evolutionary and functional insights into the mechanism underlying high-altitude adaptation of deer mouse hemoglobin. Proceedings of the National Academy of Sciences, 106(34), 14450–14455. 10.1073/pnas.0905224106

Storz, J. F., & Scott, G. R. (2019). Life Ascending: Mechanism and Process in Physiological Adaptation to High-Altitude Hypoxia. *Annual Review of Ecology*, Evolution, and Systematics, 50(1), 503–526. 10.1146/annurev-ecolsys-110218-025014

Storz, J. F., Scott, G. R., & Cheviron, Z. A. (2010). Phenotypic plasticity and genetic adaptation to high-altitude hypoxia in vertebrates. Journal of Experimental Biology, 213(24), 4125–4136. 10.1242/jeb.048181

Sylvester, J. T., Shimoda, L. A., Aaronson, P. I., & Ward, J. P. T. (2012). Hypoxic Pulmonary Vasoconstriction. Physiological reviews, 92(1), 367–520. 10.1152/physrev.00041.2010

Tate, K. B., Ivy, C. M., Velotta, J. P., Storz, J. F., McClelland, G. B., Cheviron, Z. A., & Scott, G. R. (2017). Circulatory mechanisms underlying adaptive increases in thermogenic capacity in high-altitude deer mice. Journal of Experimental Biology, 220(20), 3616–3620. 10.1242/jeb.164491

Tate, K. B., Wearing, O. H., Ivy, C. M., Cheviron, Z. A., Storz, J. F., McClelland, G. B., & Scott, G. R. (2020). Coordinated changes across the O_2_ transport pathway underlie adaptive increases in thermogenic capacity in high-altitude deer mice. Proceedings of the Royal Society B: Biological Sciences, 287(1927), 20192750. 10.1098/rspb.2019.2750

Thibert-Plante, X., & Hendry, A. P. (2011). The consequences of phenotypic plasticity for ecological speciation. Journal of Evolutionary Biology, 24(2), 326–342. 10.1111/j.1420-9101.2010.02169.x

Timón-Gómez, A., Nývltová, E., Abriata, L. A., Vila, A. J., Hosler, J., & Barrientos, A. (2019). Mitochondrial cytochrome c oxidase biogenesis: Recent developments. Semin Cell Dev Biol, 76(1096-3634 (Electronic)). 10.1016/j.semcdb.2017.08.055.

Velotta, J. P., Ivy, C. M., Wolf, C. J., Scott, G. R., & Cheviron, Z. A. (2018). Maladaptive phenotypic plasticity in cardiac muscle growth is suppressed in high-altitude deer mice. Evolution, 72(12), 2712–2727. 10.1111/evo.13626

Wearing, O. H., Ivy, C. M., Gutiérrez-Pinto, N., Velotta, J. P., Campbell-Staton, S. C., Natarajan, C., Cheviron, Z. A., Storz, J. F., & Scott, G. R. (2021). The adaptive benefit of evolved increases in hemoglobin-O2 affinity is contingent on tissue O2 diffusing capacity in high-altitude deer mice. BMC Biology, 19(1). 10.1186/s12915-021-01059-4

Wearing, O. H., & Scott, G. R. (2021). Commentary: Hierarchical reductionism approach to understanding adaptive variation in animal performance. Comparative Biochemistry and Physiology Part B: Biochemistry and Molecular Biology, 256, 110636. 10.1016/j.cbpb.2021.110636

West, C. M., Wearing, O. H., Rhem, R. G., & Scott, G. R. (2021). Pulmonary hypertension is attenuated and ventilation-perfusion matching is maintained during chronic hypoxia in deer mice native to high altitude. *American Journal of Physiology-Regulatory*, Integrative and Comparative Physiology, 320(6), R800–R811. 10.1152/ajpregu.00282.2020

West CM, Ivy CM, Husnudinov R, Scott GR. 2021. Evolution and developmental plasticity of lung structure in high-altitude deer mice. J Comp Physiol B. 191, 385–396.

Young, J. M., Williams, D. R., & Thompson, A. A. R. (2019). Thin Air, Thick Vessels: Historical and Current Perspectives on Hypoxic Pulmonary Hypertension. Frontiers in Medicine, 6. 10.3389/fmed.2019.00093

Zhang, B., & Hovarth, S. (2005). A general framework for weighted gene co-expression network analysis. Stat Appl Genet Mol Biol, 4.

Zhang, X.-J., Zhang, P., & Li, H. (2015). Interferon Regulatory Factor Signalings in Cardiometabolic Diseases. Hypertension, 66(2), 222–247. 10.1161/hypertensionaha.115.04898

Zhang, Y., Park, C., Bennett, C., Thornton, M., & Kim, D. (2021). Rapid and accurate alignment of nucleotide conversion sequencing reads with HISAT-3N. Genome Res, 31(7), 1290–1295. 10.1101/gr.275193.120

