## Supplementary Material for "Local adaptation, plasticity, and evolved resistance to hypoxic cold stress in high-altitude deer mice"

**Fig. S1.** Variation in organ masses in acclimated and control mice from different elevations. CL = control, lowland native, CH = control, highland native, AL = acclimated, lowland native, and AH = acclimated, highland native. Data are expressed as mean ± s.e.m. and individual values are over imposed. Statistical significance was considered with an $\alpha \leq 0.05$. n = 13-16 mice per data point for all organs with exception of panel 'N' in which n = 8 for all groups.

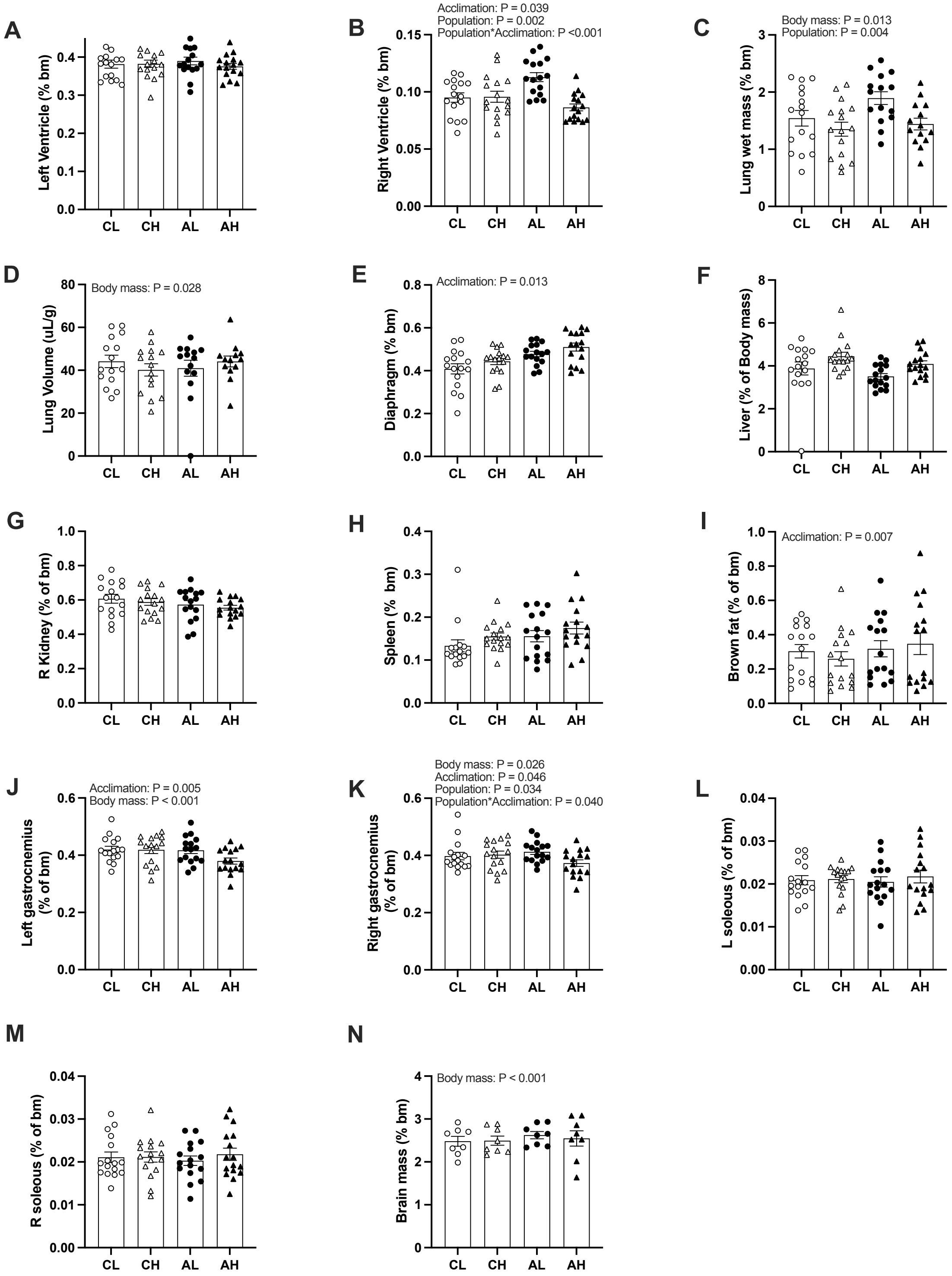

**Fig. S2.** Variation in enzyme activities in acclimated and control mice from different elevations. Enzyme activities were measured in the left ventricle (LV), right ventricle (RV), gastrocnemius, and diaphragm. HOAD = β-hydroxyacyl CoA dehydrogenase; COX =Cytochrome oxidase; CS = Citrate Synthase; LDH = Lactate dehydrogenase. CL = control, lowland native, CH = control, highland native, AL = acclimated, lowland native, and AH = acclimated, highland native. Data are expressed as mean ± s.e.m. n = 8 mice per treatment.

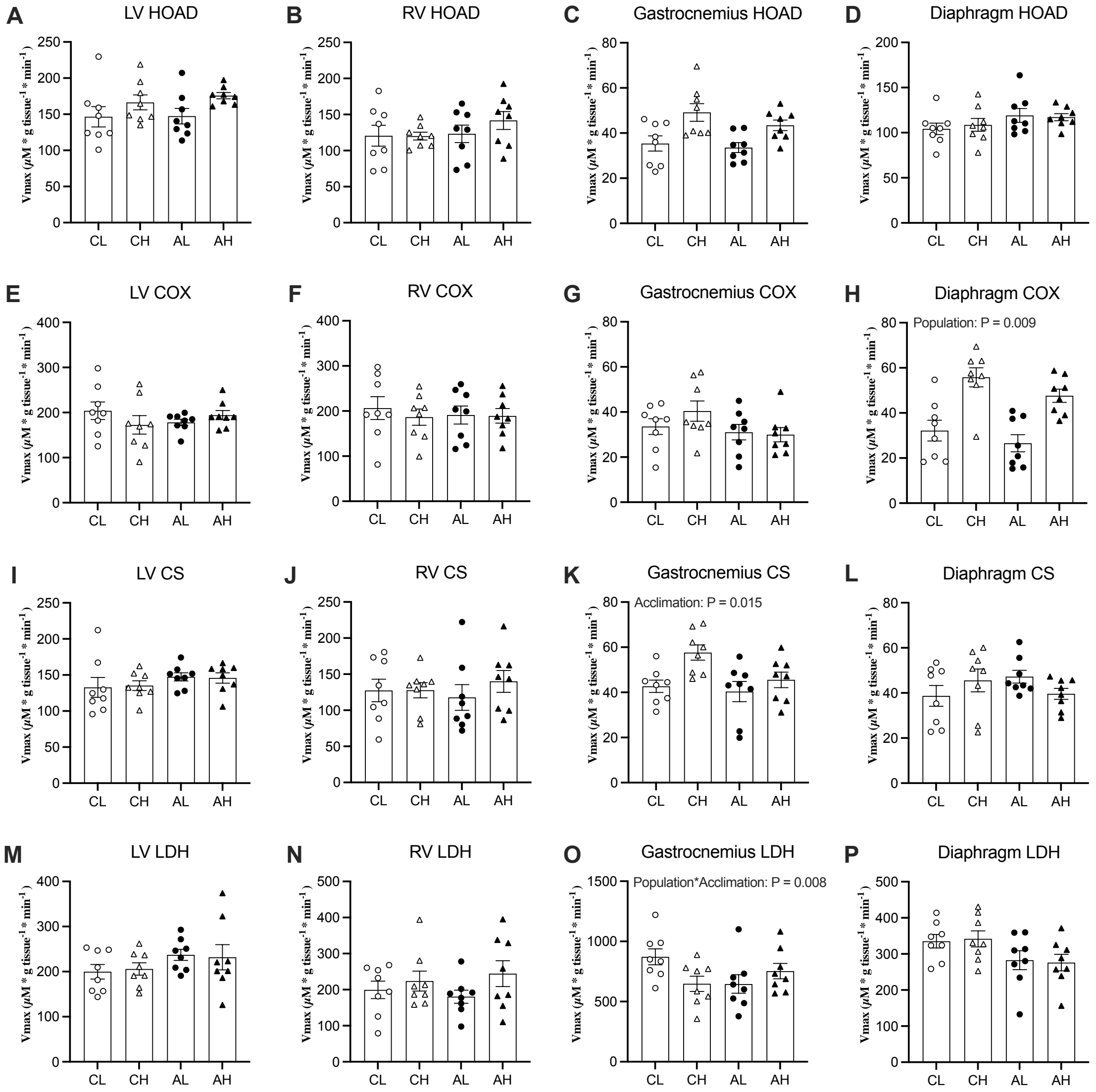

**Fig. S3.** Variation in: A) Body mass (g) of all mice remained essentially unchanged during the course of the experiment; B) Food, and C) Water consumption; D) Hemoglobin concentration; E) Hill's cooperativity coefficient, in acclimated and control mice throughout experimentation. Data are expressed as mean ± s.e.m. N = 15-16 mice per data point. N = 3–5 holding cages in B and C. Statistical significance was considered with an $\alpha \leq 0.05$.

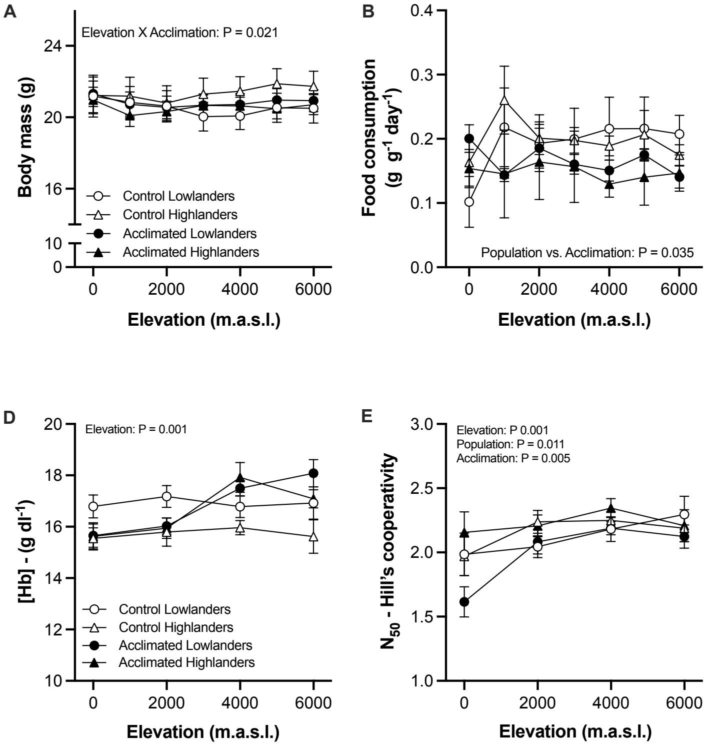

**Fig. S4.** PCA of RNASeq samples with 95% ellipse: A) lung and B) right ventricle.

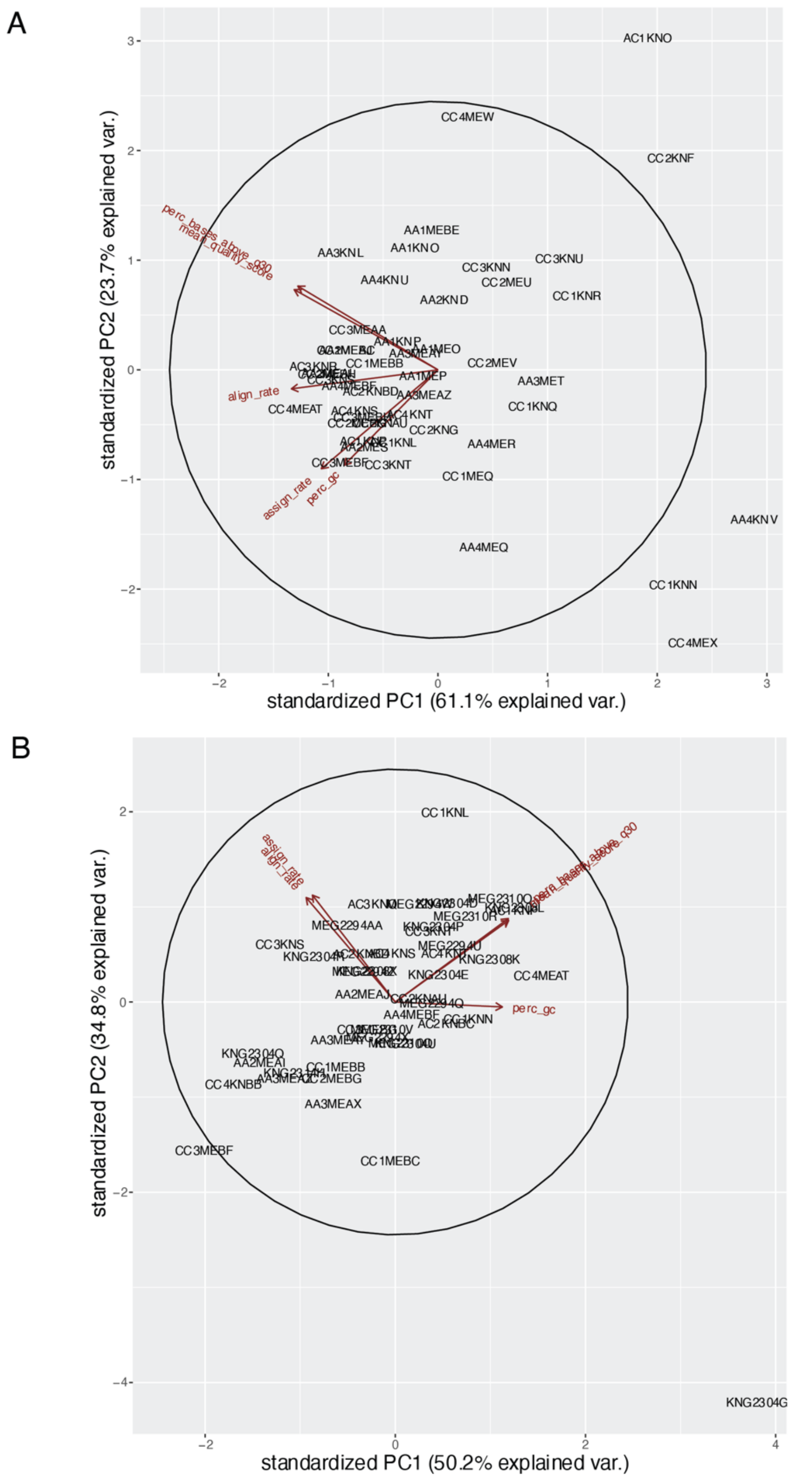

**Fig. S5.** Scale free topology model fit for: A) lung and B) right ventricle.

A

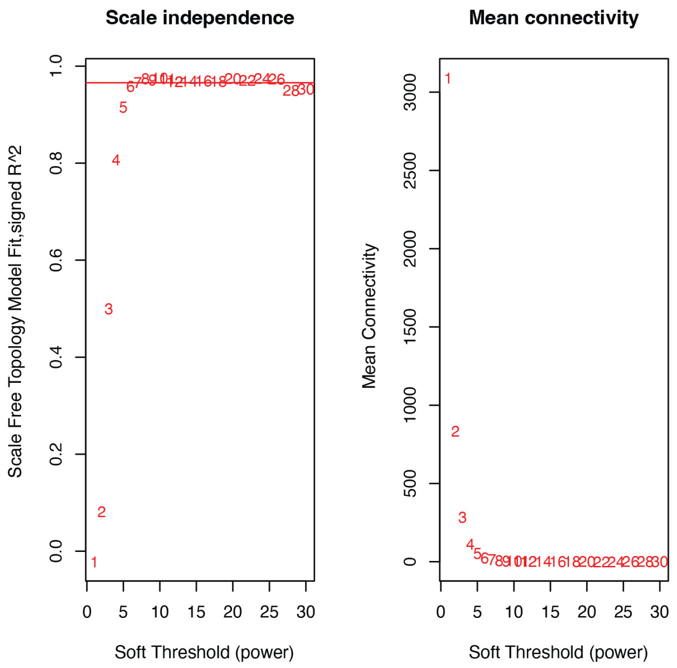

B

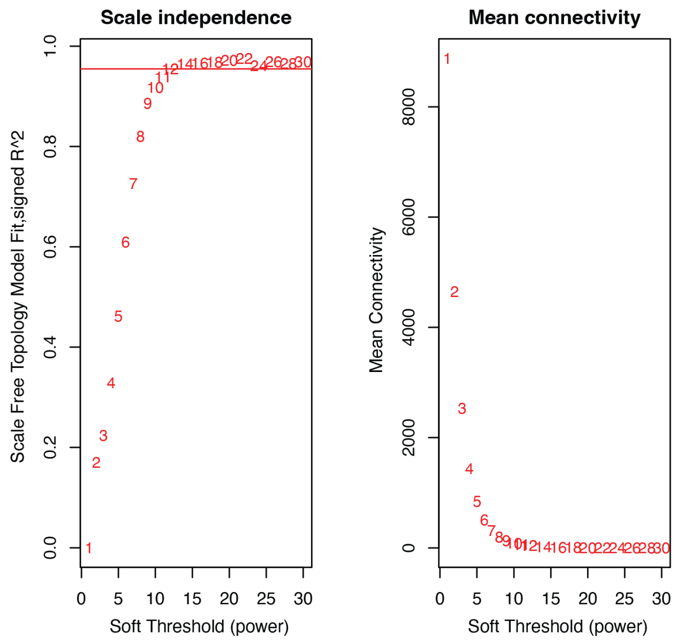

**Fig. S6.** Module expression for lung and right ventricle that show a significant effect of population and/or treatment across the Lung (L modules n =16) and right ventricle (RV modules n = 9).

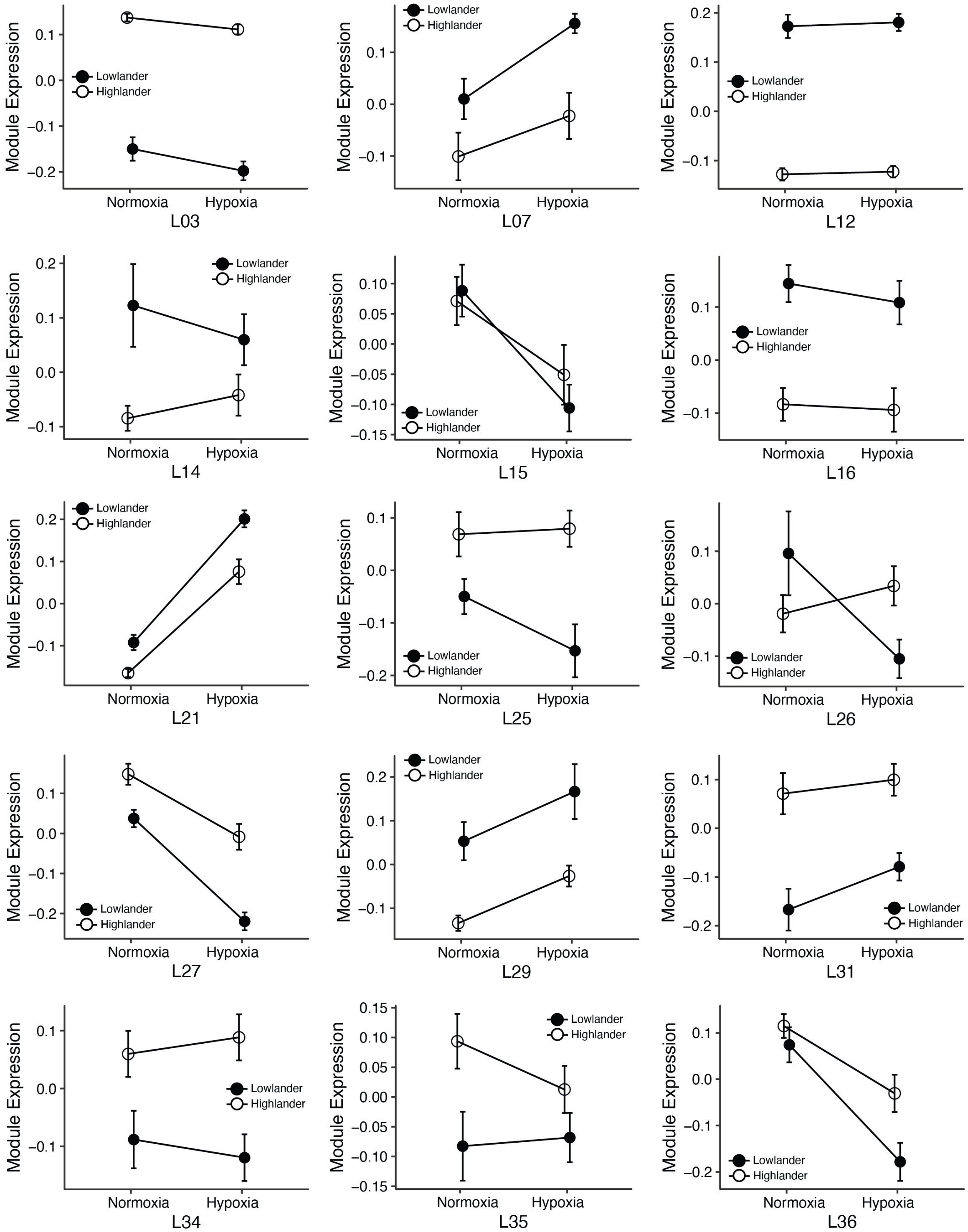

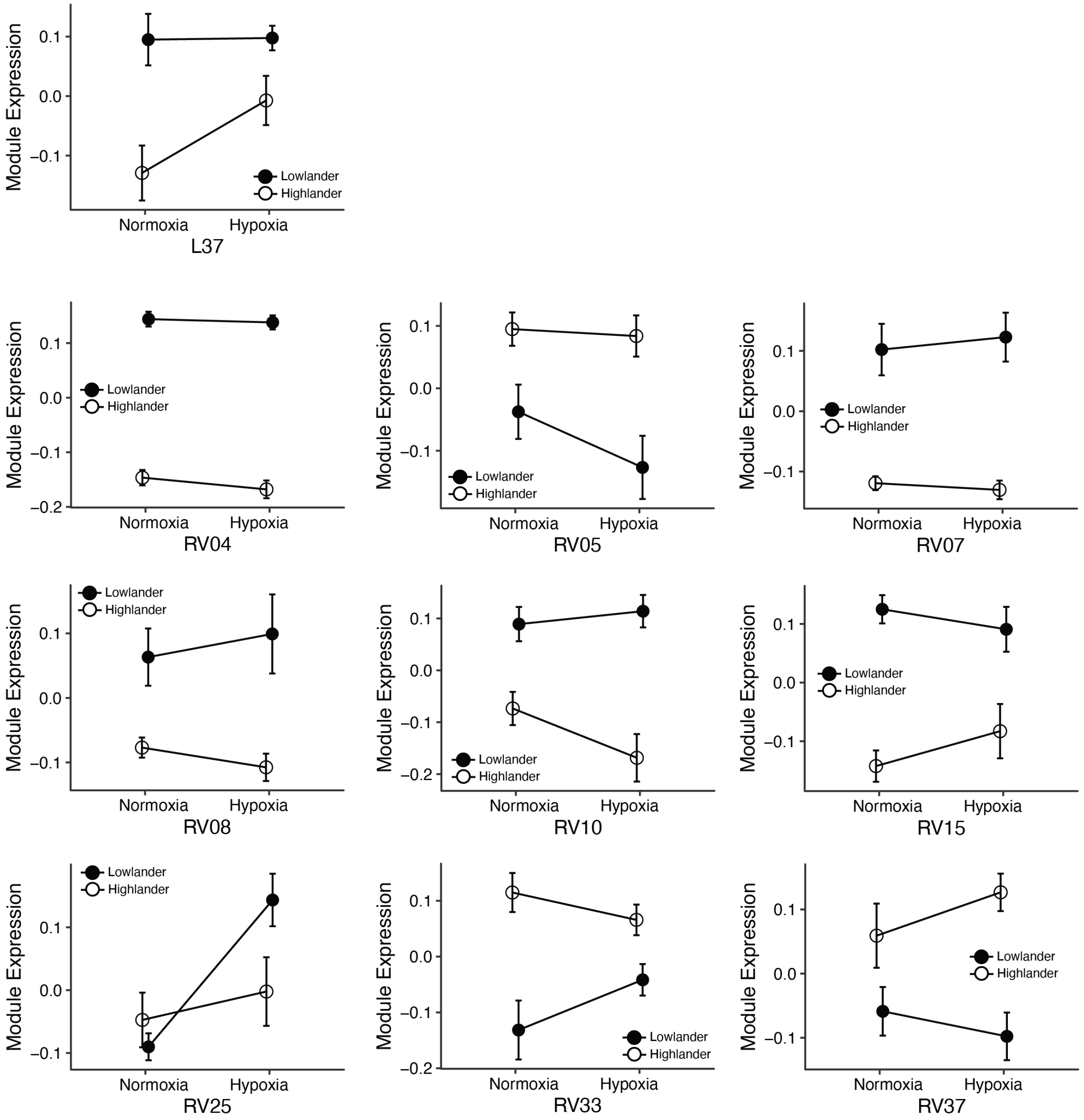
Module Expression
Normoxia
Hypoxia
Lowlander
Highlander
L37
RV04
RV05
RV07
RV08
RV10
RV15
RV25
RV33
RV37

**Table S1.** Results of statistical linear models to examine variation in physiological traits in acclimated and control mice from different elevations; thermogenic $\dot{V}O_{2max}$, breathing frequency ($f_R$), carbon dioxide production ($\dot{V}_{CO2}$), tidal volume ($V_T$), respiratory exchange ratio (RER), total ventilation ($\dot{V}_E$), air convection requirement for oxygen (ACR-$O_2$), air convection requirement for carbon dioxide (ACR-$CO_2$), pulmonary $O_2$ extraction ($\dot{V}O_{2max}/(\dot{V}_E*F_iO_2)$), arterial oxygen saturation ($S_aO_2$), hemoglobin oxygen affinity ($P_{50}$), hematocrit (Hct), heart rate ($f_H$), Hill's cooperativity coefficient ($N_{50}$), hemoglobin concentration ([Hb]), body mass, water consumption, and food consumption. Significant results among groups are highlighted in bold font. '*' Signify rank transformation. Statistical significance was considered with an $\alpha \leq 0.05$.

| Trait | Population vs Acclimation vs Elevation | Population vs Elevation | Population vs Acclimation | Elevation vs Acclimation | Acclimation | Elevation | Population | Replicate | Sex | Body mass |
|---|---|---|---|---|---|---|---|---|---|---|
| *$VO_{2max}$ | F = 0.512 | **F = 2.781** | F = 0.434 | **F = 2.851** | **F = 0.013** | F = 1.012 | F = 2.042 | **F = 20.77** | F = 1.690 | **F = 10.56** |
| | P = 0.799 | **P = 0.011** | P = 0.510 | **P = 0.010** | **P = 0.907** | P = 0.416 | P = 0.163 | **P < 0.001** | P = 0.197 | **P = 0.001** |
| *$f_R$ | F = 1.411 | F = 1.826 | **F = 4.832** | **F = 4.101** | **F = 5.365** | **F = 2.794** | F = 0.759 | **F = 13.55** | F = 0.148 | F = 0.577 |
| | P = 0.241 | P = 0.143 | **P = 0.029** | **P = 0.007** | **P = 0.021** | **P = 0.041** | P = 0.386 | **P = 0.010** | P = 0.700 | P = 0.449 |
| *$VCO_2$ | F = 0.852 | **F = 4.111** | F = 0.001 | **F = 3.442** | F = 1.331 | F = 1.895 | F = 0.073 | F = 1.639 | F = 1.450 | **F = 4.425** |
| | P = 0.466 | **P = 0.037** | P = 0.966 | **P = 0.017** | P = 0.250 | P = 0.132 | P = 0.789 | P = 0.218 | P = 0.232 | **P = 0.037** |
| $V_T$ | F = 1.149 | F = 0.267 | F = 0.947 | F = 0.547 | F = 1.622 | F = 1.988 | F = 1.805 | F = 0.308 | **F = 6.002** | **F = 17.86** |
| | P = 0.330 | P = 0.849 | P = 0.331 | P = 0.650 | P = 0.204 | P = 0.117 | P = 0.182 | P = 0.595 | **P = 0.017** | **P < 0.001** |
| *RER | F = 1.608 | F = 1.539 | F = 0.261 | F = 1.667 | F = 0.263 | F = 0.063 | F = 2.421 | **F = 53.72** | F = 0.101 | F = 0.500 |
| | P = 0.189 | P = 0.205 | P = 0.609 | P = 0.175 | P = 0.608 | P = 0.978 | P = 0.123 | **P < 0.001** | P = 0.751 | P = 0.481 |
| *$V_E$ | F = 0.682 | F = 0.118 | F = 0.614 | F = 0.704 | F = 1.045 | F = 1.720 | F = 0.291 | F = 2.540 | F = 3.910 | **F = 18.89** |
| | P = 0.563 | P = 0.949 | P = 0.433 | P = 0.550 | P = 0.307 | P = 0.164 | P = 0.590 | P = 0.163 | P = 0.052 | **P < 0.001** |
| *ACR-$O_2$ | F = 1.124 | F = 0.939 | F = 0.129 | F = 0.669 | F = 0.831 | F = 2.295 | F = 2.652 | **F = 8.744** | F = 0.019 | F = 0.732 |
| | P = 0.340 | P = 0.422 | P = 0.719 | P = 0.572 | P = 0.362 | P = 0.079 | P = 0.107 | **P = 0.026** | P = 0.890 | P = 0.395 |
| *ACR-$CO_2$ | F = 1.888 | F = 0.866 | F = 1.392 | F = 0.653 | F = 0.128 | F = 2.558 | F = 0.107 | F = 0.259 | F = 0.081 | F = 1.183 |
| | P = 0.133 | P = 0.459 | P = 0.239 | P = 0.581 | P = 0.720 | P = 0.056 | P = 0.743 | P = 0.629 | P = 0.775 | P = 0.280 |
| Pulmonary $O_2$ extraction | F = 2.095 | F = 0.951 | F = 0.198 | F = 0.480 | F = 0.791 | **F = 6.803** | F = 0.506 | **F = 10.42** | F = 0.020 | F = 0.530 |
| | P = 0.102 | P = 0.417 | P = 0.656 | P = 0.696 | P = 0.374 | **P < 0.001** | P = 0.478 | **P = 0.018** | P = 0.886 | P = 0.469 |
| *$S_aO_2$ | F = 1.294 | **F = 2.915** | F = 2.044 | F = 2.051 | F = 0.146 | **F = 7.207** | F = 1.744 | F = 0.873 | F = 2.921 | F = 0.024 |
| | P = 0.258 | **P = 0.008** | P = 0.153 | P = 0.058 | P = 0.702 | **P < 0.001** | P = 0.189 | P = 0.373 | P = 0.092 | P = 0.875 |
| *$P_{50}$ | F = 2.383 | **F = 5.007** | F = 2.758 | **F = 3.390** | **F = 16.38** | **F = 12.92** | F = 0.002 | **F = 256.7** | F = 0.375 | - |
| | P = 0.070 | **P = 0.002** | P = 0.098 | **P = 0.019** | **P < 0.001** | **P < 0.001** | P = 0.962 | **P < 0.001** | P = 0.542 | - |
| *Hct | F = 0.520 | F = 1.730 | F = 0.164 | **F = 3.635** | F = 0.078 | **F = 11.37** | F = 0.252 | **F = 31.37** | **F = 4.603** | - |
| | P = 0.668 | P = 0.162 | P = 0.685 | **P = 0.013** | P = 0.779 | **P < 0.001** | P = 0.616 | **P < 0.001** | **P = 0.035** | - |
| *$f_H$ | F = 0.548 | F = 1.180 | F = 0.094 | F = 1.577 | F = 0.033 | F = 1.503 | F = 0.382 | **F = 6.593** | F = 0.320 | F = 0.031 |
| | P = 0.771 | P = 0.315 | P = 0.758 | P = 0.152 | P = 0.855 | P = 0.175 | P = 0.537 | **P = 0.033** | P = 0.573 | P = 0.858 |
| *$N_{50}$ | F = 0.734 | F = 0.460 | F = 3.061 | F = 1.884 | **F = 7.975** | **F = 5.561** | **F = 6.564** | **F = 84.87** | F = 0.063 | - |
| | P = 0.532 | P = 0.710 | P = 0.081 | P = 0.133 | **P = 0.005** | **P = 0.001** | **P = 0.011** | **P < 0.001** | P = 0.801 | - |
| *[Hb] | F = 0.559 | F = 0.508 | F = 3.339 | F = 2.275 | F = 1.857 | **F = 5.202** | F = 0.130 | **F = 10.23** | F = 1.456 | - |
| | P = 0.642 | P = 0.676 | P = 0.068 | P = 0.081 | P = 0.174 | **P = 0.001** | P = 0.720 | **P = 0.006** | P = 0.231 | - |
| *Body mass | F = 1.502 | F = 0.567 | F = 0.701 | **F = 2.504** | F = 0.173 | F = 1.408 | F = 1.514 | F = 2.183 | **F = 38.37** | - |
| | P = 0.176 | P = 0.756 | P = 0.404 | **P = 0.021** | P = 0.678 | P = 0.210 | P = 0.241 | P = 0.158 | **P < 0.001** | - |
| *Water consumption | F = 0.346 | F = 0.572 | F = 0.595 | F = 0.769 | F = 0.108 | F = 0.878 | F = 1.764 | - | F = 0.095 | - |
| | P = 0.910 | P = 0.751 | P = 0.442 | P = 0.596 | P = 0.742 | P = 0.514 | P = 0.191 | - | P = 0.758 | - |
| *Food consumption | F = 0.772 | F = 0.781 | **F = 4.576** | F = 0.636 | F = 0.002 | F = 0.381 | F = 0.841 | - | F = 1.683 | - |
| | P = 0.593 | P = 0.586 | **P = 0.035** | P = 0.700 | P = 0.958 | P = 0.889 | P = 0.375 | - | P = 0.197 | - |

**Table S2.** Results of statistical linear models to examine variation in organ mases and tissues of acclimated and control mice from different elevations. Significant results among groups are highlighted in bold font. '*' Signify rank transformation. Statistical significance was considered with an $\alpha \leq 0.05$.

| Organ/Tissue | Population vs Acclimation | Population | Acclimation | Sex | Body Mass | Replicate |
|---|---|---|---|---|---|---|
| Left ventricle | F = 1.431<br>P = 0.236 | F = 1.148<br>P = 0.308 | F = 1.040<br>P = 0.312 | F = 0.261<br>P = 0.611 | F = 1.700<br>P = 0.197 | F = 0.634<br>P = 0.436 |
| Right ventricle | **F = 14.03**<br>**P < 0.001** | **F = 15.63**<br>**P = 0.002** | **F = 4.458**<br>**P = 0.039** | F = 0.345<br>P = 0.558 | F = 0.438<br>P = 0.510 | F = 0.873<br>P = 0.372 |
| Lungs wet mass | F = 2.586<br>P = 0.113 | **F = 11.49**<br>**P = 0.004** | F = 0.910<br>P = 0.344 | F = 0.298<br>P = 0.587 | **F = 6.579**<br>**P = 0.013** | **F = 70.87**<br>**P < 0.001** |
| Lung volume | F = 0.863<br>P = 0.357 | F = 0.013<br>P = 0.909 | F = 1.633<br>P = 0.207 | F = 0.006<br>P = 0.936 | **F = 5.075**<br>**P = 0.028** | F = 4.038<br>P = 0.069 |
| Diaphragm* | F = 0.000<br>P = 0.991 | F = 0.607<br>P = 0.451 | **F = 6.452**<br>**P = 0.013** | F = 1.042<br>P = 0.311 | F = 0.907<br>P = 0.345 | F = 3.995<br>P = 0.077 |
| Liver* | F = 0.378<br>P = 0.540 | F = 2.716<br>P = 0.127 | F = 2.006<br>P = 0.162 | F = 1.357<br>P = 0.248 | F = 4.488<br>P = 0.038 | F = 0.813<br>P = 0.380 |
| Kidney | F = 0.084<br>P = 0.772 | F = 0.324<br>P = 0.580 | F = 3.102<br>P = 0.083 | F = 1.567<br>P = 0.215 | F = 0.507<br>P = 0.479 | F = 0.817<br>P = 0.383 |
| Spleen* | F = 0.110<br>P = 0.741 | F = 1.046<br>P = 0.326 | F = 0.366<br>P = 0.547 | F = 0.389<br>P = 0.535 | F = 0.000<br>P = 0.991 | F = 0.198<br>P = 0.668 |
| Interscapular Brown fat* | F = 1.520<br>P = 0.222 | F = 0.255<br>P = 0.875 | **F = 7.818**<br>**P = 0.007** | F = 3.318<br>P = 0.073 | F = 3.945<br>P = 0.052 | **F = 12.03**<br>**P = 0.008** |
| Left gastrocnemius | F = 3.825<br>P = 0.055 | F = 4.182<br>P = 0.063 | **F = 8.336**<br>**P = 0.005** | F = 1.629<br>P = 0.206 | **F = 15.07**<br>**P < 0.001** | F = 2.325<br>P = 0.166 |
| Right gastrocnemius | **F = 4.381**<br>**P = 0.040** | **F = 5.607**<br>**P = 0.034** | **F = 4.140**<br>**P = 0.046** | F = 0.830<br>P = 0.366 | **F = 5.282**<br>**P = 0.026** | F = 1.405<br>P = 0.278 |
| Left soleus | F = 0.419<br>P = 0.519 | F = 0.633<br>P = 0.441 | F = 0.360<br>P = 0.550 | F = 0.579<br>P = 0.449 | F = 2.113<br>P = 0.152 | F = 0.234<br>P = 0.641 |
| Right soleus | F = 0.653<br>P = 0.422 | F = 0.699<br>P = 0.419 | F = 0.368<br>P = 0.546 | F = 0.729<br>P = 0.396 | F = 2.424<br>P = 0.126 | F = 0.000<br>P = 0.983 |
| Brain | F = 0.009<br>P = 0.922 | F = 1.141<br>P = 0.337 | F = 0.090<br>P = 0.766 | F = 0.237<br>P = 0.632 | **F = 37.58**<br>**P < 0.001** | -<br>- |

**Table S3.** Results of statistical linear models to examine variation in enzyme activity in the left ventricle, right ventricle, gastrocnemius, and diaphragm in acclimated and control mice from different elevations. Tissues were obtained at the end of the experimental period of replicate one. HOAD = Cytochrome oxidase; COX = β-hydroxyacyl CoA dehydrogenase; CS = Citrate Synthase; LDH = Lactate dehydrogenase. '*' Signify rank transformation. Significant results are highlighted in bold font. Statistical significance was considered with $\alpha \leq 0.05$.

| Tissue | Enzyme | Population vs Acclimation | Population | Acclimation | Sex | Mass |
|---|---|---|---|---|---|---|
| Left ventricle | HOAD | F = 0.423<br>P = 0.521 | F = 3.243<br>P = 0.116 | F = 0.658<br>P = 0.425 | F = 0.888<br>P = 0.355 | F = 1.126<br>P = 0.299 |
| | COX | F = 1.233<br>P = 0.277 | F = 0.565<br>P =0.482 | F = 0.512<br>P = 0.481 | F = 0.254<br>P = 0.618 | F = 2.776<br>P = 0.108 |
| | CS | F = 0.014<br>P = 0.905 | F = 0.031<br>P = 0.865 | F = 1.412<br>P = 0.246 | F = 1.930<br>P = 0.177 | F = 0.442<br>P = 0.512 |
| | LDH | F = 0.028<br>P = 0.867 | F = 0.047<br>P = 0.834 | F = 1.125<br>P = 0.299 | F = 0.641<br>P = 0.431 | F = 0.212<br>P = 0.649 |
| Rigth ventricle | HOAD | F = 0.496<br>P = 0.487 | F = 1.275<br>P = 0.297 | F = 1.527<br>P = 0.228 | F = 1.125<br>P = 0.299 | F = 0.097<br>P = 0.757 |
| | COX | F = 0.153<br>P = 0.698 | F = 0.002<br>P = 0.961 | F = 0.013<br>P = 0.909 | F = 0.195<br>P = 0.662 | F = 0.007<br>P = 0.933 |
| | CS | F = 0.809<br>P = 0.376 | F = 0.272<br>P = 0.618 | F = 1.402<br>P = 0.248 | F = 1.744<br>P = 0.199 | F = 0.648<br>P = 0.428 |
| | LDH | F = 0.548<br>P = 0.465 | F = 0.237<br>P = 0.641 | F = 2.631<br>P = 0.118 | F = 0.021<br>P = 0.885 | F = 0.088<br>P = 0.768 |
| Left Gastrocnemius | HOAD | F = 0.067<br>P = 0.796 | F = 3.700<br>P = 0.134 | F = 0.677<br>P = 0.418 | F = 1.891<br>P = 0.182 | F = 0.0857<br>P = 0.772 |
| | COX | F = 0.772<br>P = 0.387 | F = 0.063<br>P = 0.810 | F = 3.372<br>P = 0.078 | F = 0.303<br>P = 0.586 | F = 0.580<br>P = 0.453 |
| | CS | F = 2.650<br>P = 0.116 | F = 0.532<br>P = 0.507 | **F = 6.748**<br>**P = 0.015** | F = 3.031<br>P = 0.094 | F = 0.107<br>P = 0.745 |
| | LDH | **F = 8.300**<br>**P = 0.008** | F = 0.773<br>P = 0.429 | F = 2.468<br>P = 0.129 | F = 2.310<br>P = 0.141 | F = 3.699<br>P = 0.066 |
| Diaphragm | HOAD | F = 0.117<br>P = 0.734 | F = 0.779<br>P = 0.407 | F = 0.013<br>P = 0.907 | F = 0.649<br>P = 0.428 | F = 0.191<br>P = 0.665 |
| | COX | F = 0.085<br>P = 0.772 | **F = 12.83**<br>**P = 0.009** | F = 1.906<br>P = 0.180 | F = 0.118<br>P = 0.733 | F = 0.001<br>P = 0.976 |
| | CS | F = 2.259<br>P = 0.145 | F = 1.275<br>P = 0.297 | F = 1.319<br>P = 0.262 | F = 0.106<br>P = 0.747 | F = 0.122<br>P = 0.299 |
| | LDH | F = 0.094<br>P = 0.761 | F = 3.142<br>P = 0.133 | F = 0.066<br>P = 0.799 | F = 0.233<br>P = 0.633 | F = 0.081<br>P = 0.778 |

**Table S4** Sequence quality metrics, mapping, and assignment rates for A) lung tissue, and B) right ventricle.

A

| ***P. maniculatus* Lung (N = 48)** | |
|---|---|
| **Mean Quality Score** | |
| min | 35.54 |
| max | 35.86 |
| mean (sd) | 35.72 ± 0.08 |
| **Percent Bases Above Q30** | |
| min | 92.4 |
| median | 93.4 |
| max | 93.9 |
| mean (sd) | 93.33 ± 0.37 |
| **GC (%)** | |
| min | 48.6 |
| median | 50.2 |
| max | 51.2 |
| mean (sd) | 50.17 ± 0.54 |
| **Reads (millions)** | |
| min | 33.9 |
| median | 47 |
| max | 65 |
| mean (sd) | 46.84 ± 6.1 |
| **Genome Alignment Rate (%)** | |
| min | 93.9 |
| max | 95.5 |
| mean (sd) | 94.93 ± 0.38 |
| **Assignment Rate (%)** | |
| min | 71.6 |
| max | 77.9 |
| mean (sd) | 75.92 ± 1.76 |

B

| ***P. maniculatus* RV (N = 45)** | |
|---|---|
| **Mean Quality Score** | |
| min | 35.65 |
| max | 33.85 |
| mean (sd) | 33.75 ± 0.05 |
| **Percent Bases Above Q30** | |
| min | 92.7 |
| median | 93.3 |
| max | 93.8 |
| mean (sd) | 93.30 ± 0.26 |
| **GC (%)** | |
| min | 46.5 |
| median | 47.5 |
| max | 48.5 |
| mean (sd) | 47.49 ± 0.48 |
| **Reads (millions)** | |
| min | 32.4 |
| median | 38.7 |
| max | 48.5 |
| mean (sd) | 38.36 ± 3.14 |
| **Genome Alignment Rate (%)** | |
| min | 95.2 |
| max | 96.5 |
| mean (sd) | 96.09 ± 0.24 |
| **Assignment Rate (%)** | |
| min | 81.9 |
| max | 84.8 |
| mean (sd) | 83.03 ± 0.56 |

**Table S5.** WGCNA modules.

| Lung | | Right ventricle | |
|---|---|---|---|
| **Module** | **# Genes** | **Module** | **# Genes** |
| L0 | 2119 | RV0 | 3379 |
| L01 | 2193 | RV1 | 2001 |
| L02 | 1286 | RV2 | 1057 |
| L03 | 935 | RV3 | 944 |
| L04 | 917 | RV4 | 917 |
| L05 | 875 | RV5 | 748 |
| L06 | 852 | RV6 | 558 |
| L07 | 836 | RV7 | 538 |
| L08 | 766 | RV8 | 426 |
| L09 | 643 | RV9 | 399 |
| L10 | 638 | RV10 | 382 |
| L11 | 590 | RV11 | 312 |
| L12 | 485 | RV12 | 245 |
| L13 | 462 | RV13 | 220 |
| L14 | 454 | RV14 | 214 |
| L15 | 293 | RV15 | 179 |
| L16 | 240 | RV16 | 161 |
| L17 | 236 | RV17 | 152 |
| L18 | 232 | RV18 | 149 |
| L19 | 224 | RV19 | 137 |
| L20 | 218 | RV20 | 135 |
| L21 | 214 | RV21 | 133 |
| L22 | 207 | RV22 | 132 |
| L23 | 193 | RV23 | 120 |
| L24 | 176 | RV24 | 115 |
| L25 | 159 | RV25 | 114 |
| L26 | 155 | RV26 | 106 |
| L27 | 142 | RV27 | 105 |
| L28 | 119 | RV28 | 105 |
| L29 | 119 | RV29 | 99 |
| L30 | 113 | RV30 | 97 |
| L31 | 110 | RV31 | 92 |
| L32 | 102 | RV32 | 91 |
| L33 | 97 | RV33 | 86 |
| L34 | 87 | RV34 | 73 |
| L35 | 60 | RV35 | 66 |
| L36 | 60 | RV36 | 54 |
| L37 | 59 | RV37 | 49 |
| L38 | 44 | | |
| L39 | 42 | | |

**Table S6.** Lung module expression ANOVA results.

| Module | population | treatment | population*treatment |
|---|---|---|---|
| L01 | 0.2295 | 0.8487 | 0.987 |
| L02 | 0.8583 | 0.7684 | 0.9288 |
| L03 | **0** | 0.1489 | 0.8487 |
| L04 | 0.4287 | 0.8819 | 0.9288 |
| L05 | 0.987 | 0.8487 | 0.8487 |
| L06 | 0.4287 | 0.8819 | 0.987 |
| L07 | **6.00E-04** | **0.0058** | 0.4287 |
| L08 | 0.7717 | 0.6782 | 0.9288 |
| L09 | 0.222 | 0.6717 | 0.8487 |
| L10 | 0.1389 | 0.9288 | 0.8572 |
| L11 | 0.2867 | 0.8487 | 0.8487 |
| L12 | **0** | 0.8583 | 0.992 |
| L13 | 0.3539 | 0.7817 | 0.7792 |
| L14 | **0.0179** | 0.8487 | 0.7717 |
| L15 | 0.8071 | **0.0073** | 0.7977 |
| L16 | **0** | 0.8583 | 0.8487 |
| L17 | 0.7734 | 0.9288 | 0.8487 |
| L18 | 0.8487 | 0.8487 | 0.7717 |
| L19 | 0.4287 | 0.8487 | 0.76 |
| L20 | 0.3297 | 0.3911 | 0.8583 |
| L21 | **0** | **0** | 0.8487 |
| L22 | 0.6717 | 0.8487 | 0.7734 |
| L23 | 0.2478 | 0.8583 | 0.8487 |
| L24 | 0.331 | 0.4287 | 0.6896 |
| L25 | **0.0028** | 0.8487 | 0.7529 |
| L26 | 0.507 | 0.7461 | **0.0397** |
| L27 | **0** | **0** | 0.5164 |
| L28 | 0.8487 | 0.8487 | 0.9428 |
| L29 | **0** | **0.0058** | 0.4287 |
| L30 | 0.987 | 0.4695 | 0.582 |
| L31 | **2.00E-04** | 0.582 | 0.8583 |
| L32 | 0.7717 | 0.9288 | 0.7717 |
| L33 | 0.4287 | 0.8018 | 0.4287 |
| L34 | **5.00E-04** | 0.9441 | 0.8071 |
| L35 | **0.0241** | 0.8487 | 0.6896 |
| L36 | **0.0326** | **1.00E-04** | 0.582 |
| L37 | **0.0023** | 0.4943 | 0.4695 |
| L38 | 0.8583 | 0.4287 | 0.987 |
| L39 | 0.4943 | 0.8572 | 0.987 |

**Table S7.** Lung module association with $S_aO_2$.

| Module | cor | p | p.adjust |
|---|---|---|---|
| **L38** | **-0.5851778** | **5.86E-05** | **0.00234296** |
| L21 | 0.45329906 | 0.0029171 | 0.05834194 |
| L05 | -0.4157323 | 0.00686692 | 0.09155887 |
| L29 | 0.39315625 | 0.01099371 | 0.10993709 |
| L36 | -0.3774894 | 0.01496854 | 0.11974828 |
| L09 | 0.35784212 | 0.02161313 | 0.14408756 |
| L17 | -0.3420112 | 0.02861927 | 0.16353871 |
| L14 | 0.32622907 | 0.03737761 | 0.16853978 |
| L27 | -0.3253535 | 0.03792145 | 0.16853978 |
| L06 | 0.28063535 | 0.07552222 | 0.27462627 |
| L20 | -0.2815922 | 0.07448917 | 0.27462627 |
| L08 | -0.2713314 | 0.08616257 | 0.28720856 |
| L15 | -0.2652424 | 0.0937328 | 0.28840861 |
| L19 | 0.23918975 | 0.13202697 | 0.37721992 |
| L16 | -0.2235908 | 0.15994156 | 0.42651083 |
| L32 | -0.2149642 | 0.17710708 | 0.44276769 |
| L30 | 0.20213968 | 0.20500172 | 0.48235699 |
| L01 | 0.1691371 | 0.29044155 | 0.53524877 |
| L04 | -0.1640674 | 0.30535276 | 0.53524877 |
| L13 | -0.1726103 | 0.28050278 | 0.53524877 |
| L24 | 0.16326121 | 0.30776805 | 0.53524877 |
| L25 | 0.18512066 | 0.24656133 | 0.53524877 |
| L37 | 0.16883785 | 0.29130841 | 0.53524877 |
| L31 | 0.15479478 | 0.33386445 | 0.55644076 |
| L03 | -0.1429754 | 0.37251561 | 0.57310094 |
| L26 | 0.14404497 | 0.36891228 | 0.57310094 |
| L10 | 0.12623685 | 0.43159408 | 0.63939864 |
| L34 | 0.12164599 | 0.44866263 | 0.64094662 |
| L02 | -0.1117893 | 0.48652252 | 0.66797642 |
| L22 | -0.1039734 | 0.51768172 | 0.66797642 |
| L33 | 0.09816695 | 0.54145513 | 0.67681892 |
| L11 | 0.09315525 | 0.56238666 | 0.6816808 |
| L12 | 0.07046548 | 0.66153807 | 0.75604351 |
| L28 | 0.07194718 | 0.65486067 | 0.75604351 |
| L07 | -0.0491341 | 0.76031568 | 0.79228348 |
| L18 | -0.0530004 | 0.74207669 | 0.79228348 |
| L23 | 0.0510308 | 0.75135135 | 0.79228348 |
| L39 | -0.0465714 | 0.77247639 | 0.79228348 |
| L35 | -0.0298676 | 0.85293638 | 0.85293638 |

**Table S8.** Hub genes for the lung and right ventricle modules.

| Lung | | Right Ventricle | |
|---|---|---|---|
| **Module** | **Hub Gene** | **module** | **hub gene** |
| L1 | Nme9 | RV01 | Cnot1 |
| L2 | Tln1 | RV02 | LOC102923034 |
| L3 | LOC121830738 | RV03 | Dpy19l2 |
| L4 | Herc2 | RV04 | LOC102910557 |
| L5 | Fgr | RV05 | LOC121830936 |
| L6 | Zbtb17 | RV06 | Rps3 |
| L7 | LOC121824238 | RV07 | LOC102902778 |
| L8 | Cyfip2 | RV08 | Klhl40 |
| L9 | Rexo1 | RV09 | LOC102904132 |
| L10 | Ryr2 | RV10 | Coq10a |
| L11 | Dnaja1 | RV11 | Actn4 |
| L12 | Calhm6 | RV12 | Tob2 |
| L13 | Usp34 | RV13 | Per3 |
| L14 | LOC102925516 | RV14 | Mbd6 |
| L15 | Flt4 | RV15 | Otud6b |
| L16 | LOC102913793 | RV16 | Arntl |
| L17 | Cltc | RV17 | Pcdh1 |
| L18 | Fus | RV18 | Bag3 |
| L19 | Ccar1 | RV19 | Gapdh |
| L20 | Rps21 | RV20 | Fgb |
| L21 | Ntrk2 | RV21 | Ptges2 |
| L22 | Zbtb16 | RV22 | LOC121828149 |
| L23 | Ubl5 | RV23 | Vwc2 |
| L24 | Rab11b | RV24 | LOC121824812 |
| L25 | Elmo3 | RV25 | Clic1 |
| L26 | EI127_mgp04 | RV26 | C1qc |
| L27 | Esm1 | RV27 | Cd74 |
| L28 | LOC102914926 | RV28 | EI127_mgp08 |
| L29 | Tpm2 | RV29 | Lipe |
| L30 | Acp4 | RV30 | Acta2 |
| L31 | Epb41l5 | RV31 | Igfbp4 |
| L32 | Lamp3 | RV32 | Gnas |
| L33 | Cnbd2 | RV33 | Arap3 |
| L34 | Hipk3 | RV34 | Col3a1 |
| L35 | Hlf | RV35 | LOC102924108 |
| L36 | Tmx3 | RV36 | LOC102927417 |
| L37 | Arntl | RV37 | Mylpf |
| L38 | Bnc1 | | |
| L39 | Dip2a | | |

**Table S9.** Lung GO results of significant terms.

| Module | p_value | term_size | query_size | intersection_size | precision | recall | term_id | source | term_name |
|---|---|---|---|---|---|---|---|---|---|
| L3 | 0.008084442 | 4430 | 486 | 172 | 0.353909465 | 0.038826185 | GO:1901564 | GO:BP | organonitrogen compound metabolic process |
| L3 | 0.023714775 | 3073 | 486 | 126 | 0.259259259 | 0.041002278 | GO:0006810 | GO:BP | transport |
| L3 | 0.026331201 | 923 | 486 | 50 | 0.102880658 | 0.054171181 | GO:0006629 | GO:BP | lipid metabolic process |
| L3 | 4.44E-11 | 7299 | 469 | 289 | 0.616204691 | 0.039594465 | GO:0005737 | GO:CC | cytoplasm |
| L3 | 0.000385863 | 2533 | 469 | 112 | 0.23880597 | 0.044216344 | GO:0012505 | GO:CC | endomembrane system |
| L3 | 0.002536451 | 1013 | 469 | 54 | 0.115138593 | 0.053307009 | GO:0005783 | GO:CC | endoplasmic reticulum |
| L3 | 0.014864333 | 319 | 469 | 23 | 0.049040512 | 0.072100313 | GO:0000323 | GO:CC | lytic vacuole |
| L3 | 0.014864333 | 319 | 469 | 23 | 0.049040512 | 0.072100313 | GO:0005764 | GO:CC | lysosome |
| L3 | 0.049994563 | 11 | 469 | 4 | 0.008528785 | 0.363636364 | GO:0031464 | GO:CC | Cul4A-RING E3 ubiquitin ligase complex |
| L3 | 2.91E-06 | 5159 | 484 | 208 | 0.429752066 | 0.040317891 | GO:0003824 | GO:MF | catalytic activity |
| L3 | 0.000488832 | 72 | 484 | 12 | 0.024793388 | 0.166666667 | GO:0008135 | GO:MF | translation factor activity, RNA binding |
| L3 | 0.002312544 | 83 | 484 | 12 | 0.024793388 | 0.144578313 | GO:0090079 | GO:MF | translation regulator activity, nucleic acid binding |
| L3 | 0.018199614 | 3602 | 484 | 141 | 0.291322314 | 0.039144919 | GO:0043167 | GO:MF | ion binding |
| L3 | 0.019668132 | 102 | 484 | 12 | 0.024793388 | 0.117647059 | GO:0045182 | GO:MF | translation regulator activity |
| L7 | 9.89E-25 | 2754 | 557 | 195 | 0.350089767 | 0.0708061 | GO:0048731 | GO:BP | system development |
| L7 | 4.12E-24 | 5110 | 557 | 289 | 0.518850987 | 0.056555773 | GO:0032501 | GO:BP | multicellular organismal process |
| L7 | 3.28E-23 | 3317 | 557 | 216 | 0.387791741 | 0.065119084 | GO:0007275 | GO:BP | multicellular organism development |
| L7 | 2.54E-20 | 4205 | 557 | 245 | 0.439856373 | 0.058263971 | GO:0048856 | GO:BP | anatomical structure development |
| L7 | 4.03E-19 | 4492 | 557 | 253 | 0.454219031 | 0.056322351 | GO:0032502 | GO:BP | developmental process |
| L7 | 1.31E-18 | 1977 | 557 | 146 | 0.262118492 | 0.073849267 | GO:0009653 | GO:BP | anatomical structure morphogenesis |
| L7 | 6.47E-18 | 2183 | 557 | 154 | 0.276481149 | 0.070545121 | GO:0048513 | GO:BP | animal organ development |
| L7 | 2.52E-14 | 226 | 557 | 39 | 0.070017953 | 0.172566372 | GO:0030198 | GO:BP | extracellular matrix organization |
| L7 | 2.52E-14 | 226 | 557 | 39 | 0.070017953 | 0.172566372 | GO:0045229 | GO:BP | external encapsulating structure organization |
| L7 | 3.46E-14 | 228 | 557 | 39 | 0.070017953 | 0.171052632 | GO:0043062 | GO:BP | extracellular structure organization |
| L7 | 9.73E-13 | 853 | 557 | 77 | 0.138240575 | 0.090269637 | GO:0072359 | GO:BP | circulatory system development |
| L7 | 1.90E-12 | 2401 | 557 | 150 | 0.26929982 | 0.062473969 | GO:0023051 | GO:BP | regulation of signaling |
| L7 | 3.25E-12 | 2140 | 557 | 138 | 0.247755835 | 0.064485981 | GO:0051239 | GO:BP | regulation of multicellular organismal process |
| L7 | 4.74E-12 | 2403 | 557 | 149 | 0.267504488 | 0.062005826 | GO:0010646 | GO:BP | regulation of cell communication |
| L7 | 1.16E-11 | 1859 | 557 | 124 | 0.222621185 | 0.066702528 | GO:0007166 | GO:BP | cell surface receptor signaling pathway |
| L7 | 1.67E-11 | 4447 | 557 | 229 | 0.411131059 | 0.05149539 | GO:0048518 | GO:BP | positive regulation of biological process |
| L7 | 2.89E-11 | 908 | 557 | 77 | 0.138240575 | 0.084801762 | GO:0071495 | GO:BP | cellular response to endogenous stimulus |
| L7 | 3.25E-11 | 929 | 557 | 78 | 0.140035907 | 0.083961249 | GO:0007155 | GO:BP | cell adhesion |
| L7 | 4.10E-11 | 1520 | 557 | 107 | 0.192100539 | 0.070394737 | GO:0009888 | GO:BP | tissue development |
| L7 | 4.13E-11 | 2094 | 557 | 133 | 0.238779174 | 0.063514804 | GO:0009966 | GO:BP | regulation of signal transduction |
| L7 | 1.11E-10 | 2758 | 557 | 160 | 0.287253142 | 0.058013053 | GO:0048583 | GO:BP | regulation of response to stimulus |

| | | | | | | | | | |
|---|---|---|---|---|---|---|---|---|---|
| L7 | 1.21E-10 | 857 | 557 | 73 | 0.131059246 | 0.085180863 | GO:0035295 | GO:BP | tube development |
| L7 | 3.16E-10 | 418 | 557 | 47 | 0.08438061 | 0.112440191 | GO:0001501 | GO:BP | skeletal system development |
| L7 | 3.28E-10 | 990 | 557 | 79 | 0.141831239 | 0.07979798 | GO:0009719 | GO:BP | response to endogenous stimulus |
| L7 | 6.73E-10 | 1785 | 557 | 116 | 0.208258528 | 0.064985994 | GO:0050793 | GO:BP | regulation of developmental process |
| L7 | 1.02E-09 | 4050 | 557 | 208 | 0.373429084 | 0.051358025 | GO:0048522 | GO:BP | positive regulation of cellular process |
| L7 | 1.14E-09 | 820 | 557 | 69 | 0.123877917 | 0.084146341 | GO:0009887 | GO:BP | animal organ morphogenesis |
| L7 | 1.92E-09 | 906 | 557 | 73 | 0.131059246 | 0.080573951 | GO:0048646 | GO:BP | anatomical structure formation involved in morphogenesis |
| L7 | 6.36E-09 | 454 | 557 | 47 | 0.08438061 | 0.103524229 | GO:0071363 | GO:BP | cellular response to growth factor stimulus |
| L7 | 7.44E-09 | 1072 | 557 | 80 | 0.143626571 | 0.074626866 | GO:0016477 | GO:BP | cell migration |
| L7 | 1.86E-08 | 468 | 557 | 47 | 0.08438061 | 0.10042735 | GO:0070848 | GO:BP | response to growth factor |
| L7 | 2.04E-08 | 682 | 557 | 59 | 0.105924596 | 0.086510264 | GO:0035239 | GO:BP | tube morphogenesis |
| L7 | 2.84E-08 | 669 | 557 | 58 | 0.104129264 | 0.086696562 | GO:0030334 | GO:BP | regulation of cell migration |
| L7 | 2.99E-08 | 313 | 557 | 37 | 0.066427289 | 0.118210863 | GO:0008015 | GO:BP | blood circulation |
| L7 | 3.02E-08 | 1000 | 557 | 75 | 0.13464991 | 0.075 | GO:0007267 | GO:BP | cell-cell signaling |
| L7 | 3.85E-08 | 444 | 557 | 45 | 0.080789946 | 0.101351351 | GO:0007507 | GO:BP | heart development |
| L7 | 4.20E-08 | 332 | 557 | 38 | 0.068222621 | 0.114457831 | GO:0003013 | GO:BP | circulatory system process |
| L7 | 4.94E-08 | 551 | 557 | 51 | 0.091561939 | 0.092558984 | GO:0001568 | GO:BP | blood vessel development |
| L7 | 5.68E-08 | 260 | 557 | 33 | 0.059245961 | 0.126923077 | GO:0072001 | GO:BP | renal system development |
| L7 | 5.72E-08 | 2897 | 557 | 157 | 0.281867145 | 0.054193994 | GO:0030154 | GO:BP | cell differentiation |
| L7 | 5.72E-08 | 2897 | 557 | 157 | 0.281867145 | 0.054193994 | GO:0048869 | GO:BP | cellular developmental process |
| L7 | 8.18E-08 | 249 | 557 | 32 | 0.057450628 | 0.128514056 | GO:0001822 | GO:BP | kidney development |
| L7 | 1.01E-07 | 580 | 557 | 52 | 0.093357271 | 0.089655172 | GO:0001944 | GO:BP | vasculature development |
| L7 | 1.33E-07 | 870 | 557 | 67 | 0.120287253 | 0.077011494 | GO:0040011 | GO:BP | locomotion |
| L7 | 1.37E-07 | 715 | 557 | 59 | 0.105924596 | 0.082517483 | GO:2000145 | GO:BP | regulation of cell motility |
| L7 | 2.04E-07 | 303 | 557 | 35 | 0.062836625 | 0.115511551 | GO:0001503 | GO:BP | ossification |
| L7 | 3.23E-07 | 750 | 557 | 60 | 0.107719928 | 0.08 | GO:0040012 | GO:BP | regulation of locomotion |
| L7 | 3.77E-07 | 1577 | 557 | 99 | 0.177737882 | 0.062777425 | GO:0007399 | GO:BP | nervous system development |
| L7 | 4.11E-07 | 250 | 557 | 31 | 0.055655296 | 0.124 | GO:0003012 | GO:BP | muscle system process |
| L7 | 6.95E-07 | 9739 | 557 | 391 | 0.701974865 | 0.040147859 | GO:0065007 | GO:BP | biological regulation |
| L7 | 8.96E-07 | 352 | 557 | 37 | 0.066427289 | 0.105113636 | GO:0009611 | GO:BP | response to wounding |
| L7 | 9.71E-07 | 932 | 557 | 68 | 0.122082585 | 0.072961373 | GO:0051094 | GO:BP | positive regulation of developmental process |
| L7 | 1.52E-06 | 177 | 557 | 25 | 0.044883303 | 0.141242938 | GO:0030323 | GO:BP | respiratory tube development |
| L7 | 1.56E-06 | 515 | 557 | 46 | 0.082585278 | 0.089320388 | GO:0048598 | GO:BP | embryonic morphogenesis |
| L7 | 1.91E-06 | 1158 | 557 | 78 | 0.140035907 | 0.067357513 | GO:0022008 | GO:BP | neurogenesis |
| L7 | 1.94E-06 | 266 | 557 | 31 | 0.055655296 | 0.116541353 | GO:0042060 | GO:BP | wound healing |
| L7 | 2.71E-06 | 1232 | 557 | 81 | 0.145421903 | 0.065746753 | GO:0048870 | GO:BP | cell motility |
| L7 | 3.10E-06 | 197 | 557 | 26 | 0.046678636 | 0.131979695 | GO:0006936 | GO:BP | muscle contraction |
| L7 | 3.34E-06 | 1194 | 557 | 79 | 0.141831239 | 0.066164154 | GO:0048585 | GO:BP | negative regulation of response to stimulus |
| L7 | 4.29E-06 | 339 | 557 | 35 | 0.062836625 | 0.103244838 | GO:0044057 | GO:BP | regulation of system process |
| L7 | 5.09E-06 | 516 | 557 | 45 | 0.080789946 | 0.087209302 | GO:0048729 | GO:BP | tissue morphogenesis |
| L7 | 5.45E-06 | 122 | 557 | 20 | 0.035906643 | 0.163934426 | GO:0072006 | GO:BP | nephron development |

| | | | | | | | | | |
|---|---|---|---|---|---|---|---|---|---|
| L7 | 8.11E-06 | 1810 | 557 | 105 | 0.188509874 | 0.05801105 | GO:0065008 | GO:BP | regulation of biological quality |
| L7 | 8.18E-06 | 1196 | 557 | 78 | 0.140035907 | 0.065217391 | GO:0003008 | GO:BP | system process |
| L7 | 8.55E-06 | 178 | 557 | 24 | 0.043087971 | 0.134831461 | GO:0060348 | GO:BP | bone development |
| L7 | 8.65E-06 | 581 | 557 | 48 | 0.086175943 | 0.082616179 | GO:0022603 | GO:BP | regulation of anatomical structure morphogenesis |
| L7 | 1.31E-05 | 2162 | 557 | 119 | 0.213644524 | 0.055041628 | GO:0042221 | GO:BP | response to chemical |
| L7 | 1.31E-05 | 5276 | 557 | 237 | 0.425493716 | 0.044920394 | GO:0023052 | GO:BP | signaling |
| L7 | 1.45E-05 | 5366 | 557 | 240 | 0.430879713 | 0.044726053 | GO:0007154 | GO:BP | cell communication |
| L7 | 1.71E-05 | 6965 | 557 | 295 | 0.52962298 | 0.04235463 | GO:0050896 | GO:BP | response to stimulus |
| L7 | 1.99E-05 | 1699 | 557 | 99 | 0.177737882 | 0.05826957 | GO:0051128 | GO:BP | regulation of cellular component organization |
| L7 | 2.20E-05 | 431 | 557 | 39 | 0.070017953 | 0.090487239 | GO:0002009 | GO:BP | morphogenesis of an epithelium |
| L7 | 2.27E-05 | 1158 | 557 | 75 | 0.13464991 | 0.064766839 | GO:0051240 | GO:BP | positive regulation of multicellular organismal process |
| L7 | 2.35E-05 | 173 | 557 | 23 | 0.041292639 | 0.132947977 | GO:0071560 | GO:BP | cellular response to transforming growth factor beta stimulus |
| L7 | 2.49E-05 | 1095 | 557 | 72 | 0.129263914 | 0.065753425 | GO:0009967 | GO:BP | positive regulation of signal transduction |
| L7 | 2.63E-05 | 174 | 557 | 23 | 0.041292639 | 0.132183908 | GO:0030324 | GO:BP | lung development |
| L7 | 2.73E-05 | 54 | 557 | 13 | 0.023339318 | 0.240740741 | GO:0032963 | GO:BP | collagen metabolic process |
| L7 | 3.15E-05 | 190 | 557 | 24 | 0.043087971 | 0.126315789 | GO:0001763 | GO:BP | morphogenesis of a branching structure |
| L7 | 3.64E-05 | 177 | 557 | 23 | 0.041292639 | 0.129943503 | GO:0071559 | GO:BP | response to transforming growth factor beta |
| L7 | 3.98E-05 | 9499 | 557 | 376 | 0.675044883 | 0.039583114 | GO:0050789 | GO:BP | regulation of biological process |
| L7 | 4.50E-05 | 691 | 557 | 52 | 0.093357271 | 0.075253256 | GO:0007167 | GO:BP | enzyme-linked receptor protein signaling pathway |
| L7 | 4.74E-05 | 372 | 557 | 35 | 0.062836625 | 0.094086022 | GO:0048667 | GO:BP | cell morphogenesis involved in neuron differentiation |
| L7 | 4.79E-05 | 138 | 557 | 20 | 0.035906643 | 0.144927536 | GO:0035108 | GO:BP | limb morphogenesis |
| L7 | 4.79E-05 | 138 | 557 | 20 | 0.035906643 | 0.144927536 | GO:0035107 | GO:BP | appendage morphogenesis |
| L7 | 4.79E-05 | 47 | 557 | 12 | 0.021543986 | 0.255319149 | GO:0030199 | GO:BP | collagen fibril organization |
| L7 | 5.10E-05 | 1225 | 557 | 77 | 0.138240575 | 0.062857143 | GO:0141124 | GO:BP | intracellular signaling cassette |
| L7 | 5.16E-05 | 500 | 557 | 42 | 0.07540395 | 0.084 | GO:0034330 | GO:BP | cell junction organization |
| L7 | 5.52E-05 | 257 | 557 | 28 | 0.0502693 | 0.108949416 | GO:0031589 | GO:BP | cell-substrate adhesion |
| L7 | 5.76E-05 | 1009 | 557 | 67 | 0.120287253 | 0.066402379 | GO:2000026 | GO:BP | regulation of multicellular organismal development |
| L7 | 5.79E-05 | 196 | 557 | 24 | 0.043087971 | 0.12244898 | GO:0060541 | GO:BP | respiratory system development |
| L7 | 6.27E-05 | 1231 | 557 | 77 | 0.138240575 | 0.062550772 | GO:0010647 | GO:BP | positive regulation of cell communication |
| L7 | 6.72E-05 | 620 | 557 | 48 | 0.086175943 | 0.077419355 | GO:0031175 | GO:BP | neuron projection development |
| L7 | 7.44E-05 | 1236 | 557 | 77 | 0.138240575 | 0.062297735 | GO:0023056 | GO:BP | positive regulation of signaling |
| L7 | 7.81E-05 | 470 | 557 | 40 | 0.071813285 | 0.085106383 | GO:0048514 | GO:BP | blood vessel morphogenesis |
| L7 | 8.09E-05 | 932 | 557 | 63 | 0.113105925 | 0.067596567 | GO:0009790 | GO:BP | embryo development |
| L7 | 8.18E-05 | 129 | 557 | 19 | 0.034111311 | 0.147286822 | GO:0051897 | GO:BP | positive regulation of phosphatidylinositol 3-kinase/protein kinase B signal transduction |
| L7 | 9.57E-05 | 1682 | 557 | 96 | 0.172351885 | 0.057074911 | GO:0070887 | GO:BP | cellular response to chemical stimulus |
| L7 | 0.000119485 | 1572 | 557 | 91 | 0.163375224 | 0.057888041 | GO:0010033 | GO:BP | response to organic substance |
| L7 | 0.000121392 | 652 | 557 | 49 | 0.087971275 | 0.075153374 | GO:0000902 | GO:BP | cell morphogenesis |
| L7 | 0.000165754 | 177 | 557 | 22 | 0.039497307 | 0.124293785 | GO:0061138 | GO:BP | morphogenesis of a branching epithelium |
| L7 | 0.000208078 | 413 | 557 | 36 | 0.064631957 | 0.08716707 | GO:0040017 | GO:BP | positive regulation of locomotion |
| L7 | 0.000219727 | 1593 | 557 | 91 | 0.163375224 | 0.057124922 | GO:0048584 | GO:BP | positive regulation of response to stimulus |
| L7 | 0.000232341 | 44 | 557 | 11 | 0.019748654 | 0.25 | GO:0032330 | GO:BP | regulation of chondrocyte differentiation |

| | | | | | | | | | |
|---|---|---|---|---|---|---|---|---|---|
| L7 | 0.000265631 | 167 | 557 | 21 | 0.037701975 | 0.125748503 | GO:0060173 | GO:BP | limb development |
| L7 | 0.000265631 | 167 | 557 | 21 | 0.037701975 | 0.125748503 | GO:0048736 | GO:BP | appendage development |
| L7 | 0.000270658 | 942 | 557 | 62 | 0.111310592 | 0.06581741 | GO:0060429 | GO:BP | epithelium development |
| L7 | 0.000272671 | 5965 | 557 | 255 | 0.457809695 | 0.042749371 | GO:0051716 | GO:BP | cellular response to stimulus |
| L7 | 0.000287214 | 382 | 557 | 34 | 0.061041293 | 0.089005236 | GO:0030335 | GO:BP | positive regulation of cell migration |
| L7 | 0.00029183 | 229 | 557 | 25 | 0.044883303 | 0.109170306 | GO:0090287 | GO:BP | regulation of cellular response to growth factor stimulus |
| L7 | 0.000299072 | 1010 | 557 | 65 | 0.116696589 | 0.064356436 | GO:0023057 | GO:BP | negative regulation of signaling |
| L7 | 0.000319958 | 990 | 557 | 64 | 0.114901257 | 0.064646465 | GO:0048699 | GO:BP | generation of neurons |
| L7 | 0.000333848 | 926 | 557 | 61 | 0.10951526 | 0.06587473 | GO:0030182 | GO:BP | neuron differentiation |
| L7 | 0.000347713 | 141 | 557 | 19 | 0.034111311 | 0.134751773 | GO:0090257 | GO:BP | regulation of muscle system process |
| L7 | 0.000359997 | 800 | 557 | 55 | 0.098743268 | 0.06875 | GO:0051241 | GO:BP | negative regulation of multicellular organismal process |
| L7 | 0.000379163 | 128 | 557 | 18 | 0.032315978 | 0.140625 | GO:0003018 | GO:BP | vascular process in circulatory system |
| L7 | 0.000540973 | 131 | 557 | 18 | 0.032315978 | 0.13740458 | GO:0031214 | GO:BP | biomineral tissue development |
| L7 | 0.000547886 | 411 | 557 | 35 | 0.062836625 | 0.085158151 | GO:0048858 | GO:BP | cell projection morphogenesis |
| L7 | 0.000586127 | 1007 | 557 | 64 | 0.114901257 | 0.063555114 | GO:0010648 | GO:BP | negative regulation of cell communication |
| L7 | 0.000586242 | 322 | 557 | 30 | 0.053859964 | 0.093167702 | GO:0198738 | GO:BP | cell-cell signaling by wnt |
| L7 | 0.000605376 | 146 | 557 | 19 | 0.034111311 | 0.130136986 | GO:0048754 | GO:BP | branching morphogenesis of an epithelial tube |
| L7 | 0.000631679 | 395 | 557 | 34 | 0.061041293 | 0.086075949 | GO:0048812 | GO:BP | neuron projection morphogenesis |
| L7 | 0.000715909 | 59 | 557 | 12 | 0.021543986 | 0.203389831 | GO:0061035 | GO:BP | regulation of cartilage development |
| L7 | 0.000877179 | 1108 | 557 | 68 | 0.122082585 | 0.061371841 | GO:0045595 | GO:BP | regulation of cell differentiation |
| L7 | 0.00094397 | 3938 | 557 | 180 | 0.323159785 | 0.045708481 | GO:0048519 | GO:BP | negative regulation of biological process |
| L7 | 0.000949498 | 402 | 557 | 34 | 0.061041293 | 0.084577114 | GO:2000147 | GO:BP | positive regulation of cell motility |
| L7 | 0.000955921 | 165 | 557 | 20 | 0.035906643 | 0.121212121 | GO:0003015 | GO:BP | heart process |
| L7 | 0.000961423 | 9049 | 557 | 355 | 0.637342908 | 0.039230854 | GO:0050794 | GO:BP | regulation of cellular process |
| L7 | 0.000979629 | 244 | 557 | 25 | 0.044883303 | 0.102459016 | GO:0030111 | GO:BP | regulation of Wnt signaling pathway |
| L7 | 0.001097567 | 24 | 557 | 8 | 0.014362657 | 0.333333333 | GO:0032964 | GO:BP | collagen biosynthetic process |
| L7 | 0.001152441 | 787 | 557 | 53 | 0.095152603 | 0.067344346 | GO:0051130 | GO:BP | positive regulation of cellular component organization |
| L7 | 0.001287906 | 98 | 557 | 15 | 0.026929982 | 0.153061224 | GO:0072009 | GO:BP | nephron epithelium development |
| L7 | 0.001334487 | 408 | 557 | 34 | 0.061041293 | 0.083333333 | GO:0120039 | GO:BP | plasma membrane bounded cell projection morphogenesis |
| L7 | 0.001406165 | 1055 | 557 | 65 | 0.116696589 | 0.061611374 | GO:0030030 | GO:BP | cell projection organization |
| L7 | 0.001494917 | 301 | 557 | 28 | 0.0502693 | 0.093023256 | GO:0061564 | GO:BP | axon development |
| L7 | 0.001631941 | 320 | 557 | 29 | 0.052064632 | 0.090625 | GO:0016055 | GO:BP | Wnt signaling pathway |
| L7 | 0.001678055 | 949 | 557 | 60 | 0.107719928 | 0.063224447 | GO:0009968 | GO:BP | negative regulation of signal transduction |
| L7 | 0.001821388 | 2014 | 557 | 105 | 0.188509874 | 0.052135055 | GO:0048468 | GO:BP | cell development |
| L7 | 0.001879508 | 157 | 557 | 19 | 0.034111311 | 0.121019108 | GO:0060047 | GO:BP | heart contraction |
| L7 | 0.002211302 | 740 | 557 | 50 | 0.089766607 | 0.067567568 | GO:0048666 | GO:BP | neuron development |
| L7 | 0.002282354 | 159 | 557 | 19 | 0.034111311 | 0.119496855 | GO:1903522 | GO:BP | regulation of blood circulation |
| L7 | 0.002391907 | 256 | 557 | 25 | 0.044883303 | 0.09765625 | GO:0141091 | GO:BP | transforming growth factor beta receptor superfamily signaling pathway |
| L7 | 0.002466795 | 291 | 557 | 27 | 0.048473968 | 0.092783505 | GO:0060562 | GO:BP | epithelial tube morphogenesis |
| L7 | 0.002472683 | 103 | 557 | 15 | 0.026929982 | 0.145631068 | GO:0030282 | GO:BP | bone mineralization |
| L7 | 0.002481377 | 145 | 557 | 18 | 0.032315978 | 0.124137931 | GO:0007179 | GO:BP | transforming growth factor beta receptor signaling pathway |

| | | | | | | | | | |
|---|---|---|---|---|---|---|---|---|---|
| L7 | 0.003035481 | 27 | 557 | 8 | 0.014362657 | 0.296296296 | GO:0048566 | GO:BP | embryonic digestive tract development |
| L7 | 0.003152112 | 482 | 557 | 37 | 0.066427289 | 0.076763485 | GO:0032989 | GO:BP | cellular anatomical entity morphogenesis |
| L7 | 0.003526422 | 4931 | 557 | 213 | 0.382405745 | 0.043196106 | GO:0007165 | GO:BP | signal transduction |
| L7 | 0.004085456 | 81 | 557 | 13 | 0.023339318 | 0.160493827 | GO:0072080 | GO:BP | nephron tubule development |
| L7 | 0.004151746 | 568 | 557 | 41 | 0.073608618 | 0.072183099 | GO:1901698 | GO:BP | response to nitrogen compound |
| L7 | 0.004216646 | 94 | 557 | 14 | 0.02513465 | 0.14893617 | GO:0097746 | GO:BP | blood vessel diameter maintenance |
| L7 | 0.004216646 | 94 | 557 | 14 | 0.02513465 | 0.14893617 | GO:0035296 | GO:BP | regulation of tube diameter |
| L7 | 0.004351272 | 1045 | 557 | 63 | 0.113105925 | 0.060287081 | GO:0120036 | GO:BP | plasma membrane bounded cell projection organization |
| L7 | 0.00479815 | 95 | 557 | 14 | 0.02513465 | 0.147368421 | GO:0035150 | GO:BP | regulation of tube size |
| L7 | 0.005285832 | 285 | 557 | 26 | 0.046678636 | 0.09122807 | GO:0007178 | GO:BP | transmembrane receptor protein serine/threonine kinase signaling pathway |
| L7 | 0.00530848 | 48 | 557 | 10 | 0.017953321 | 0.208333333 | GO:0060688 | GO:BP | regulation of morphogenesis of a branching structure |
| L7 | 0.005364896 | 38 | 557 | 9 | 0.016157989 | 0.236842105 | GO:0085029 | GO:BP | extracellular matrix assembly |
| L7 | 0.005365234 | 14 | 557 | 6 | 0.010771993 | 0.428571429 | GO:0019934 | GO:BP | cGMP-mediated signaling |
| L7 | 0.005758578 | 110 | 557 | 15 | 0.026929982 | 0.136363636 | GO:0048706 | GO:BP | embryonic skeletal system development |
| L7 | 0.005842397 | 185 | 557 | 20 | 0.035906643 | 0.108108108 | GO:0051896 | GO:BP | regulation of phosphatidylinositol 3-kinase/protein kinase B signal transduction |
| L7 | 0.006351272 | 125 | 557 | 16 | 0.028725314 | 0.128 | GO:0072073 | GO:BP | kidney epithelium development |
| L7 | 0.006363341 | 537 | 557 | 39 | 0.070017953 | 0.072625698 | GO:0060322 | GO:BP | head development |
| L7 | 0.006435867 | 60 | 557 | 11 | 0.019748654 | 0.183333333 | GO:0060675 | GO:BP | ureteric bud morphogenesis |
| L7 | 0.006542144 | 1243 | 557 | 71 | 0.127468582 | 0.057119871 | GO:0071310 | GO:BP | cellular response to organic substance |
| L7 | 0.006772157 | 39 | 557 | 9 | 0.016157989 | 0.230769231 | GO:0048146 | GO:BP | positive regulation of fibroblast proliferation |
| L7 | 0.006836929 | 171 | 557 | 19 | 0.034111311 | 0.111111111 | GO:0051216 | GO:BP | cartilage development |
| L7 | 0.007271066 | 141 | 557 | 17 | 0.030520646 | 0.120567376 | GO:0003205 | GO:BP | cardiac chamber development |
| L7 | 0.007513771 | 1704 | 557 | 90 | 0.161579892 | 0.052816901 | GO:0009605 | GO:BP | response to external stimulus |
| L7 | 0.007620966 | 61 | 557 | 11 | 0.019748654 | 0.180327869 | GO:0072171 | GO:BP | mesonephric tubule morphogenesis |
| L7 | 0.007732579 | 9 | 557 | 5 | 0.008976661 | 0.555555556 | GO:0048251 | GO:BP | elastic fiber assembly |
| L7 | 0.007827789 | 709 | 557 | 47 | 0.08438061 | 0.06629055 | GO:1901701 | GO:BP | cellular response to oxygen-containing compound |
| L7 | 0.007904567 | 99 | 557 | 14 | 0.02513465 | 0.141414141 | GO:0002062 | GO:BP | chondrocyte differentiation |
| L7 | 0.008086033 | 222 | 557 | 22 | 0.039497307 | 0.099099099 | GO:0043491 | GO:BP | phosphatidylinositol 3-kinase/protein kinase B signal transduction |
| L7 | 0.008148151 | 86 | 557 | 13 | 0.023339318 | 0.151162791 | GO:0061326 | GO:BP | renal tubule development |
| L7 | 0.008222847 | 1417 | 557 | 78 | 0.140035907 | 0.055045872 | GO:0008283 | GO:BP | cell population proliferation |
| L7 | 0.008795951 | 524 | 557 | 38 | 0.068222621 | 0.072519084 | GO:0030029 | GO:BP | actin filament-based process |
| L7 | 0.009201235 | 207 | 557 | 21 | 0.037701975 | 0.101449275 | GO:0003007 | GO:BP | heart morphogenesis |
| L7 | 0.009310743 | 387 | 557 | 31 | 0.055655296 | 0.080103359 | GO:0001525 | GO:BP | angiogenesis |
| L7 | 0.009356174 | 224 | 557 | 22 | 0.039497307 | 0.098214286 | GO:0060485 | GO:BP | mesenchyme development |
| L7 | 0.009454734 | 934 | 557 | 57 | 0.102333932 | 0.061027837 | GO:1901700 | GO:BP | response to oxygen-containing compound |
| L7 | 0.009957443 | 427 | 557 | 33 | 0.059245961 | 0.077283372 | GO:0007169 | GO:BP | transmembrane receptor protein tyrosine kinase signaling pathway |
| L7 | 0.010542165 | 278 | 557 | 25 | 0.044883303 | 0.089928058 | GO:0007409 | GO:BP | axonogenesis |
| L7 | 0.010550736 | 717 | 557 | 47 | 0.08438061 | 0.065550907 | GO:0044087 | GO:BP | regulation of cellular component biogenesis |
| L7 | 0.011305985 | 23 | 557 | 7 | 0.012567325 | 0.304347826 | GO:0032331 | GO:BP | negative regulation of chondrocyte differentiation |
| L7 | 0.012977285 | 132 | 557 | 16 | 0.028725314 | 0.121212121 | GO:0030178 | GO:BP | negative regulation of Wnt signaling pathway |
| L7 | 0.012979611 | 595 | 557 | 41 | 0.073608618 | 0.068907563 | GO:0045597 | GO:BP | positive regulation of cell differentiation |

| | | | | | | | | | |
|---|---|---|---|---|---|---|---|---|---|
| L7 | 0.013521594 | 16 | 557 | 6 | 0.010771993 | 0.375 | GO:0045932 | GO:BP | negative regulation of muscle contraction |
| L7 | 0.013521594 | 16 | 557 | 6 | 0.010771993 | 0.375 | GO:0007263 | GO:BP | nitric oxide mediated signal transduction |
| L7 | 0.013645807 | 53 | 557 | 10 | 0.017953321 | 0.188679245 | GO:0001658 | GO:BP | branching involved in ureteric bud morphogenesis |
| L7 | 0.013692475 | 77 | 557 | 12 | 0.021543986 | 0.155844156 | GO:0070167 | GO:BP | regulation of biomineral tissue development |
| L7 | 0.013692475 | 77 | 557 | 12 | 0.021543986 | 0.155844156 | GO:0060021 | GO:BP | roof of mouth development |
| L7 | 0.01422135 | 104 | 557 | 14 | 0.02513465 | 0.134615385 | GO:0006937 | GO:BP | regulation of muscle contraction |
| L7 | 0.015045015 | 10 | 557 | 5 | 0.008976661 | 0.5 | GO:0010749 | GO:BP | regulation of nitric oxide mediated signal transduction |
| L7 | 0.015045015 | 10 | 557 | 5 | 0.008976661 | 0.5 | GO:0045986 | GO:BP | negative regulation of smooth muscle contraction |
| L7 | 0.015317136 | 284 | 557 | 25 | 0.044883303 | 0.088028169 | GO:0034329 | GO:BP | cell junction assembly |
| L7 | 0.015769712 | 3696 | 557 | 165 | 0.296229803 | 0.044642857 | GO:0048523 | GO:BP | negative regulation of cellular process |
| L7 | 0.017449434 | 92 | 557 | 13 | 0.023339318 | 0.141304348 | GO:0001823 | GO:BP | mesonephros development |
| L7 | 0.017554847 | 380 | 557 | 30 | 0.053859964 | 0.078947368 | GO:0048568 | GO:BP | embryonic organ development |
| L7 | 0.018304144 | 646 | 557 | 43 | 0.077199282 | 0.066563467 | GO:0040007 | GO:BP | growth |
| L7 | 0.018864579 | 234 | 557 | 22 | 0.039497307 | 0.094017094 | GO:0060070 | GO:BP | canonical Wnt signaling pathway |
| L7 | 0.019011564 | 647 | 557 | 43 | 0.077199282 | 0.066460587 | GO:0008284 | GO:BP | positive regulation of cell population proliferation |
| L7 | 0.019444869 | 252 | 557 | 23 | 0.041292639 | 0.091269841 | GO:0010975 | GO:BP | regulation of neuron projection development |
| L7 | 0.019587466 | 67 | 557 | 11 | 0.019748654 | 0.164179104 | GO:0072078 | GO:BP | nephron tubule morphogenesis |
| L7 | 0.020933696 | 671 | 557 | 44 | 0.078994614 | 0.06557377 | GO:0007417 | GO:BP | central nervous system development |
| L7 | 0.021825165 | 1850 | 557 | 94 | 0.168761221 | 0.050810811 | GO:0035556 | GO:BP | intracellular signal transduction |
| L7 | 0.023376113 | 81 | 557 | 12 | 0.021543986 | 0.148148148 | GO:2000027 | GO:BP | regulation of animal organ morphogenesis |
| L7 | 0.023786732 | 203 | 557 | 20 | 0.035906643 | 0.098522167 | GO:0048705 | GO:BP | skeletal system morphogenesis |
| L7 | 0.025221192 | 1151 | 557 | 65 | 0.116696589 | 0.056472632 | GO:0042127 | GO:BP | regulation of cell population proliferation |
| L7 | 0.026191607 | 69 | 557 | 11 | 0.019748654 | 0.15942029 | GO:0061333 | GO:BP | renal tubule morphogenesis |
| L7 | 0.026191607 | 69 | 557 | 11 | 0.019748654 | 0.15942029 | GO:0072088 | GO:BP | nephron epithelium morphogenesis |
| L7 | 0.027832518 | 26 | 557 | 7 | 0.012567325 | 0.269230769 | GO:0052652 | GO:BP | cyclic purine nucleotide metabolic process |
| L7 | 0.028350103 | 276 | 557 | 24 | 0.043087971 | 0.086956522 | GO:0032102 | GO:BP | negative regulation of response to external stimulus |
| L7 | 0.029398284 | 511 | 557 | 36 | 0.064631957 | 0.070450098 | GO:0010243 | GO:BP | response to organonitrogen compound |
| L7 | 0.029759826 | 6 | 557 | 4 | 0.007181329 | 0.666666667 | GO:0060831 | GO:BP | smoothened signaling pathway involved in dorsal/ventral neural tube patterning |
| L7 | 0.030016032 | 241 | 557 | 22 | 0.039497307 | 0.091286307 | GO:0010631 | GO:BP | epithelial cell migration |
| L7 | 0.030312024 | 333 | 557 | 27 | 0.048473968 | 0.081081081 | GO:0001667 | GO:BP | ameboidal-type cell migration |
| L7 | 0.033309235 | 225 | 557 | 21 | 0.037701975 | 0.093333333 | GO:0061448 | GO:BP | connective tissue development |
| L7 | 0.033416238 | 1185 | 557 | 66 | 0.118491921 | 0.055696203 | GO:0051246 | GO:BP | regulation of protein metabolic process |
| L7 | 0.033577446 | 393 | 557 | 30 | 0.053859964 | 0.076335878 | GO:1901699 | GO:BP | cellular response to nitrogen compound |
| L7 | 0.034146322 | 243 | 557 | 22 | 0.039497307 | 0.090534979 | GO:0090132 | GO:BP | epithelium migration |
| L7 | 0.034646203 | 71 | 557 | 11 | 0.019748654 | 0.154929577 | GO:0006939 | GO:BP | smooth muscle contraction |
| L7 | 0.034646203 | 71 | 557 | 11 | 0.019748654 | 0.154929577 | GO:0072028 | GO:BP | nephron morphogenesis |
| L7 | 0.036511481 | 27 | 557 | 7 | 0.012567325 | 0.259259259 | GO:0030511 | GO:BP | positive regulation of transforming growth factor beta receptor signaling pathway |
| L7 | 0.036511481 | 27 | 557 | 7 | 0.012567325 | 0.259259259 | GO:0061037 | GO:BP | negative regulation of cartilage development |
| L7 | 0.036511481 | 27 | 557 | 7 | 0.012567325 | 0.259259259 | GO:1903846 | GO:BP | positive regulation of cellular response to transforming growth factor beta stimulus |
| L7 | 0.038579899 | 85 | 557 | 12 | 0.021543986 | 0.141176471 | GO:0060993 | GO:BP | kidney morphogenesis |
| L7 | 0.038781889 | 245 | 557 | 22 | 0.039497307 | 0.089795918 | GO:0090130 | GO:BP | tissue migration |

| | | | | | | | | | |
|---|---|---|---|---|---|---|---|---|---|
| L7 | 0.04134918 | 114 | 557 | 14 | 0.02513465 | 0.122807018 | GO:0030326 | GO:BP | embryonic limb morphogenesis |
| L7 | 0.04134918 | 114 | 557 | 14 | 0.02513465 | 0.122807018 | GO:0035113 | GO:BP | embryonic appendage morphogenesis |
| L7 | 0.041606003 | 177 | 557 | 18 | 0.032315978 | 0.101694915 | GO:0010632 | GO:BP | regulation of epithelial cell migration |
| L7 | 0.042934774 | 161 | 557 | 17 | 0.030520646 | 0.105590062 | GO:0001649 | GO:BP | osteoblast differentiation |
| L7 | 0.044037414 | 500 | 557 | 35 | 0.062836625 | 0.07 | GO:0097435 | GO:BP | supramolecular fiber organization |
| L7 | 0.04475029 | 399 | 557 | 30 | 0.053859964 | 0.07518797 | GO:0120035 | GO:BP | regulation of plasma membrane bounded cell projection organization |
| L7 | 0.04855164 | 38 | 557 | 8 | 0.014362657 | 0.210526316 | GO:0006940 | GO:BP | regulation of smooth muscle contraction |
| L7 | 0.04883082 | 196 | 557 | 19 | 0.034111311 | 0.096938776 | GO:0090092 | GO:BP | regulation of transmembrane receptor protein serine/threonine kinase signaling pathway |
| L7 | 0.049510485 | 131 | 557 | 15 | 0.026929982 | 0.114503817 | GO:0007596 | GO:BP | blood coagulation |
| L7 | 2.16E-36 | 3202 | 532 | 235 | 0.441729323 | 0.07339163 | GO:0071944 | GO:CC | cell periphery |
| L7 | 8.26E-30 | 228 | 532 | 55 | 0.103383459 | 0.24122807 | GO:0031012 | GO:CC | extracellular matrix |
| L7 | 1.06E-29 | 229 | 532 | 55 | 0.103383459 | 0.240174672 | GO:0030312 | GO:CC | external encapsulating structure |
| L7 | 2.59E-20 | 2917 | 532 | 189 | 0.355263158 | 0.064792595 | GO:0005886 | GO:CC | plasma membrane |
| L7 | 3.56E-18 | 138 | 532 | 34 | 0.063909774 | 0.246376812 | GO:0062023 | GO:CC | collagen-containing extracellular matrix |
| L7 | 1.65E-08 | 1116 | 532 | 79 | 0.148496241 | 0.07078853 | GO:0005576 | GO:CC | extracellular region |
| L7 | 1.17E-07 | 692 | 532 | 56 | 0.105263158 | 0.080924855 | GO:0005615 | GO:CC | extracellular space |
| L7 | 1.62E-07 | 1214 | 532 | 81 | 0.152255639 | 0.066721582 | GO:0030054 | GO:CC | cell junction |
| L7 | 8.12E-07 | 7299 | 532 | 306 | 0.57518797 | 0.041923551 | GO:0005737 | GO:CC | cytoplasm |
| L7 | 1.50E-06 | 69 | 532 | 15 | 0.028195489 | 0.217391304 | GO:0042383 | GO:CC | sarcolemma |
| L7 | 2.31E-06 | 61 | 532 | 14 | 0.026315789 | 0.229508197 | GO:0005604 | GO:CC | basement membrane |
| L7 | 7.86E-05 | 6409 | 532 | 267 | 0.501879699 | 0.041660165 | GO:0016020 | GO:CC | membrane |
| L7 | 0.000222167 | 7 | 532 | 5 | 0.009398496 | 0.714285714 | GO:0001527 | GO:CC | microfibril |
| L7 | 0.002455598 | 10 | 532 | 5 | 0.009398496 | 0.5 | GO:0005583 | GO:CC | fibrillar collagen trimer |
| L7 | 0.002455598 | 10 | 532 | 5 | 0.009398496 | 0.5 | GO:0098643 | GO:CC | banded collagen fibril |
| L7 | 0.003677903 | 742 | 532 | 47 | 0.088345865 | 0.063342318 | GO:0045202 | GO:CC | synapse |
| L7 | 0.00962396 | 2533 | 532 | 118 | 0.221804511 | 0.046585077 | GO:0012505 | GO:CC | endomembrane system |
| L7 | 0.011096622 | 7 | 532 | 4 | 0.007518797 | 0.571428571 | GO:1990454 | GO:CC | L-type voltage-gated calcium channel complex |
| L7 | 0.012096434 | 892 | 532 | 52 | 0.097744361 | 0.058295964 | GO:0005794 | GO:CC | Golgi apparatus |
| L7 | 0.013482984 | 170 | 532 | 17 | 0.031954887 | 0.1 | GO:0098984 | GO:CC | neuron to neuron synapse |
| L7 | 0.017296703 | 22 | 532 | 6 | 0.011278195 | 0.272727273 | GO:0005581 | GO:CC | collagen trimer |
| L7 | 0.03613119 | 16 | 532 | 5 | 0.009398496 | 0.3125 | GO:0098644 | GO:CC | complex of collagen trimers |
| L7 | 0.041035952 | 4 | 532 | 3 | 0.005639098 | 0.75 | GO:0071953 | GO:CC | elastic fiber |
| L7 | 0.047731175 | 26 | 532 | 6 | 0.011278195 | 0.230769231 | GO:0030315 | GO:CC | T-tubule |
| L7 | 1.03E-15 | 7099 | 563 | 339 | 0.602131439 | 0.047753205 | GO:0005515 | GO:MF | protein binding |
| L7 | 1.08E-06 | 12006 | 563 | 462 | 0.820603908 | 0.03848076 | GO:0005488 | GO:MF | binding |
| L7 | 2.71E-06 | 50 | 563 | 13 | 0.023090586 | 0.26 | GO:0005518 | GO:MF | collagen binding |
| L7 | 5.44E-06 | 11 | 563 | 7 | 0.012433393 | 0.636363636 | GO:0048407 | GO:MF | platelet-derived growth factor binding |
| L7 | 2.28E-05 | 1007 | 563 | 67 | 0.119005329 | 0.06653426 | GO:0005102 | GO:MF | signaling receptor binding |
| L7 | 2.76E-05 | 1455 | 563 | 87 | 0.154529307 | 0.059793814 | GO:0019899 | GO:MF | enzyme binding |
| L7 | 3.79E-05 | 83 | 563 | 15 | 0.026642984 | 0.180722892 | GO:0008201 | GO:MF | heparin binding |
| L7 | 0.000109122 | 128 | 563 | 18 | 0.031971581 | 0.140625 | GO:0005539 | GO:MF | glycosaminoglycan binding |

| | | | | | | | | | |
|---|---|---|---|---|---|---|---|---|---|
| L7 | 0.000285522 | 32 | 563 | 9 | 0.01598579 | 0.28125 | GO:0005201 | GO:MF | extracellular matrix structural constituent |
| L7 | 0.00044096 | 582 | 563 | 43 | 0.076376554 | 0.073883162 | GO:0005509 | GO:MF | calcium ion binding |
| L7 | 0.000504781 | 4 | 563 | 4 | 0.007104796 | 1 | GO:0086007 | GO:MF | voltage-gated calcium channel activity involved in cardiac muscle cell action potential |
| L7 | 0.001050179 | 164 | 563 | 19 | 0.03374778 | 0.115853659 | GO:0050839 | GO:MF | cell adhesion molecule binding |
| L7 | 0.001242427 | 136 | 563 | 17 | 0.030195382 | 0.125 | GO:0005096 | GO:MF | GTPase activator activity |
| L7 | 0.006969994 | 139 | 563 | 16 | 0.028419183 | 0.115107914 | GO:1901681 | GO:MF | sulfur compound binding |
| L7 | 0.011712974 | 100 | 563 | 13 | 0.023090586 | 0.13 | GO:0019838 | GO:MF | growth factor binding |
| L7 | 0.012473622 | 320 | 563 | 26 | 0.046181172 | 0.08125 | GO:0060589 | GO:MF | nucleoside-triphosphatase regulator activity |
| L7 | 0.012473622 | 320 | 563 | 26 | 0.046181172 | 0.08125 | GO:0030695 | GO:MF | GTPase regulator activity |
| L7 | 0.012997974 | 87 | 563 | 12 | 0.021314387 | 0.137931034 | GO:0005178 | GO:MF | integrin binding |
| L7 | 0.042621847 | 98 | 563 | 12 | 0.021314387 | 0.12244898 | GO:0015085 | GO:MF | calcium ion transmembrane transporter activity |
| L7 | 1.02E-05 | 108 | 223 | 21 | 0.094170404 | 0.194444444 | HP:0100699 | HP | Scarring |
| L7 | 1.19E-05 | 6 | 223 | 6 | 0.02690583 | 1 | HP:0010749 | HP | Blepharochalasis |
| L7 | 5.52E-05 | 10 | 223 | 7 | 0.031390135 | 0.7 | HP:0001073 | HP | Cigarette-paper scars |
| L7 | 0.000264605 | 458 | 223 | 45 | 0.201793722 | 0.098253275 | HP:0000766 | HP | Abnormal sternum morphology |
| L7 | 0.000565984 | 81 | 223 | 16 | 0.071748879 | 0.197530864 | HP:0000987 | HP | Atypical scarring of skin |
| L7 | 0.002073015 | 34 | 223 | 10 | 0.044843049 | 0.294117647 | HP:0001075 | HP | Atrophic scars |
| L7 | 0.002163962 | 89 | 223 | 16 | 0.071748879 | 0.179775281 | HP:0001191 | HP | Abnormal carpal morphology |
| L7 | 0.002691398 | 70 | 223 | 14 | 0.062780269 | 0.2 | HP:0002616 | HP | Aortic root aneurysm |
| L7 | 0.002691398 | 70 | 223 | 14 | 0.062780269 | 0.2 | HP:0000974 | HP | Hyperextensible skin |
| L7 | 0.003128766 | 28 | 223 | 9 | 0.040358744 | 0.321428571 | HP:0032153 | HP | Joint subluxation |
| L7 | 0.003346861 | 162 | 223 | 22 | 0.098654709 | 0.135802469 | HP:0003019 | HP | Abnormality of the wrist |
| L7 | 0.004006338 | 127 | 223 | 19 | 0.085201794 | 0.149606299 | HP:0001634 | HP | Mitral valve prolapse |
| L7 | 0.005090098 | 911 | 223 | 68 | 0.304932735 | 0.074643249 | HP:0000765 | HP | Abnormal thorax morphology |
| L7 | 0.005995191 | 30 | 223 | 9 | 0.040358744 | 0.3 | HP:0005743 | HP | Avascular necrosis of the capital femoral epiphysis |
| L7 | 0.006299353 | 4 | 223 | 4 | 0.01793722 | 1 | HP:0030009 | HP | Cervical insufficiency |
| L7 | 0.013777119 | 8 | 223 | 5 | 0.022421525 | 0.625 | HP:0025019 | HP | Arterial rupture |
| L7 | 0.014073605 | 304 | 223 | 31 | 0.139013453 | 0.101973684 | HP:0000767 | HP | Pectus excavatum |
| L7 | 0.014573604 | 528 | 223 | 45 | 0.201793722 | 0.085227273 | HP:0004298 | HP | Abnormality of the abdominal wall |
| L7 | 0.01572684 | 13 | 223 | 6 | 0.02690583 | 0.461538462 | HP:0001027 | HP | Soft, doughy skin |
| L7 | 0.015785981 | 103 | 223 | 16 | 0.071748879 | 0.155339806 | HP:0004334 | HP | Dermal atrophy |
| L7 | 0.016418428 | 81 | 223 | 14 | 0.062780269 | 0.172839506 | HP:0025487 | HP | Abnormal bladder morphology |
| L7 | 0.024372593 | 488 | 223 | 42 | 0.188340807 | 0.086065574 | HP:0004299 | HP | Hernia of the abdominal wall |
| L7 | 0.026120732 | 119 | 223 | 17 | 0.076233184 | 0.142857143 | HP:0012727 | HP | Thoracic aortic aneurysm |
| L7 | 0.026873187 | 376 | 223 | 35 | 0.156950673 | 0.093085106 | HP:0001384 | HP | Abnormal hip joint morphology |
| L7 | 0.031234762 | 36 | 223 | 9 | 0.040358744 | 0.25 | HP:0000015 | HP | Bladder diverticulum |
| L7 | 0.031364952 | 493 | 223 | 42 | 0.188340807 | 0.085192698 | HP:0010866 | HP | Abdominal wall defect |
| L7 | 0.032183079 | 242 | 223 | 26 | 0.116591928 | 0.107438017 | HP:0009826 | HP | Limb undergrowth |
| L7 | 0.035467122 | 160 | 223 | 20 | 0.089686099 | 0.125 | HP:0001633 | HP | Abnormal mitral valve morphology |
| L7 | 0.039043169 | 188 | 223 | 22 | 0.098654709 | 0.117021277 | HP:0008067 | HP | Abnormally lax or hyperextensible skin |
| L12 | 0.000949552 | 46 | 234 | 7 | 0.02991453 | 0.152173913 | GO:0005758 | GO:CC | mitochondrial intermembrane space |

| | | | | | | | | | |
|---|---|---|---|---|---|---|---|---|---|
| L12 | 0.002611667 | 7299 | 234 | 138 | 0.58974359 | 0.0189067 | GO:0005737 | GO:CC | cytoplasm |
| L12 | 0.00322425 | 55 | 234 | 7 | 0.02991453 | 0.127272727 | GO:0031970 | GO:CC | organelle envelope lumen |
| L12 | 0.002115184 | 605 | 240 | 24 | 0.1 | 0.039669421 | GO:0015075 | GO:MF | monoatomic ion transmembrane transporter activity |
| L12 | 0.003198214 | 497 | 240 | 21 | 0.0875 | 0.042253521 | GO:0008324 | GO:MF | monoatomic cation transmembrane transporter activity |
| L12 | 0.004054925 | 465 | 240 | 20 | 0.083333333 | 0.043010753 | GO:0022890 | GO:MF | inorganic cation transmembrane transporter activity |
| L12 | 0.005434472 | 556 | 240 | 22 | 0.091666667 | 0.039568345 | GO:0015318 | GO:MF | inorganic molecular entity transmembrane transporter activity |
| L12 | 0.008423112 | 1025 | 240 | 32 | 0.133333333 | 0.031219512 | GO:0005215 | GO:MF | transporter activity |
| L12 | 0.026129788 | 941 | 240 | 29 | 0.120833333 | 0.030818278 | GO:0022857 | GO:MF | transmembrane transporter activity |
| L12 | 0.026886305 | 126 | 240 | 9 | 0.0375 | 0.071428571 | GO:0046943 | GO:MF | carboxylic acid transmembrane transporter activity |
| L12 | 0.028582173 | 127 | 240 | 9 | 0.0375 | 0.070866142 | GO:0005342 | GO:MF | organic acid transmembrane transporter activity |
| L12 | 0.009831688 | 54 | 94 | 8 | 0.085106383 | 0.148148148 | HP:0012705 | HP | Abnormal metabolic brain imaging by MRS |
| L12 | 0.044065658 | 20 | 94 | 5 | 0.053191489 | 0.25 | HP:0025052 | HP | Abnormal brain N-acetyl aspartate level by MRS |
| L14 | 0.000129901 | 288 | 255 | 19 | 0.074509804 | 0.065972222 | GO:0034470 | GO:BP | ncRNA processing |
| L14 | 0.002023462 | 635 | 255 | 27 | 0.105882353 | 0.042519685 | GO:0006974 | GO:BP | DNA damage response |
| L14 | 0.005194676 | 438 | 255 | 21 | 0.082352941 | 0.047945205 | GO:0034660 | GO:BP | ncRNA metabolic process |
| L14 | 0.008547022 | 108 | 255 | 10 | 0.039215686 | 0.092592593 | GO:0006364 | GO:BP | rRNA processing |
| L14 | 0.010599286 | 694 | 255 | 27 | 0.105882353 | 0.038904899 | GO:0006259 | GO:BP | DNA metabolic process |
| L14 | 0.012549992 | 2065 | 255 | 56 | 0.219607843 | 0.027118644 | GO:0043412 | GO:BP | macromolecule modification |
| L14 | 0.041141258 | 388 | 255 | 18 | 0.070588235 | 0.046391753 | GO:0006281 | GO:BP | DNA repair |
| L14 | 0.044857776 | 4050 | 255 | 90 | 0.352941176 | 0.022222222 | GO:0048522 | GO:BP | positive regulation of cellular process |
| L14 | 8.89E-07 | 7299 | 246 | 156 | 0.634146341 | 0.021372791 | GO:0005737 | GO:CC | cytoplasm |
| L14 | 2.46E-06 | 11922 | 246 | 217 | 0.882113821 | 0.018201644 | GO:0005622 | GO:CC | intracellular anatomical structure |
| L14 | 0.000124672 | 2919 | 246 | 76 | 0.308943089 | 0.026036314 | GO:0005654 | GO:CC | nucleoplasm |
| L14 | 0.000266986 | 3979 | 246 | 94 | 0.382113821 | 0.023624026 | GO:0070013 | GO:CC | intracellular organelle lumen |
| L14 | 0.000266986 | 3979 | 246 | 94 | 0.382113821 | 0.023624026 | GO:0043233 | GO:CC | organelle lumen |
| L14 | 0.000266986 | 3979 | 246 | 94 | 0.382113821 | 0.023624026 | GO:0031974 | GO:CC | membrane-enclosed lumen |
| L14 | 0.000680155 | 9673 | 246 | 181 | 0.735772358 | 0.018711878 | GO:0043227 | GO:CC | membrane-bounded organelle |
| L14 | 0.001109454 | 1292 | 246 | 41 | 0.166666667 | 0.031733746 | GO:1902494 | GO:CC | catalytic complex |
| L14 | 0.001173586 | 9306 | 246 | 175 | 0.711382114 | 0.018805072 | GO:0043231 | GO:CC | intracellular membrane-bounded organelle |
| L14 | 0.003158544 | 4 | 246 | 3 | 0.012195122 | 0.75 | GO:0032541 | GO:CC | cortical endoplasmic reticulum |
| L14 | 0.004969163 | 3725 | 246 | 85 | 0.345528455 | 0.022818792 | GO:0031981 | GO:CC | nuclear lumen |
| L14 | 0.005695245 | 39 | 246 | 6 | 0.024390244 | 0.153846154 | GO:0044232 | GO:CC | organelle membrane contact site |
| L14 | 0.00774355 | 1163 | 246 | 36 | 0.146341463 | 0.030954428 | GO:0005739 | GO:CC | mitochondrion |
| L14 | 0.027181206 | 10760 | 246 | 190 | 0.772357724 | 0.017657993 | GO:0043229 | GO:CC | intracellular organelle |
| L14 | 0.0354675 | 11014 | 246 | 193 | 0.784552846 | 0.017523152 | GO:0043226 | GO:CC | organelle |
| L14 | 0.001405322 | 597 | 251 | 25 | 0.099601594 | 0.041876047 | GO:0030674 | GO:MF | protein-macromolecule adaptor activity |
| L14 | 0.004398281 | 679 | 251 | 26 | 0.103585657 | 0.038291605 | GO:0060090 | GO:MF | molecular adaptor activity |
| L14 | 0.008439943 | 2001 | 251 | 53 | 0.211155378 | 0.026486757 | GO:0016740 | GO:MF | transferase activity |
| L14 | 0.015818316 | 5159 | 251 | 107 | 0.426294821 | 0.020740454 | GO:0003824 | GO:MF | catalytic activity |
| L14 | 0.025152552 | 6 | 251 | 3 | 0.011952191 | 0.5 | GO:0042799 | GO:MF | histone H4K20 methyltransferase activity |
| L14 | 0.014113464 | 2089 | 79 | 52 | 0.658227848 | 0.024892293 | HP:0002060 | HP | Abnormal cerebral morphology |

| | | | | | | | | | |
|---|---|---|---|---|---|---|---|---|---|
| L14 | 0.015729868 | 1910 | 79 | 49 | 0.620253165 | 0.02565445 | HP:0002977 | HP | Aplasia/Hypoplasia involving the central nervous system |
| L14 | 0.023291147 | 2120 | 79 | 52 | 0.658227848 | 0.024528302 | HP:0100547 | HP | Abnormal forebrain morphology |
| L15 | 8.75E-19 | 682 | 204 | 46 | 0.225490196 | 0.06744868 | GO:0035239 | GO:BP | tube morphogenesis |
| L15 | 1.11E-17 | 470 | 204 | 38 | 0.18627451 | 0.080851064 | GO:0048514 | GO:BP | blood vessel morphogenesis |
| L15 | 3.77E-17 | 857 | 204 | 49 | 0.240196078 | 0.057176196 | GO:0035295 | GO:BP | tube development |
| L15 | 3.54E-16 | 551 | 204 | 39 | 0.191176471 | 0.070780399 | GO:0001568 | GO:BP | blood vessel development |
| L15 | 2.11E-15 | 580 | 204 | 39 | 0.191176471 | 0.067241379 | GO:0001944 | GO:BP | vasculature development |
| L15 | 5.17E-15 | 1977 | 204 | 71 | 0.348039216 | 0.035912999 | GO:0009653 | GO:BP | anatomical structure morphogenesis |
| L15 | 9.41E-15 | 387 | 204 | 32 | 0.156862745 | 0.082687339 | GO:0001525 | GO:BP | angiogenesis |
| L15 | 3.93E-14 | 853 | 204 | 45 | 0.220588235 | 0.052754982 | GO:0072359 | GO:BP | circulatory system development |
| L15 | 2.34E-13 | 1859 | 204 | 66 | 0.323529412 | 0.035502959 | GO:0007166 | GO:BP | cell surface receptor signaling pathway |
| L15 | 1.97E-12 | 906 | 204 | 44 | 0.215686275 | 0.048565121 | GO:0048646 | GO:BP | anatomical structure formation involved in morphogenesis |
| L15 | 4.68E-12 | 691 | 204 | 38 | 0.18627451 | 0.054992764 | GO:0007167 | GO:BP | enzyme-linked receptor protein signaling pathway |
| L15 | 8.89E-12 | 103 | 204 | 17 | 0.083333333 | 0.165048544 | GO:0003158 | GO:BP | endothelium development |
| L15 | 1.09E-11 | 1850 | 204 | 63 | 0.308823529 | 0.034054054 | GO:0035556 | GO:BP | intracellular signal transduction |
| L15 | 1.16E-11 | 88 | 204 | 16 | 0.078431373 | 0.181818182 | GO:0045446 | GO:BP | endothelial cell differentiation |
| L15 | 1.59E-11 | 1225 | 204 | 50 | 0.245098039 | 0.040816327 | GO:0141124 | GO:BP | intracellular signaling cassette |
| L15 | 3.88E-11 | 942 | 204 | 43 | 0.210784314 | 0.045647558 | GO:0060429 | GO:BP | epithelium development |
| L15 | 8.14E-11 | 2094 | 204 | 66 | 0.323529412 | 0.031518625 | GO:0009966 | GO:BP | regulation of signal transduction |
| L15 | 1.64E-10 | 2758 | 204 | 77 | 0.37745098 | 0.027918782 | GO:0048583 | GO:BP | regulation of response to stimulus |
| L15 | 3.13E-10 | 454 | 204 | 29 | 0.142156863 | 0.063876652 | GO:0071363 | GO:BP | cellular response to growth factor stimulus |
| L15 | 3.21E-10 | 1520 | 204 | 54 | 0.264705882 | 0.035526316 | GO:0009888 | GO:BP | tissue development |
| L15 | 6.73E-10 | 468 | 204 | 29 | 0.142156863 | 0.061965812 | GO:0070848 | GO:BP | response to growth factor |
| L15 | 3.25E-09 | 245 | 204 | 21 | 0.102941176 | 0.085714286 | GO:0090130 | GO:BP | tissue migration |
| L15 | 3.30E-09 | 105 | 204 | 15 | 0.073529412 | 0.142857143 | GO:0002040 | GO:BP | sprouting angiogenesis |
| L15 | 4.98E-09 | 581 | 204 | 31 | 0.151960784 | 0.053356282 | GO:0022603 | GO:BP | regulation of anatomical structure morphogenesis |
| L15 | 7.55E-09 | 1009 | 204 | 41 | 0.200980392 | 0.040634291 | GO:2000026 | GO:BP | regulation of multicellular organismal development |
| L15 | 1.36E-08 | 3317 | 204 | 82 | 0.401960784 | 0.024721134 | GO:0007275 | GO:BP | multicellular organism development |
| L15 | 1.36E-08 | 2754 | 204 | 73 | 0.357843137 | 0.026506899 | GO:0048731 | GO:BP | system development |
| L15 | 1.51E-08 | 2401 | 204 | 67 | 0.328431373 | 0.02790504 | GO:0023051 | GO:BP | regulation of signaling |
| L15 | 1.57E-08 | 2403 | 204 | 67 | 0.328431373 | 0.027881814 | GO:0010646 | GO:BP | regulation of cell communication |
| L15 | 1.89E-08 | 9739 | 204 | 163 | 0.799019608 | 0.016736831 | GO:0065007 | GO:BP | biological regulation |
| L15 | 2.02E-08 | 241 | 204 | 20 | 0.098039216 | 0.082987552 | GO:0010631 | GO:BP | epithelial cell migration |
| L15 | 2.35E-08 | 243 | 204 | 20 | 0.098039216 | 0.082304527 | GO:0090132 | GO:BP | epithelium migration |
| L15 | 2.50E-08 | 4931 | 204 | 105 | 0.514705882 | 0.021293855 | GO:0007165 | GO:BP | signal transduction |
| L15 | 3.18E-08 | 510 | 204 | 28 | 0.137254902 | 0.054901961 | GO:0030855 | GO:BP | epithelial cell differentiation |
| L15 | 3.60E-08 | 750 | 204 | 34 | 0.166666667 | 0.045333333 | GO:0040012 | GO:BP | regulation of locomotion |
| L15 | 3.91E-08 | 9049 | 204 | 155 | 0.759803922 | 0.017128965 | GO:0050794 | GO:BP | regulation of cellular process |
| L15 | 5.30E-08 | 1785 | 204 | 55 | 0.269607843 | 0.030812325 | GO:0050793 | GO:BP | regulation of developmental process |
| L15 | 5.80E-08 | 285 | 204 | 21 | 0.102941176 | 0.073684211 | GO:0007178 | GO:BP | transmembrane receptor protein serine/threonine kinase signaling pathway |
| L15 | 6.07E-08 | 256 | 204 | 20 | 0.098039216 | 0.078125 | GO:0141091 | GO:BP | transforming growth factor beta receptor superfamily signaling pathway |

| | | | | | | | | | |
|---|---|---|---|---|---|---|---|---|---|
| L15 | 8.57E-08 | 291 | 204 | 21 | 0.102941176 | 0.072164948 | GO:0060562 | GO:BP | epithelial tube morphogenesis |
| L15 | 9.16E-08 | 908 | 204 | 37 | 0.181372549 | 0.040748899 | GO:0071495 | GO:BP | cellular response to endogenous stimulus |
| L15 | 1.13E-07 | 870 | 204 | 36 | 0.176470588 | 0.04137931 | GO:0040011 | GO:BP | locomotion |
| L15 | 1.34E-07 | 4205 | 204 | 93 | 0.455882353 | 0.022116528 | GO:0048856 | GO:BP | anatomical structure development |
| L15 | 1.62E-07 | 333 | 204 | 22 | 0.107843137 | 0.066066066 | GO:0001667 | GO:BP | ameboidal-type cell migration |
| L15 | 1.78E-07 | 669 | 204 | 31 | 0.151960784 | 0.046337818 | GO:0030334 | GO:BP | regulation of cell migration |
| L15 | 2.09E-07 | 715 | 204 | 32 | 0.156862745 | 0.044755245 | GO:2000145 | GO:BP | regulation of cell motility |
| L15 | 2.22E-07 | 675 | 204 | 31 | 0.151960784 | 0.045925926 | GO:0051093 | GO:BP | negative regulation of developmental process |
| L15 | 2.38E-07 | 1176 | 204 | 42 | 0.205882353 | 0.035714286 | GO:1902531 | GO:BP | regulation of intracellular signal transduction |
| L15 | 2.40E-07 | 2140 | 204 | 60 | 0.294117647 | 0.028037383 | GO:0051239 | GO:BP | regulation of multicellular organismal process |
| L15 | 2.44E-07 | 444 | 204 | 25 | 0.12254902 | 0.056306306 | GO:0007507 | GO:BP | heart development |
| L15 | 2.52E-07 | 9499 | 204 | 158 | 0.774509804 | 0.01663333 | GO:0050789 | GO:BP | regulation of biological process |
| L15 | 2.74E-07 | 990 | 204 | 38 | 0.18627451 | 0.038383838 | GO:0009719 | GO:BP | response to endogenous stimulus |
| L15 | 3.04E-07 | 15 | 204 | 7 | 0.034313725 | 0.466666667 | GO:0060841 | GO:BP | venous blood vessel development |
| L15 | 3.49E-07 | 5965 | 204 | 116 | 0.568627451 | 0.019446773 | GO:0051716 | GO:BP | cellular response to stimulus |
| L15 | 3.71E-07 | 196 | 204 | 17 | 0.083333333 | 0.086734694 | GO:0090092 | GO:BP | regulation of transmembrane receptor protein serine/threonine kinase signaling pathway |
| L15 | 3.76E-07 | 4492 | 204 | 96 | 0.470588235 | 0.021371327 | GO:0032502 | GO:BP | developmental process |
| L15 | 3.79E-07 | 224 | 204 | 18 | 0.088235294 | 0.080357143 | GO:0060485 | GO:BP | mesenchyme development |
| L15 | 3.85E-07 | 3938 | 204 | 88 | 0.431372549 | 0.022346369 | GO:0048519 | GO:BP | negative regulation of biological process |
| L15 | 3.91E-07 | 820 | 204 | 34 | 0.166666667 | 0.041463415 | GO:0009887 | GO:BP | animal organ morphogenesis |
| L15 | 4.07E-07 | 146 | 204 | 15 | 0.073529412 | 0.102739726 | GO:0048754 | GO:BP | branching morphogenesis of an epithelial tube |
| L15 | 6.68E-07 | 51 | 204 | 10 | 0.049019608 | 0.196078431 | GO:0002042 | GO:BP | cell migration involved in sprouting angiogenesis |
| L15 | 9.74E-07 | 88 | 204 | 12 | 0.058823529 | 0.136363636 | GO:0060840 | GO:BP | artery development |
| L15 | 1.21E-06 | 54 | 204 | 10 | 0.049019608 | 0.185185185 | GO:0035924 | GO:BP | cellular response to vascular endothelial growth factor stimulus |
| L15 | 1.30E-06 | 4447 | 204 | 94 | 0.460784314 | 0.021137846 | GO:0048518 | GO:BP | positive regulation of biological process |
| L15 | 1.58E-06 | 1151 | 204 | 40 | 0.196078431 | 0.034752389 | GO:0042127 | GO:BP | regulation of cell population proliferation |
| L15 | 1.73E-06 | 691 | 204 | 30 | 0.147058824 | 0.04341534 | GO:1902533 | GO:BP | positive regulation of intracellular signal transduction |
| L15 | 1.86E-06 | 1108 | 204 | 39 | 0.191176471 | 0.035198556 | GO:0045595 | GO:BP | regulation of cell differentiation |
| L15 | 1.89E-06 | 5276 | 204 | 105 | 0.514705882 | 0.01990144 | GO:0023052 | GO:BP | signaling |
| L15 | 2.59E-06 | 1072 | 204 | 38 | 0.18627451 | 0.035447761 | GO:0016477 | GO:BP | cell migration |
| L15 | 3.50E-06 | 6965 | 204 | 126 | 0.617647059 | 0.018090452 | GO:0050896 | GO:BP | response to stimulus |
| L15 | 3.63E-06 | 3696 | 204 | 82 | 0.401960784 | 0.022186147 | GO:0048523 | GO:BP | negative regulation of cellular process |
| L15 | 3.72E-06 | 145 | 204 | 14 | 0.068627451 | 0.096551724 | GO:0007179 | GO:BP | transforming growth factor beta receptor signaling pathway |
| L15 | 4.10E-06 | 229 | 204 | 17 | 0.083333333 | 0.074235808 | GO:0090287 | GO:BP | regulation of cellular response to growth factor stimulus |
| L15 | 4.76E-06 | 174 | 204 | 15 | 0.073529412 | 0.086206897 | GO:0043542 | GO:BP | endothelial cell migration |
| L15 | 5.01E-06 | 62 | 204 | 10 | 0.049019608 | 0.161290323 | GO:0048844 | GO:BP | artery morphogenesis |
| L15 | 5.33E-06 | 5366 | 204 | 105 | 0.514705882 | 0.019567648 | GO:0007154 | GO:BP | cell communication |
| L15 | 5.43E-06 | 516 | 204 | 25 | 0.12254902 | 0.048449612 | GO:0048729 | GO:BP | tissue morphogenesis |
| L15 | 5.52E-06 | 102 | 204 | 12 | 0.058823529 | 0.117647059 | GO:0043534 | GO:BP | blood vessel endothelial cell migration |
| L15 | 6.03E-06 | 177 | 204 | 15 | 0.073529412 | 0.084745763 | GO:0010632 | GO:BP | regulation of epithelial cell migration |
| L15 | 6.03E-06 | 177 | 204 | 15 | 0.073529412 | 0.084745763 | GO:0061138 | GO:BP | morphogenesis of a branching epithelium |

| | | | | | | | | | |
|---|---|---|---|---|---|---|---|---|---|
| L15 | 6.48E-06 | 1417 | 204 | 44 | 0.215686275 | 0.031051517 | GO:0008283 | GO:BP | cell population proliferation |
| L15 | 7.80E-06 | 2897 | 204 | 69 | 0.338235294 | 0.023817742 | GO:0030154 | GO:BP | cell differentiation |
| L15 | 7.80E-06 | 2897 | 204 | 69 | 0.338235294 | 0.023817742 | GO:0048869 | GO:BP | cellular developmental process |
| L15 | 9.67E-06 | 929 | 204 | 34 | 0.166666667 | 0.036598493 | GO:0007155 | GO:BP | cell adhesion |
| L15 | 1.05E-05 | 932 | 204 | 34 | 0.166666667 | 0.036480687 | GO:0009790 | GO:BP | embryo development |
| L15 | 1.37E-05 | 51 | 204 | 9 | 0.044117647 | 0.176470588 | GO:0003170 | GO:BP | heart valve development |
| L15 | 1.59E-05 | 1095 | 204 | 37 | 0.181372549 | 0.033789954 | GO:0009967 | GO:BP | positive regulation of signal transduction |
| L15 | 1.59E-05 | 190 | 204 | 15 | 0.073529412 | 0.078947368 | GO:0001763 | GO:BP | morphogenesis of a branching structure |
| L15 | 1.71E-05 | 37 | 204 | 8 | 0.039215686 | 0.216216216 | GO:0090049 | GO:BP | regulation of cell migration involved in sprouting angiogenesis |
| L15 | 1.98E-05 | 431 | 204 | 22 | 0.107843137 | 0.051044084 | GO:0002009 | GO:BP | morphogenesis of an epithelium |
| L15 | 2.40E-05 | 4050 | 204 | 85 | 0.416666667 | 0.020987654 | GO:0048522 | GO:BP | positive regulation of cellular process |
| L15 | 2.78E-05 | 26 | 204 | 7 | 0.034313725 | 0.269230769 | GO:0001945 | GO:BP | lymph vessel development |
| L15 | 3.53E-05 | 1232 | 204 | 39 | 0.191176471 | 0.031655844 | GO:0048870 | GO:BP | cell motility |
| L15 | 3.65E-05 | 173 | 204 | 14 | 0.068627451 | 0.080924855 | GO:0071560 | GO:BP | cellular response to transforming growth factor beta stimulus |
| L15 | 3.71E-05 | 932 | 204 | 33 | 0.161764706 | 0.035407725 | GO:0051094 | GO:BP | positive regulation of developmental process |
| L15 | 4.07E-05 | 41 | 204 | 8 | 0.039215686 | 0.195121951 | GO:0003197 | GO:BP | endocardial cushion development |
| L15 | 4.87E-05 | 177 | 204 | 14 | 0.068627451 | 0.079096045 | GO:0071559 | GO:BP | response to transforming growth factor beta |
| L15 | 4.90E-05 | 28 | 204 | 7 | 0.034313725 | 0.25 | GO:0003176 | GO:BP | aortic valve development |
| L15 | 4.96E-05 | 78 | 204 | 10 | 0.049019608 | 0.128205128 | GO:0043535 | GO:BP | regulation of blood vessel endothelial cell migration |
| L15 | 4.97E-05 | 42 | 204 | 8 | 0.039215686 | 0.19047619 | GO:0003179 | GO:BP | heart valve morphogenesis |
| L15 | 5.03E-05 | 207 | 204 | 15 | 0.073529412 | 0.072463768 | GO:0003007 | GO:BP | heart morphogenesis |
| L15 | 5.65E-05 | 125 | 204 | 12 | 0.058823529 | 0.096 | GO:0030509 | GO:BP | BMP signaling pathway |
| L15 | 6.22E-05 | 382 | 204 | 20 | 0.098039216 | 0.052356021 | GO:0030335 | GO:BP | positive regulation of cell migration |
| L15 | 6.46E-05 | 310 | 204 | 18 | 0.088235294 | 0.058064516 | GO:0007264 | GO:BP | small GTPase-mediated signal transduction |
| L15 | 6.47E-05 | 181 | 204 | 14 | 0.068627451 | 0.077348066 | GO:0048762 | GO:BP | mesenchymal cell differentiation |
| L15 | 7.38E-05 | 128 | 204 | 12 | 0.058823529 | 0.09375 | GO:0001837 | GO:BP | epithelial to mesenchymal transition |
| L15 | 7.83E-05 | 214 | 204 | 15 | 0.073529412 | 0.070093458 | GO:0045765 | GO:BP | regulation of angiogenesis |
| L15 | 8.21E-05 | 2183 | 204 | 55 | 0.269607843 | 0.025194686 | GO:0048513 | GO:BP | animal organ development |
| L15 | 9.05E-05 | 595 | 204 | 25 | 0.12254902 | 0.042016807 | GO:0045597 | GO:BP | positive regulation of cell differentiation |
| L15 | 9.75E-05 | 5110 | 204 | 98 | 0.480392157 | 0.019178082 | GO:0032501 | GO:BP | multicellular organismal process |
| L15 | 0.000106536 | 557 | 204 | 24 | 0.117647059 | 0.043087971 | GO:0030155 | GO:BP | regulation of cell adhesion |
| L15 | 0.000112867 | 220 | 204 | 15 | 0.073529412 | 0.068181818 | GO:1901342 | GO:BP | regulation of vasculature development |
| L15 | 0.000113135 | 133 | 204 | 12 | 0.058823529 | 0.090225564 | GO:0071773 | GO:BP | cellular response to BMP stimulus |
| L15 | 0.000113135 | 133 | 204 | 12 | 0.058823529 | 0.090225564 | GO:0071772 | GO:BP | response to BMP |
| L15 | 0.000118799 | 1236 | 204 | 38 | 0.18627451 | 0.030744337 | GO:0023056 | GO:BP | positive regulation of signaling |
| L15 | 0.00012296 | 134 | 204 | 12 | 0.058823529 | 0.089552239 | GO:0010594 | GO:BP | regulation of endothelial cell migration |
| L15 | 0.000124008 | 65 | 204 | 9 | 0.044117647 | 0.138461538 | GO:0001570 | GO:BP | vasculogenesis |
| L15 | 0.000143429 | 402 | 204 | 20 | 0.098039216 | 0.049751244 | GO:2000147 | GO:BP | positive regulation of cell motility |
| L15 | 0.000168045 | 33 | 204 | 7 | 0.034313725 | 0.212121212 | GO:1905314 | GO:BP | semi-lunar valve development |
| L15 | 0.000176745 | 49 | 204 | 8 | 0.039215686 | 0.163265306 | GO:0035904 | GO:BP | aorta development |
| L15 | 0.000177408 | 800 | 204 | 29 | 0.142156863 | 0.03625 | GO:0051241 | GO:BP | negative regulation of multicellular organismal process |

| | | | | | | | | | |
|---|---|---|---|---|---|---|---|---|---|
| L15 | 0.000204172 | 21 | 204 | 6 | 0.029411765 | 0.285714286 | GO:0045920 | GO:BP | negative regulation of exocytosis |
| L15 | 0.000208246 | 50 | 204 | 8 | 0.039215686 | 0.16 | GO:0060390 | GO:BP | regulation of SMAD protein signal transduction |
| L15 | 0.000212427 | 169 | 204 | 13 | 0.06372549 | 0.076923077 | GO:0002064 | GO:BP | epithelial cell development |
| L15 | 0.000221993 | 413 | 204 | 20 | 0.098039216 | 0.04842615 | GO:0040017 | GO:BP | positive regulation of locomotion |
| L15 | 0.000263524 | 2509 | 204 | 59 | 0.289215686 | 0.023515345 | GO:0009893 | GO:BP | positive regulation of metabolic process |
| L15 | 0.000282697 | 3635 | 204 | 76 | 0.37254902 | 0.02090784 | GO:0080090 | GO:BP | regulation of primary metabolic process |
| L15 | 0.000315281 | 2272 | 204 | 55 | 0.269607843 | 0.024207746 | GO:0031325 | GO:BP | positive regulation of cellular metabolic process |
| L15 | 0.000318436 | 36 | 204 | 7 | 0.034313725 | 0.194444444 | GO:0001974 | GO:BP | blood vessel remodeling |
| L15 | 0.000320619 | 1231 | 204 | 37 | 0.181372549 | 0.030056864 | GO:0010647 | GO:BP | positive regulation of cell communication |
| L15 | 0.00036804 | 96 | 204 | 10 | 0.049019608 | 0.104166667 | GO:0017015 | GO:BP | regulation of transforming growth factor beta receptor signaling pathway |
| L15 | 0.000387101 | 3525 | 204 | 74 | 0.362745098 | 0.020992908 | GO:0051171 | GO:BP | regulation of nitrogen compound metabolic process |
| L15 | 0.000403282 | 122 | 204 | 11 | 0.053921569 | 0.090163934 | GO:0001935 | GO:BP | endothelial cell proliferation |
| L15 | 0.00040607 | 97 | 204 | 10 | 0.049019608 | 0.103092784 | GO:1903844 | GO:BP | regulation of cellular response to transforming growth factor beta stimulus |
| L15 | 0.00041516 | 2291 | 204 | 55 | 0.269607843 | 0.024006984 | GO:0010604 | GO:BP | positive regulation of macromolecule metabolic process |
| L15 | 0.000441662 | 13 | 204 | 5 | 0.024509804 | 0.384615385 | GO:1903306 | GO:BP | negative regulation of regulated secretory pathway |
| L15 | 0.000447079 | 1194 | 204 | 36 | 0.176470588 | 0.030150754 | GO:0048585 | GO:BP | negative regulation of response to stimulus |
| L15 | 0.000492404 | 1417 | 204 | 40 | 0.196078431 | 0.028228652 | GO:0006357 | GO:BP | regulation of transcription by RNA polymerase II |
| L15 | 0.000500215 | 1938 | 204 | 49 | 0.240196078 | 0.025283798 | GO:0051173 | GO:BP | positive regulation of nitrogen compound metabolic process |
| L15 | 0.000516362 | 125 | 204 | 11 | 0.053921569 | 0.088 | GO:0072073 | GO:BP | kidney epithelium development |
| L15 | 0.000569 | 39 | 204 | 7 | 0.034313725 | 0.179487179 | GO:0061028 | GO:BP | establishment of endothelial barrier |
| L15 | 0.000634768 | 1158 | 204 | 35 | 0.171568627 | 0.030224525 | GO:0051240 | GO:BP | positive regulation of multicellular organismal process |
| L15 | 0.000652694 | 102 | 204 | 10 | 0.049019608 | 0.098039216 | GO:0001936 | GO:BP | regulation of endothelial cell proliferation |
| L15 | 0.000769499 | 80 | 204 | 9 | 0.044117647 | 0.1125 | GO:0060395 | GO:BP | SMAD protein signal transduction |
| L15 | 0.000922959 | 494 | 204 | 21 | 0.102941176 | 0.042510121 | GO:0045596 | GO:BP | negative regulation of cell differentiation |
| L15 | 0.000923166 | 332 | 204 | 17 | 0.083333333 | 0.051204819 | GO:0045785 | GO:BP | positive regulation of cell adhesion |
| L15 | 0.001010437 | 15 | 204 | 5 | 0.024509804 | 0.333333333 | GO:1903587 | GO:BP | regulation of blood vessel endothelial cell proliferation involved in sprouting angiogenesis |
| L15 | 0.001010437 | 15 | 204 | 5 | 0.024509804 | 0.333333333 | GO:1901201 | GO:BP | regulation of extracellular matrix assembly |
| L15 | 0.001048419 | 27 | 204 | 6 | 0.029411765 | 0.222222222 | GO:0035909 | GO:BP | aorta morphogenesis |
| L15 | 0.001051127 | 1516 | 204 | 41 | 0.200980392 | 0.027044855 | GO:0006366 | GO:BP | transcription by RNA polymerase II |
| L15 | 0.001066209 | 7 | 204 | 4 | 0.019607843 | 0.571428571 | GO:0060836 | GO:BP | lymphatic endothelial cell differentiation |
| L15 | 0.001121911 | 135 | 204 | 11 | 0.053921569 | 0.081481481 | GO:0030856 | GO:BP | regulation of epithelial cell differentiation |
| L15 | 0.001346978 | 44 | 204 | 7 | 0.034313725 | 0.159090909 | GO:0072132 | GO:BP | mesenchyme morphogenesis |
| L15 | 0.001582399 | 87 | 204 | 9 | 0.044117647 | 0.103448276 | GO:0090100 | GO:BP | positive regulation of transmembrane receptor protein serine/threonine kinase signaling pathway |
| L15 | 0.001659656 | 427 | 204 | 19 | 0.093137255 | 0.044496487 | GO:0007169 | GO:BP | transmembrane receptor protein tyrosine kinase signaling pathway |
| L15 | 0.001709219 | 470 | 204 | 20 | 0.098039216 | 0.042553191 | GO:0000165 | GO:BP | MAPK cascade |
| L15 | 0.001972671 | 4596 | 204 | 87 | 0.426470588 | 0.018929504 | GO:0019222 | GO:BP | regulation of metabolic process |
| L15 | 0.002112229 | 8 | 204 | 4 | 0.019607843 | 0.5 | GO:0048845 | GO:BP | venous blood vessel morphogenesis |
| L15 | 0.002336167 | 117 | 204 | 10 | 0.049019608 | 0.085470085 | GO:0010634 | GO:BP | positive regulation of epithelial cell migration |
| L15 | 0.002632876 | 3 | 204 | 3 | 0.014705882 | 1 | GO:1903847 | GO:BP | regulation of aorta morphogenesis |
| L15 | 0.002632876 | 3 | 204 | 3 | 0.014705882 | 1 | GO:1903849 | GO:BP | positive regulation of aorta morphogenesis |
| L15 | 0.002632876 | 3 | 204 | 3 | 0.014705882 | 1 | GO:0003250 | GO:BP | regulation of cell proliferation involved in heart valve morphogenesis |

| | | | | | | | | | |
|---|---|---|---|---|---|---|---|---|---|
| L15 | 0.002632876 | 3 | 204 | 3 | 0.014705882 | 1 | GO:0003249 | GO:BP | cell proliferation involved in heart valve morphogenesis |
| L15 | 0.002632876 | 3 | 204 | 3 | 0.014705882 | 1 | GO:2000793 | GO:BP | cell proliferation involved in heart valve development |
| L15 | 0.00270581 | 359 | 204 | 17 | 0.083333333 | 0.04735376 | GO:0050673 | GO:BP | epithelial cell proliferation |
| L15 | 0.00279901 | 18 | 204 | 5 | 0.024509804 | 0.277777778 | GO:0001946 | GO:BP | lymphangiogenesis |
| L15 | 0.002869046 | 49 | 204 | 7 | 0.034313725 | 0.142857143 | GO:0001885 | GO:BP | endothelial cell development |
| L15 | 0.003052037 | 32 | 204 | 6 | 0.029411765 | 0.1875 | GO:0003203 | GO:BP | endocardial cushion morphogenesis |
| L15 | 0.003340337 | 491 | 204 | 20 | 0.098039216 | 0.040733198 | GO:0048589 | GO:BP | developmental growth |
| L15 | 0.003393655 | 406 | 204 | 18 | 0.088235294 | 0.044334975 | GO:0043408 | GO:BP | regulation of MAPK cascade |
| L15 | 0.003692924 | 33 | 204 | 6 | 0.029411765 | 0.181818182 | GO:0060391 | GO:BP | positive regulation of SMAD protein signal transduction |
| L15 | 0.003692924 | 33 | 204 | 6 | 0.029411765 | 0.181818182 | GO:0045601 | GO:BP | regulation of endothelial cell differentiation |
| L15 | 0.003736812 | 1593 | 204 | 41 | 0.200980392 | 0.025737602 | GO:0048584 | GO:BP | positive regulation of response to stimulus |
| L15 | 0.003761455 | 19 | 204 | 5 | 0.024509804 | 0.263157895 | GO:0003181 | GO:BP | atrioventricular valve morphogenesis |
| L15 | 0.003766028 | 9 | 204 | 4 | 0.019607843 | 0.444444444 | GO:2001214 | GO:BP | positive regulation of vasculogenesis |
| L15 | 0.004080644 | 588 | 204 | 22 | 0.107843137 | 0.037414966 | GO:0098609 | GO:BP | cell-cell adhesion |
| L15 | 0.004940372 | 595 | 204 | 22 | 0.107843137 | 0.03697479 | GO:0043009 | GO:BP | chordate embryonic development |
| L15 | 0.004966168 | 20 | 204 | 5 | 0.024509804 | 0.25 | GO:0002043 | GO:BP | blood vessel endothelial cell proliferation involved in sprouting angiogenesis |
| L15 | 0.005126468 | 100 | 204 | 9 | 0.044117647 | 0.09 | GO:0090101 | GO:BP | negative regulation of transmembrane receptor protein serine/threonine kinase signaling pathway |
| L15 | 0.005241364 | 260 | 204 | 14 | 0.068627451 | 0.053846154 | GO:0072001 | GO:BP | renal system development |
| L15 | 0.005292398 | 4262 | 204 | 81 | 0.397058824 | 0.019005162 | GO:0031323 | GO:BP | regulation of cellular metabolic process |
| L15 | 0.005304255 | 35 | 204 | 6 | 0.029411765 | 0.171428571 | GO:0001569 | GO:BP | branching involved in blood vessel morphogenesis |
| L15 | 0.005473504 | 646 | 204 | 23 | 0.112745098 | 0.035603715 | GO:0040007 | GO:BP | growth |
| L15 | 0.005615687 | 647 | 204 | 23 | 0.112745098 | 0.035548686 | GO:0008284 | GO:BP | positive regulation of cell population proliferation |
| L15 | 0.005620672 | 54 | 204 | 7 | 0.034313725 | 0.12962963 | GO:0030512 | GO:BP | negative regulation of transforming growth factor beta receptor signaling pathway |
| L15 | 0.005743416 | 949 | 204 | 29 | 0.142156863 | 0.030558483 | GO:0009968 | GO:BP | negative regulation of signal transduction |
| L15 | 0.005844488 | 192 | 204 | 12 | 0.058823529 | 0.0625 | GO:0070372 | GO:BP | regulation of ERK1 and ERK2 cascade |
| L15 | 0.006287773 | 604 | 204 | 22 | 0.107843137 | 0.036423841 | GO:0009792 | GO:BP | embryo development ending in birth or egg hatching |
| L15 | 0.006373365 | 1007 | 204 | 30 | 0.147058824 | 0.02979146 | GO:0010648 | GO:BP | negative regulation of cell communication |
| L15 | 0.006454292 | 21 | 204 | 5 | 0.024509804 | 0.238095238 | GO:0036303 | GO:BP | lymph vessel morphogenesis |
| L15 | 0.006454292 | 21 | 204 | 5 | 0.024509804 | 0.238095238 | GO:0090050 | GO:BP | positive regulation of cell migration involved in sprouting angiogenesis |
| L15 | 0.006761723 | 1010 | 204 | 30 | 0.147058824 | 0.02970297 | GO:0023057 | GO:BP | negative regulation of signaling |
| L15 | 0.007445808 | 37 | 204 | 6 | 0.029411765 | 0.162162162 | GO:0038084 | GO:BP | vascular endothelial growth factor signaling pathway |
| L15 | 0.00768844 | 105 | 204 | 9 | 0.044117647 | 0.085714286 | GO:0003231 | GO:BP | cardiac ventricle development |
| L15 | 0.007713314 | 911 | 204 | 28 | 0.137254902 | 0.030735456 | GO:0045944 | GO:BP | positive regulation of transcription by RNA polymerase II |
| L15 | 0.00776834 | 269 | 204 | 14 | 0.068627451 | 0.05204461 | GO:0043010 | GO:BP | camera-type eye development |
| L15 | 0.008232202 | 1825 | 204 | 44 | 0.215686275 | 0.024109589 | GO:0010557 | GO:BP | positive regulation of macromolecule biosynthetic process |
| L15 | 0.008248441 | 80 | 204 | 8 | 0.039215686 | 0.1 | GO:0010717 | GO:BP | regulation of epithelial to mesenchymal transition |
| L15 | 0.008314952 | 106 | 204 | 9 | 0.044117647 | 0.08490566 | GO:0003206 | GO:BP | cardiac chamber morphogenesis |
| L15 | 0.008753458 | 38 | 204 | 6 | 0.029411765 | 0.157894737 | GO:0085029 | GO:BP | extracellular matrix assembly |
| L15 | 0.009160856 | 58 | 204 | 7 | 0.034313725 | 0.120689655 | GO:0003208 | GO:BP | cardiac ventricle morphogenesis |
| L15 | 0.009492473 | 717 | 204 | 24 | 0.117647059 | 0.033472803 | GO:0044087 | GO:BP | regulation of cellular component biogenesis |
| L15 | 0.00967769 | 11 | 204 | 4 | 0.019607843 | 0.363636364 | GO:0003183 | GO:BP | mitral valve morphogenesis |

| | | | | | | | | | |
|---|---|---|---|---|---|---|---|---|---|
| L15 | 0.00967769 | 11 | 204 | 4 | 0.019607843 | 0.363636364 | GO:0003157 | GO:BP | endocardium development |
| L15 | 0.010437468 | 4 | 204 | 3 | 0.014705882 | 0.75 | GO:0003273 | GO:BP | cell migration involved in endocardial cushion formation |
| L15 | 0.010464936 | 23 | 204 | 5 | 0.024509804 | 0.217391304 | GO:0003171 | GO:BP | atrioventricular valve development |
| L15 | 0.010464936 | 23 | 204 | 5 | 0.024509804 | 0.217391304 | GO:0003180 | GO:BP | aortic valve morphogenesis |
| L15 | 0.011928399 | 40 | 204 | 6 | 0.029411765 | 0.15 | GO:0055010 | GO:BP | ventricular cardiac muscle tissue morphogenesis |
| L15 | 0.011928399 | 40 | 204 | 6 | 0.029411765 | 0.15 | GO:1903053 | GO:BP | regulation of extracellular matrix organization |
| L15 | 0.012584157 | 141 | 204 | 10 | 0.049019608 | 0.070921986 | GO:0003205 | GO:BP | cardiac chamber development |
| L15 | 0.013089566 | 24 | 204 | 5 | 0.024509804 | 0.208333333 | GO:0003272 | GO:BP | endocardial cushion formation |
| L15 | 0.013370325 | 174 | 204 | 11 | 0.053921569 | 0.063218391 | GO:0030324 | GO:BP | lung development |
| L15 | 0.01419282 | 86 | 204 | 8 | 0.039215686 | 0.093023256 | GO:0010595 | GO:BP | positive regulation of endothelial cell migration |
| L15 | 0.014379293 | 12 | 204 | 4 | 0.019607843 | 0.333333333 | GO:0003174 | GO:BP | mitral valve development |
| L15 | 0.014379293 | 12 | 204 | 4 | 0.019607843 | 0.333333333 | GO:2001212 | GO:BP | regulation of vasculogenesis |
| L15 | 0.014615599 | 210 | 204 | 12 | 0.058823529 | 0.057142857 | GO:0070371 | GO:BP | ERK1 and ERK2 cascade |
| L15 | 0.014796888 | 1274 | 204 | 34 | 0.166666667 | 0.026687598 | GO:0051254 | GO:BP | positive regulation of RNA metabolic process |
| L15 | 0.015648634 | 1810 | 204 | 43 | 0.210784314 | 0.023756906 | GO:0065008 | GO:BP | regulation of biological quality |
| L15 | 0.015724145 | 177 | 204 | 11 | 0.053921569 | 0.062146893 | GO:0030323 | GO:BP | respiratory tube development |
| L15 | 0.016440327 | 249 | 204 | 13 | 0.06372549 | 0.052208835 | GO:0001822 | GO:BP | kidney development |
| L15 | 0.017585038 | 289 | 204 | 14 | 0.068627451 | 0.048442907 | GO:0050678 | GO:BP | regulation of epithelial cell proliferation |
| L15 | 0.017798636 | 1881 | 204 | 44 | 0.215686275 | 0.023391813 | GO:0031328 | GO:BP | positive regulation of cellular biosynthetic process |
| L15 | 0.017803598 | 64 | 204 | 7 | 0.034313725 | 0.109375 | GO:0003151 | GO:BP | outflow tract morphogenesis |
| L15 | 0.018185916 | 1699 | 204 | 41 | 0.200980392 | 0.024131842 | GO:0051128 | GO:BP | regulation of cellular component organization |
| L15 | 0.0183833 | 43 | 204 | 6 | 0.029411765 | 0.139534884 | GO:0030857 | GO:BP | negative regulation of epithelial cell differentiation |
| L15 | 0.020071019 | 554 | 204 | 20 | 0.098039216 | 0.036101083 | GO:0060284 | GO:BP | regulation of cell development |
| L15 | 0.020573796 | 13 | 204 | 4 | 0.019607843 | 0.307692308 | GO:0003198 | GO:BP | epithelial to mesenchymal transition involved in endocardial cushion formation |
| L15 | 0.020619197 | 1410 | 204 | 36 | 0.176470588 | 0.025531915 | GO:0045935 | GO:BP | positive regulation of nucleobase-containing compound metabolic process |
| L15 | 0.020719616 | 463 | 204 | 18 | 0.088235294 | 0.03887689 | GO:0061061 | GO:BP | muscle structure development |
| L15 | 0.02132591 | 294 | 204 | 14 | 0.068627451 | 0.047619048 | GO:0043410 | GO:BP | positive regulation of MAPK cascade |
| L15 | 0.021758993 | 335 | 204 | 15 | 0.073529412 | 0.044776119 | GO:0022407 | GO:BP | regulation of cell-cell adhesion |
| L15 | 0.022014652 | 1897 | 204 | 44 | 0.215686275 | 0.023194518 | GO:0009891 | GO:BP | positive regulation of biosynthetic process |
| L15 | 0.023025268 | 257 | 204 | 13 | 0.06372549 | 0.050583658 | GO:0031589 | GO:BP | cell-substrate adhesion |
| L15 | 0.024072377 | 45 | 204 | 6 | 0.029411765 | 0.133333333 | GO:0043536 | GO:BP | positive regulation of blood vessel endothelial cell migration |
| L15 | 0.02411534 | 4207 | 204 | 78 | 0.382352941 | 0.018540528 | GO:0060255 | GO:BP | regulation of macromolecule metabolic process |
| L15 | 0.024139583 | 27 | 204 | 5 | 0.024509804 | 0.185185185 | GO:0060317 | GO:BP | cardiac epithelial to mesenchymal transition |
| L15 | 0.024172912 | 67 | 204 | 7 | 0.034313725 | 0.104477612 | GO:0001938 | GO:BP | positive regulation of endothelial cell proliferation |
| L15 | 0.025860806 | 5 | 204 | 3 | 0.014705882 | 0.6 | GO:1905653 | GO:BP | positive regulation of artery morphogenesis |
| L15 | 0.025860806 | 5 | 204 | 3 | 0.014705882 | 0.6 | GO:0003190 | GO:BP | atrioventricular valve formation |
| L15 | 0.025860806 | 5 | 204 | 3 | 0.014705882 | 0.6 | GO:1905651 | GO:BP | regulation of artery morphogenesis |
| L15 | 0.027406732 | 46 | 204 | 6 | 0.029411765 | 0.130434783 | GO:0048286 | GO:BP | lung alveolus development |
| L15 | 0.029998752 | 476 | 204 | 18 | 0.088235294 | 0.037815126 | GO:0042325 | GO:BP | regulation of phosphorylation |
| L15 | 0.029998752 | 476 | 204 | 18 | 0.088235294 | 0.037815126 | GO:0007423 | GO:BP | sensory organ development |
| L15 | 0.03053477 | 156 | 204 | 10 | 0.049019608 | 0.064102564 | GO:0051056 | GO:BP | regulation of small GTPase mediated signal transduction |

| | | | | | | | | | |
|---|---|---|---|---|---|---|---|---|---|
| L15 | 0.034533839 | 307 | 204 | 14 | 0.068627451 | 0.045602606 | GO:0001654 | GO:BP | eye development |
| L15 | 0.037103567 | 309 | 204 | 14 | 0.068627451 | 0.045307443 | GO:0150063 | GO:BP | visual system development |
| L15 | 0.03970241 | 49 | 204 | 6 | 0.029411765 | 0.12244898 | GO:0003229 | GO:BP | ventricular cardiac muscle tissue development |
| L15 | 0.040811552 | 196 | 204 | 11 | 0.053921569 | 0.056122449 | GO:0060541 | GO:BP | respiratory system development |
| L15 | 0.042752736 | 313 | 204 | 14 | 0.068627451 | 0.044728435 | GO:0048880 | GO:BP | sensory system development |
| L15 | 0.043839523 | 2466 | 204 | 52 | 0.254901961 | 0.02108678 | GO:0051252 | GO:BP | regulation of RNA metabolic process |
| L15 | 0.0488499 | 31 | 204 | 5 | 0.024509804 | 0.161290323 | GO:0030513 | GO:BP | positive regulation of BMP signaling pathway |
| L15 | 0.049703204 | 1177 | 204 | 31 | 0.151960784 | 0.026338148 | GO:0045893 | GO:BP | positive regulation of DNA-templated transcription |
| L15 | 7.45E-07 | 3202 | 187 | 72 | 0.385026738 | 0.022485946 | GO:0071944 | GO:CC | cell periphery |
| L15 | 1.05E-05 | 2917 | 187 | 65 | 0.347593583 | 0.022283168 | GO:0005886 | GO:CC | plasma membrane |
| L15 | 0.001054989 | 6409 | 187 | 105 | 0.561497326 | 0.016383211 | GO:0016020 | GO:CC | membrane |
| L15 | 0.00781463 | 73 | 187 | 7 | 0.037433155 | 0.095890411 | GO:0005912 | GO:CC | adherens junction |
| L15 | 0.013086113 | 892 | 187 | 25 | 0.13368984 | 0.028026906 | GO:0005794 | GO:CC | Golgi apparatus |
| L15 | 0.01551956 | 385 | 187 | 15 | 0.080213904 | 0.038961039 | GO:0005667 | GO:CC | transcription regulator complex |
| L15 | 0.018070402 | 259 | 187 | 12 | 0.064171123 | 0.046332046 | GO:0043235 | GO:CC | receptor complex |
| L15 | 0.018659895 | 536 | 187 | 18 | 0.096256684 | 0.03358209 | GO:0009986 | GO:CC | cell surface |
| L15 | 0.01948202 | 7299 | 187 | 111 | 0.593582888 | 0.015207563 | GO:0005737 | GO:CC | cytoplasm |
| L15 | 3.03E-05 | 20 | 198 | 6 | 0.03030303 | 0.3 | GO:0005160 | GO:MF | transforming growth factor beta receptor binding |
| L15 | 8.14E-05 | 1104 | 198 | 34 | 0.171717172 | 0.030797101 | GO:0044877 | GO:MF | protein-containing complex binding |
| L15 | 0.000221135 | 7099 | 198 | 118 | 0.595959596 | 0.016622059 | GO:0005515 | GO:MF | protein binding |
| L15 | 0.000451551 | 12006 | 198 | 170 | 0.858585859 | 0.014159587 | GO:0005488 | GO:MF | binding |
| L15 | 0.000541385 | 1369 | 198 | 37 | 0.186868687 | 0.027027027 | GO:0098772 | GO:MF | molecular function regulator activity |
| L15 | 0.000812359 | 9 | 198 | 4 | 0.02020202 | 0.444444444 | GO:0034713 | GO:MF | type I transforming growth factor beta receptor binding |
| L15 | 0.00444203 | 13 | 198 | 4 | 0.02020202 | 0.307692308 | GO:0070411 | GO:MF | I-SMAD binding |
| L15 | 0.005911225 | 95 | 198 | 8 | 0.04040404 | 0.084210526 | GO:0019955 | GO:MF | cytokine binding |
| L15 | 0.006151304 | 28 | 198 | 5 | 0.025252525 | 0.178571429 | GO:0015026 | GO:MF | coreceptor activity |
| L15 | 0.007361977 | 29 | 198 | 5 | 0.025252525 | 0.172413793 | GO:0017046 | GO:MF | peptide hormone binding |
| L15 | 0.00860671 | 100 | 198 | 8 | 0.04040404 | 0.08 | GO:0019838 | GO:MF | growth factor binding |
| L15 | 0.01292951 | 137 | 198 | 9 | 0.045454545 | 0.065693431 | GO:0042277 | GO:MF | peptide binding |
| L15 | 0.013456884 | 1455 | 198 | 35 | 0.176767677 | 0.024054983 | GO:0019899 | GO:MF | enzyme binding |
| L15 | 0.015062083 | 174 | 198 | 10 | 0.050505051 | 0.057471264 | GO:0033218 | GO:MF | amide binding |
| L15 | 0.016443892 | 34 | 198 | 5 | 0.025252525 | 0.147058824 | GO:0001223 | GO:MF | transcription coactivator binding |
| L15 | 0.017674636 | 889 | 198 | 25 | 0.126262626 | 0.028121485 | GO:0030234 | GO:MF | enzyme regulator activity |
| L15 | 0.019559688 | 1007 | 198 | 27 | 0.136363636 | 0.026812314 | GO:0005102 | GO:MF | signaling receptor binding |
| L15 | 0.019647225 | 7 | 198 | 3 | 0.015151515 | 0.428571429 | GO:0070697 | GO:MF | activin receptor binding |
| L15 | 0.027727956 | 1570 | 198 | 36 | 0.181818182 | 0.022929936 | GO:0042802 | GO:MF | identical protein binding |
| L15 | 0.028212692 | 20 | 198 | 4 | 0.02020202 | 0.2 | GO:0050431 | GO:MF | transforming growth factor beta binding |
| L15 | 0.029299638 | 544 | 198 | 18 | 0.090909091 | 0.033088235 | GO:0042803 | GO:MF | protein homodimerization activity |
| L15 | 0.03487 | 63 | 198 | 6 | 0.03030303 | 0.095238095 | GO:0019199 | GO:MF | transmembrane receptor protein kinase activity |
| L15 | 0.038536803 | 3802 | 198 | 68 | 0.343434343 | 0.017885324 | GO:0036094 | GO:MF | small molecule binding |
| L15 | 0.041570764 | 65 | 198 | 6 | 0.03030303 | 0.092307692 | GO:0046332 | GO:MF | SMAD binding |

| | | | | | | | | | |
|---|---|---|---|---|---|---|---|---|---|
| L15 | 0.049786841 | 2 | 198 | 2 | 0.01010101 | 1 | GO:0050659 | GO:MF | N-acetylgalactosamine 4-sulfate 6-O-sulfotransferase activity |
| L15 | 5.40E-05 | 4 | 70 | 4 | 0.057142857 | 1 | HP:0001694 | HP | Right-to-left shunt |
| L15 | 0.001830728 | 7 | 70 | 4 | 0.057142857 | 0.571428571 | HP:0004296 | HP | Abnormal gastrointestinal vascular morphology |
| L15 | 0.002751514 | 16 | 70 | 5 | 0.071428571 | 0.3125 | HP:0100761 | HP | Visceral angiomatosis |
| L15 | 0.004046944 | 3 | 70 | 3 | 0.042857143 | 1 | HP:0000471 | HP | Gastrointestinal angiodysplasia |
| L15 | 0.008634981 | 360 | 70 | 17 | 0.242857143 | 0.047222222 | HP:0011028 | HP | Abnormality of blood circulation |
| L15 | 0.012142819 | 21 | 70 | 5 | 0.071428571 | 0.238095238 | HP:0007461 | HP | Hemangiomatosis |
| L15 | 0.015818068 | 56 | 70 | 7 | 0.1 | 0.125 | HP:0100026 | HP | Arteriovenous malformation |
| L15 | 0.016025802 | 4 | 70 | 3 | 0.042857143 | 0.75 | HP:0000227 | HP | Tongue telangiectasia |
| L15 | 0.016025802 | 4 | 70 | 3 | 0.042857143 | 0.75 | HP:0000434 | HP | Nasal mucosa telangiectasia |
| L15 | 0.016025802 | 4 | 70 | 3 | 0.042857143 | 0.75 | HP:0006574 | HP | Hepatic arteriovenous malformation |
| L15 | 0.028137178 | 41 | 70 | 6 | 0.085714286 | 0.146341463 | HP:0002647 | HP | Aortic dissection |
| L15 | 0.028137178 | 41 | 70 | 6 | 0.085714286 | 0.146341463 | HP:0001048 | HP | Cavernous hemangioma |
| L15 | 0.035104077 | 13 | 70 | 4 | 0.057142857 | 0.307692308 | HP:0001693 | HP | Cardiac shunt |
| L15 | 0.039663826 | 5 | 70 | 3 | 0.042857143 | 0.6 | HP:0002629 | HP | Gastrointestinal arteriovenous malformation |
| L15 | 0.039663826 | 5 | 70 | 3 | 0.042857143 | 0.6 | HP:0006548 | HP | Pulmonary arteriovenous malformation |
| L16 | 0.00038718 | 2140 | 75 | 27 | 0.36 | 0.01261682 | GO:0051239 | GO:BP | regulation of multicellular organismal process |
| L16 | 0.005923 | 5110 | 75 | 42 | 0.56 | 0.00821918 | GO:0032501 | GO:BP | multicellular organismal process |
| L16 | 0.01861999 | 1158 | 75 | 17 | 0.22666667 | 0.01468048 | GO:0051240 | GO:BP | positive regulation of multicellular organismal process |
| L16 | 0.04883961 | 756 | 75 | 13 | 0.17333333 | 0.01719577 | GO:0002684 | GO:BP | positive regulation of immune system process |
| L16 | 0.01931699 | 2917 | 69 | 26 | 0.37681159 | 0.00891327 | GO:0005886 | GO:CC | plasma membrane |
| L16 | 0.01668439 | 199 | 29 | 8 | 0.27586207 | 0.04020101 | HP:0002829 | HP | Arthralgia |
| L21 | 6.96E-11 | 2754 | 142 | 61 | 0.429577465 | 0.022149601 | GO:0048731 | GO:BP | system development |
| L21 | 3.26E-10 | 4492 | 142 | 79 | 0.556338028 | 0.017586821 | GO:0032502 | GO:BP | developmental process |
| L21 | 1.24E-09 | 4205 | 142 | 75 | 0.528169014 | 0.01783591 | GO:0048856 | GO:BP | anatomical structure development |
| L21 | 2.11E-09 | 3317 | 142 | 65 | 0.457746479 | 0.019596021 | GO:0007275 | GO:BP | multicellular organism development |
| L21 | 1.17E-06 | 5110 | 142 | 78 | 0.549295775 | 0.015264188 | GO:0032501 | GO:BP | multicellular organismal process |
| L21 | 4.94E-06 | 580 | 142 | 22 | 0.154929577 | 0.037931034 | GO:0001944 | GO:BP | vasculature development |
| L21 | 1.08E-05 | 551 | 142 | 21 | 0.147887324 | 0.038112523 | GO:0001568 | GO:BP | blood vessel development |
| L21 | 1.53E-05 | 853 | 142 | 26 | 0.183098592 | 0.030480657 | GO:0072359 | GO:BP | circulatory system development |
| L21 | 2.15E-05 | 2183 | 142 | 44 | 0.309859155 | 0.020155749 | GO:0048513 | GO:BP | animal organ development |
| L21 | 3.65E-05 | 1977 | 142 | 41 | 0.288732394 | 0.020738493 | GO:0009653 | GO:BP | anatomical structure morphogenesis |
| L21 | 0.000387342 | 1072 | 142 | 27 | 0.190140845 | 0.025186567 | GO:0016477 | GO:BP | cell migration |
| L21 | 0.000650898 | 2897 | 142 | 49 | 0.345070423 | 0.016914049 | GO:0030154 | GO:BP | cell differentiation |
| L21 | 0.000650898 | 2897 | 142 | 49 | 0.345070423 | 0.016914049 | GO:0048869 | GO:BP | cellular developmental process |
| L21 | 0.001406076 | 1520 | 142 | 32 | 0.225352113 | 0.021052632 | GO:0009888 | GO:BP | tissue development |
| L21 | 0.003126898 | 1577 | 142 | 32 | 0.225352113 | 0.020291693 | GO:0007399 | GO:BP | nervous system development |
| L21 | 0.00388989 | 103 | 142 | 8 | 0.056338028 | 0.077669903 | GO:0003158 | GO:BP | endothelium development |
| L21 | 0.005687065 | 1232 | 142 | 27 | 0.190140845 | 0.021915584 | GO:0048870 | GO:BP | cell motility |
| L21 | 0.006556197 | 51 | 142 | 6 | 0.042253521 | 0.117647059 | GO:0045123 | GO:BP | cellular extravasation |
| L21 | 0.007658336 | 387 | 142 | 14 | 0.098591549 | 0.036175711 | GO:0001525 | GO:BP | angiogenesis |

| | | | | | | | | | |
|---|---|---|---|---|---|---|---|---|---|
| L21 | 0.008986142 | 5 | 142 | 3 | 0.021126761 | 0.6 | GO:0035696 | GO:BP | monocyte extravasation |
| L21 | 0.010552888 | 906 | 142 | 22 | 0.154929577 | 0.024282561 | GO:0048646 | GO:BP | anatomical structure formation involved in morphogenesis |
| L21 | 0.015673623 | 470 | 142 | 15 | 0.105633803 | 0.031914894 | GO:0048514 | GO:BP | blood vessel morphogenesis |
| L21 | 0.017123258 | 2401 | 142 | 40 | 0.281690141 | 0.016659725 | GO:0023051 | GO:BP | regulation of signaling |
| L21 | 0.01747058 | 2403 | 142 | 40 | 0.281690141 | 0.016645859 | GO:0010646 | GO:BP | regulation of cell communication |
| L21 | 0.032434346 | 500 | 142 | 15 | 0.105633803 | 0.03 | GO:0034330 | GO:BP | cell junction organization |
| L21 | 0.036016405 | 1859 | 142 | 33 | 0.232394366 | 0.017751479 | GO:0007166 | GO:BP | cell surface receptor signaling pathway |
| L21 | 0.045575755 | 43 | 142 | 5 | 0.035211268 | 0.11627907 | GO:0051785 | GO:BP | positive regulation of nuclear division |
| L21 | 0.001796641 | 16 | 132 | 4 | 0.03030303 | 0.25 | GO:0098644 | GO:CC | complex of collagen trimers |
| L21 | 0.002629396 | 6 | 132 | 3 | 0.022727273 | 0.5 | GO:0098642 | GO:CC | network-forming collagen trimer |
| L21 | 0.002629396 | 6 | 132 | 3 | 0.022727273 | 0.5 | GO:0098645 | GO:CC | collagen network |
| L21 | 0.002629396 | 6 | 132 | 3 | 0.022727273 | 0.5 | GO:0098651 | GO:CC | basement membrane collagen trimer |
| L21 | 0.002629396 | 6 | 132 | 3 | 0.022727273 | 0.5 | GO:0005587 | GO:CC | collagen type IV trimer |
| L21 | 0.004324563 | 138 | 132 | 8 | 0.060606061 | 0.057971014 | GO:0062023 | GO:CC | collagen-containing extracellular matrix |
| L21 | 0.006952553 | 22 | 132 | 4 | 0.03030303 | 0.181818182 | GO:0005581 | GO:CC | collagen trimer |
| L21 | 0.010356265 | 1214 | 132 | 24 | 0.181818182 | 0.019769357 | GO:0030054 | GO:CC | cell junction |
| L21 | 0.026426958 | 228 | 132 | 9 | 0.068181818 | 0.039473684 | GO:0031012 | GO:CC | extracellular matrix |
| L21 | 0.027321141 | 229 | 132 | 9 | 0.068181818 | 0.03930131 | GO:0030312 | GO:CC | external encapsulating structure |
| L21 | 0.034716835 | 61 | 132 | 5 | 0.037878788 | 0.081967213 | GO:0005604 | GO:CC | basement membrane |
| L21 | 0.0033377 | 32 | 136 | 5 | 0.036764706 | 0.15625 | GO:0005201 | GO:MF | extracellular matrix structural constituent |
| L21 | 0.003436128 | 3802 | 136 | 54 | 0.397058824 | 0.014203051 | GO:0036094 | GO:MF | small molecule binding |
| L21 | 0.003553897 | 3602 | 136 | 52 | 0.382352941 | 0.014436424 | GO:0043167 | GO:MF | ion binding |
| L21 | 0.010228315 | 100 | 136 | 7 | 0.051470588 | 0.07 | GO:0019838 | GO:MF | growth factor binding |
| L21 | 0.012012302 | 582 | 136 | 16 | 0.117647059 | 0.027491409 | GO:0005509 | GO:MF | calcium ion binding |
| L21 | 0.037403898 | 1851 | 136 | 31 | 0.227941176 | 0.016747704 | GO:0046872 | GO:MF | metal ion binding |
| L25 | 0.02894431 | 4 | 73 | 2 | 0.02739726 | 0.5 | GO:0005655 | GO:CC | nucleolar ribonuclease P complex |
| L25 | 0.02894431 | 4 | 73 | 2 | 0.02739726 | 0.5 | GO:0032777 | GO:CC | piccolo histone acetyltransferase complex |
| L26 | 0.040629231 | 368 | 71 | 9 | 0.126760563 | 0.024456522 | GO:0034655 | GO:BP | nucleobase-containing compound catabolic process |
| L26 | 0.002388593 | 3725 | 68 | 32 | 0.470588235 | 0.008590604 | GO:0031981 | GO:CC | nuclear lumen |
| L26 | 0.003390258 | 3979 | 68 | 33 | 0.485294118 | 0.008293541 | GO:0031974 | GO:CC | membrane-enclosed lumen |
| L26 | 0.003390258 | 3979 | 68 | 33 | 0.485294118 | 0.008293541 | GO:0043233 | GO:CC | organelle lumen |
| L26 | 0.003390258 | 3979 | 68 | 33 | 0.485294118 | 0.008293541 | GO:0070013 | GO:CC | intracellular organelle lumen |
| L26 | 0.008102838 | 9306 | 68 | 55 | 0.808823529 | 0.005910165 | GO:0043231 | GO:CC | intracellular membrane-bounded organelle |
| L26 | 0.011467494 | 9673 | 68 | 56 | 0.823529412 | 0.00578931 | GO:0043227 | GO:CC | membrane-bounded organelle |
| L26 | 0.014340652 | 11922 | 68 | 63 | 0.926470588 | 0.005284348 | GO:0005622 | GO:CC | intracellular anatomical structure |
| L26 | 0.039823876 | 125 | 68 | 5 | 0.073529412 | 0.04 | GO:0001650 | GO:CC | fibrillar center |
| L29 | 0.00178602 | 2754 | 79 | 31 | 0.39240506 | 0.01125635 | GO:0048731 | GO:BP | system development |
| L29 | 0.00203953 | 5 | 79 | 3 | 0.03797468 | 0.6 | GO:0072143 | GO:BP | mesangial cell development |
| L29 | 0.00306744 | 4492 | 79 | 41 | 0.51898734 | 0.00912734 | GO:0032502 | GO:BP | developmental process |
| L29 | 0.0040653 | 6 | 79 | 3 | 0.03797468 | 0.5 | GO:0072007 | GO:BP | mesangial cell differentiation |
| L29 | 0.0047026 | 4205 | 79 | 39 | 0.49367089 | 0.00927467 | GO:0048856 | GO:BP | anatomical structure development |

| | | | | | | | | | |
|---|---|---|---|---|---|---|---|---|---|
| L29 | 0.01105103 | 3317 | 79 | 33 | 0.41772152 | 0.00994875 | GO:0007275 | GO:BP | multicellular organism development |
| L29 | 0.01690203 | 9 | 79 | 3 | 0.03797468 | 0.33333333 | GO:1904238 | GO:BP | pericyte cell differentiation |
| L29 | 0.02051383 | 463 | 79 | 11 | 0.13924051 | 0.0237581 | GO:0061061 | GO:BP | muscle structure development |
| L29 | 0.03196072 | 581 | 79 | 12 | 0.15189873 | 0.02065404 | GO:0022603 | GO:BP | regulation of anatomical structure morphogenesis |
| L29 | 0.04502488 | 2 | 79 | 2 | 0.02531646 | 1 | GO:0003275 | GO:BP | apoptotic process involved in outflow tract morphogenesis |
| L29 | 0.04502488 | 2 | 79 | 2 | 0.02531646 | 1 | GO:1902256 | GO:BP | regulation of apoptotic process involved in outflow tract morphogenesis |
| L29 | 0.04502488 | 2 | 79 | 2 | 0.02531646 | 1 | GO:1902257 | GO:BP | negative regulation of apoptotic process involved in outflow tract morphogenesis |
| L29 | 0.04859381 | 36 | 79 | 4 | 0.05063291 | 0.11111111 | GO:0035850 | GO:BP | epithelial cell differentiation involved in kidney development |
| L29 | 0.02777931 | 1532 | 73 | 18 | 0.24657534 | 0.01174935 | GO:0005856 | GO:CC | cytoskeleton |
| L29 | 0.01986117 | 7099 | 84 | 54 | 0.64285714 | 0.00760671 | GO:0005515 | GO:MF | protein binding |
| L31 | 0.000242776 | 2917 | 63 | 28 | 0.444444444 | 0.009598903 | GO:0005886 | GO:CC | plasma membrane |
| L31 | 0.000470006 | 3202 | 63 | 29 | 0.46031746 | 0.009056839 | GO:0071944 | GO:CC | cell periphery |
| L31 | 0.010058504 | 3 | 63 | 2 | 0.031746032 | 0.666666667 | GO:0045179 | GO:CC | apical cortex |
| L31 | 0.016762222 | 2627 | 63 | 23 | 0.365079365 | 0.008755234 | GO:0005829 | GO:CC | cytosol |
| L31 | 0.020796289 | 275 | 63 | 7 | 0.111111111 | 0.025454545 | GO:0045177 | GO:CC | apical part of cell |
| L31 | 0.004799563 | 182 | 69 | 7 | 0.101449275 | 0.038461538 | GO:0005085 | GO:MF | guanyl-nucleotide exchange factor activity |
| L35 | 9.32E-07 | 76 | 42 | 7 | 0.166666667 | 0.092105263 | GO:0042752 | GO:BP | regulation of circadian rhythm |
| L35 | 1.87E-06 | 136 | 42 | 8 | 0.19047619 | 0.058823529 | GO:0007623 | GO:BP | circadian rhythm |
| L35 | 1.44E-05 | 176 | 42 | 8 | 0.19047619 | 0.045454545 | GO:0048511 | GO:BP | rhythmic process |
| L35 | 0.000201645 | 50 | 42 | 5 | 0.119047619 | 0.1 | GO:0032922 | GO:BP | circadian regulation of gene expression |
| L35 | 0.000392819 | 23 | 42 | 4 | 0.095238095 | 0.173913043 | GO:0048512 | GO:BP | circadian behavior |
| L35 | 0.000559151 | 25 | 42 | 4 | 0.095238095 | 0.16 | GO:0007622 | GO:BP | rhythmic behavior |
| L35 | 0.013189491 | 17 | 42 | 3 | 0.071428571 | 0.176470588 | GO:0009648 | GO:BP | photoperiodism |
| L35 | 0.013189491 | 17 | 42 | 3 | 0.071428571 | 0.176470588 | GO:0043153 | GO:BP | entrainment of circadian clock by photoperiod |
| L35 | 0.018730021 | 19 | 42 | 3 | 0.071428571 | 0.157894737 | GO:0009649 | GO:BP | entrainment of circadian clock |
| L35 | 0.01382813 | 7 | 31 | 2 | 0.064516129 | 0.285714286 | GO:0033180 | GO:CC | proton-transporting V-type ATPase, V1 domain |
| L35 | 0.021295291 | 5 | 41 | 2 | 0.048780488 | 0.4 | GO:0043426 | GO:MF | MRF binding |
| L35 | 0.039514351 | 38 | 41 | 3 | 0.073170732 | 0.078947368 | GO:0070888 | GO:MF | E-box binding |
| L36 | 0.02432914 | 3 | 39 | 2 | 0.05128205 | 0.66666667 | GO:0018344 | GO:BP | protein geranylgeranylation |
| L36 | 0.02707694 | 364 | 39 | 7 | 0.17948718 | 0.01923077 | GO:0002253 | GO:BP | activation of immune response |
| L36 | 0.04858719 | 4 | 39 | 2 | 0.05128205 | 0.5 | GO:0038001 | GO:BP | paracrine signaling |
| L38 | 0.010517463 | 226 | 35 | 6 | 0.171428571 | 0.026548673 | GO:0030198 | GO:BP | extracellular matrix organization |
| L38 | 0.010517463 | 226 | 35 | 6 | 0.171428571 | 0.026548673 | GO:0045229 | GO:BP | external encapsulating structure organization |
| L38 | 0.011062184 | 228 | 35 | 6 | 0.171428571 | 0.026315789 | GO:0043062 | GO:BP | extracellular structure organization |
| L38 | 0.031744298 | 1116 | 31 | 9 | 0.290322581 | 0.008064516 | GO:0005576 | GO:CC | extracellular region |
| L38 | 0.049034532 | 3 | 18 | 2 | 0.111111111 | 0.666666667 | HP:0012234 | HP | Agranulocytosis |

**Table S10.** Right ventricle module expression ANOVA results.

| Module | population | treatment | population*treatment |
|---|---|---|---|
| RV01 | 0.998 | 1 | 0.4789 |
| RV02 | 0.9995 | 0.9995 | 0.4813 |
| **RV03** | **0** | 0.1132 | 0.1938 |
| **RV04** | **0** | 0.8039 | 0.8303 |
| **RV05** | **4.00E-04** | 0.3045 | 0.8303 |
| RV06 | 0.2406 | 0.3188 | 0.8303 |
| **RV07** | **0** | 0.9995 | 0.8252 |
| **RV08** | **8.00E-04** | 0.8019 | 0.8303 |
| RV09 | 0.9995 | 0.1132 | 0.5283 |
| **RV10** | **0** | 0.9217 | 0.1739 |
| RV11 | 0.1528 | 0.1396 | 0.1009 |
| RV12 | 0.0944 | 0.9769 | 0.6082 |
| RV13 | 0.125 | 0.1938 | 0.9995 |
| RV14 | 1 | 0.2205 | 0.9995 |
| **RV15** | **0** | 0.9769 | 0.2006 |
| RV16 | 0.4697 | 0.1178 | 0.6687 |
| RV17 | 0.7395 | 0.1674 | 0.4986 |
| RV18 | 0.1328 | 0.059 | 0.9769 |
| RV19 | 0.1938 | 0.672 | 0.6784 |
| RV20 | 0.5693 | 0.8547 | 0.1567 |
| RV21 | 0.7222 | 0.9995 | 0.1132 |
| RV22 | 0.4478 | 1 | 0.8303 |
| RV23 | 0.059 | 0.6082 | 0.9769 |
| RV24 | 0.9769 | 0.1528 | 0.5314 |
| **RV25** | 0.6687 | **0.0028** | **0.0572** |
| RV26 | 0.7179 | 0.9769 | 0.9995 |
| RV27 | 0.1202 | 0.8303 | 0.8303 |
| RV28 | 0.7096 | 0.1114 | 0.2897 |
| RV29 | 0.0832 | 0.9769 | 0.8039 |
| RV30 | 0.5534 | 0.6839 | 0.9995 |
| RV31 | 0.1674 | 0.7406 | 0.8204 |
| **RV32** | **0** | 0.2687 | 0.1117 |
| **RV33** | **0** | 0.9995 | 0.3188 |
| RV34 | 0.7197 | 0.5283 | 0.9769 |
| RV35 | 0.1739 | 0.9995 | 0.8303 |
| RV36 | 1 | 0.9215 | 0.1519 |
| **RV37** | **0.0028** | 0.8039 | 0.6085 |

**Table S11.** Right ventricle module association with relative right ventricle mass.

| Module | correlation | p-value | FDR p-value |
|---|---|---|---|
| RV03 | -0.3630 | 0.0181 | 0.2265 |
| RV32 | 0.3603 | 0.0191 | 0.2265 |
| RV04 | 0.3488 | 0.0236 | 0.2265 |
| RV15 | 0.3482 | 0.0238 | 0.2265 |
| RV06 | -0.2831 | 0.0693 | 0.4390 |
| RV29 | -0.2532 | 0.1057 | 0.5346 |
| RV25 | 0.2485 | 0.1125 | 0.5346 |
| RV30 | 0.2259 | 0.1503 | 0.6001 |
| RV31 | 0.2216 | 0.1584 | 0.6001 |
| RV07 | 0.2139 | 0.1737 | 0.6001 |
| RV27 | 0.2052 | 0.1924 | 0.6092 |
| RV26 | 0.1929 | 0.2209 | 0.6447 |
| RV22 | -0.1863 | 0.2375 | 0.6447 |
| RV23 | 0.1681 | 0.2872 | 0.7275 |
| RV08 | 0.1486 | 0.3478 | 0.7718 |
| RV19 | -0.1479 | 0.3498 | 0.7718 |
| RV17 | 0.1432 | 0.3656 | 0.7718 |
| RV02 | -0.1333 | 0.4001 | 0.7863 |
| RV12 | -0.1249 | 0.4306 | 0.7863 |
| RV10 | 0.1239 | 0.4345 | 0.7863 |
| RV05 | -0.1055 | 0.5062 | 0.8743 |
| RV34 | -0.0869 | 0.5844 | 0.9419 |
| RV20 | -0.0845 | 0.5949 | 0.9419 |
| RV24 | -0.0589 | 0.7109 | 0.9930 |
| RV13 | 0.0529 | 0.7391 | 0.9930 |
| RV11 | 0.0517 | 0.7449 | 0.9930 |
| RV36 | 0.0473 | 0.7663 | 0.9930 |
| RV18 | 0.0423 | 0.7904 | 0.9930 |
| RV21 | -0.0330 | 0.8359 | 0.9930 |
| RV01 | 0.0291 | 0.8550 | 0.9930 |
| RV28 | -0.0267 | 0.8669 | 0.9930 |
| RV33 | -0.0247 | 0.8765 | 0.9930 |
| RV14 | -0.0119 | 0.9404 | 0.9930 |
| RV16 | -0.0064 | 0.9680 | 0.9930 |
| RV35 | 0.0053 | 0.9734 | 0.9930 |
| RV09 | 0.0016 | 0.9921 | 0.9930 |
| RV37 | -0.0014 | 0.9930 | 0.9930 |

**Table S12.** Right ventricle GO results of significant terms.

| query | p_value | term_size | query_size | intersection_size | precision | recall | term_id | source | term_name |
|---|---|---|---|---|---|---|---|---|---|
| RV3 | 0.004263863 | 923 | 524 | 55 | 0.104961832 | 0.059588299 | GO:0006629 | GO:BP | lipid metabolic process |
| RV3 | 0.017325775 | 1188 | 524 | 64 | 0.122137405 | 0.053872054 | GO:0044281 | GO:BP | small molecule metabolic process |
| RV3 | 4.05E-09 | 7299 | 518 | 308 | 0.594594595 | 0.042197561 | GO:0005737 | GO:CC | cytoplasm |
| RV3 | 7.68E-05 | 1013 | 518 | 63 | 0.121621622 | 0.06219151 | GO:0005783 | GO:CC | endoplasmic reticulum |
| RV3 | 0.000540315 | 2533 | 518 | 121 | 0.233590734 | 0.047769443 | GO:0012505 | GO:CC | endomembrane system |
| RV3 | 0.013135425 | 125 | 518 | 14 | 0.027027027 | 0.112 | GO:0001650 | GO:CC | fibrillar center |
| RV3 | 0.018895075 | 9673 | 518 | 351 | 0.677606178 | 0.036286571 | GO:0043227 | GO:CC | membrane-bounded organelle |
| RV3 | 0.022234034 | 892 | 518 | 50 | 0.096525097 | 0.056053812 | GO:0005794 | GO:CC | Golgi apparatus |
| RV3 | 0.044734134 | 11922 | 518 | 416 | 0.803088803 | 0.034893474 | GO:0005622 | GO:CC | intracellular anatomical structure |
| RV3 | 3.45E-07 | 5159 | 549 | 235 | 0.428051002 | 0.045551463 | GO:0003824 | GO:MF | catalytic activity |
| RV3 | 0.010340871 | 7099 | 549 | 279 | 0.508196721 | 0.03930131 | GO:0005515 | GO:MF | protein binding |
| RV3 | 0.026138768 | 2001 | 549 | 96 | 0.174863388 | 0.047976012 | GO:0016740 | GO:MF | transferase activity |
| RV4 | 0.009100887 | 2437 | 525 | 113 | 0.215238095 | 0.046368486 | GO:0006996 | GO:BP | organelle organization |
| RV4 | 0.022746921 | 2349 | 525 | 108 | 0.205714286 | 0.045977011 | GO:0051641 | GO:BP | cellular localization |
| RV4 | 0.029756666 | 2065 | 525 | 97 | 0.184761905 | 0.046973366 | GO:0043412 | GO:BP | macromolecule modification |
| RV4 | 1.71E-16 | 7299 | 520 | 333 | 0.640384615 | 0.045622688 | GO:0005737 | GO:CC | cytoplasm |
| RV4 | 0.000174104 | 2627 | 520 | 127 | 0.244230769 | 0.048344119 | GO:0005829 | GO:CC | cytosol |
| RV4 | 0.000296773 | 11922 | 520 | 428 | 0.823076923 | 0.035900017 | GO:0005622 | GO:CC | intracellular anatomical structure |
| RV4 | 0.002152389 | 1163 | 520 | 65 | 0.125 | 0.05588994 | GO:0005739 | GO:CC | mitochondrion |
| RV4 | 0.004065347 | 1532 | 520 | 79 | 0.151923077 | 0.05156658 | GO:0005856 | GO:CC | cytoskeleton |
| RV4 | 0.02007146 | 198 | 520 | 18 | 0.034615385 | 0.090909091 | GO:0005759 | GO:CC | mitochondrial matrix |
| RV4 | 0.026588313 | 11014 | 520 | 392 | 0.753846154 | 0.035591066 | GO:0043226 | GO:CC | organelle |
| RV4 | 0.030043256 | 9673 | 520 | 351 | 0.675 | 0.036286571 | GO:0043227 | GO:CC | membrane-bounded organelle |

| | | | | | | | | | |
|---|---|---|---|---|---|---|---|---|---|
| RV4 | 0.044996971 | 610 | 520 | 37 | 0.071153846 | 0.060655738 | GO:0005815 | GO:CC | microtubule organizing center |
| RV4 | 3.33E-07 | 5159 | 546 | 234 | 0.428571429 | 0.045357627 | GO:0003824 | GO:MF | catalytic activity |
| RV4 | 6.71E-06 | 192 | 546 | 24 | 0.043956044 | 0.125 | GO:0008168 | GO:MF | methyltransferase activity |
| RV4 | 1.81E-05 | 202 | 546 | 24 | 0.043956044 | 0.118811881 | GO:0016741 | GO:MF | transferase activity, transferring one-carbon groups |
| RV4 | 1.95E-05 | 173 | 546 | 22 | 0.04029304 | 0.12716763 | GO:0008757 | GO:MF | S-adenosylmethionine-dependent methyltransferase activity |
| RV4 | 4.35E-05 | 2001 | 546 | 107 | 0.195970696 | 0.053473263 | GO:0016740 | GO:MF | transferase activity |
| RV4 | 0.013168378 | 7099 | 546 | 277 | 0.507326007 | 0.03901958 | GO:0005515 | GO:MF | protein binding |
| RV4 | 0.013506076 | 1570 | 546 | 80 | 0.146520147 | 0.050955414 | GO:0042802 | GO:MF | identical protein binding |
| RV4 | 0.038455692 | 71 | 546 | 10 | 0.018315018 | 0.14084507 | GO:0008170 | GO:MF | N-methyltransferase activity |
| RV4 | 0.009639437 | 2193 | 198 | 117 | 0.590909091 | 0.053351573 | HP:0003808 | HP | Abnormal muscle tone |
| RV4 | 0.034105958 | 2472 | 198 | 126 | 0.636363636 | 0.050970874 | HP:0012758 | HP | Neurodevelopmental delay |
| RV5 | 0.033151856 | 694 | 410 | 36 | 0.087804878 | 0.051873199 | GO:0006259 | GO:BP | DNA metabolic process |
| RV5 | 1.54E-10 | 11922 | 427 | 374 | 0.87587822 | 0.031370575 | GO:0005622 | GO:CC | intracellular anatomical structure |
| RV5 | 2.22E-10 | 2919 | 427 | 137 | 0.320843091 | 0.046933881 | GO:0005654 | GO:CC | nucleoplasm |
| RV5 | 1.92E-08 | 7299 | 427 | 258 | 0.604215457 | 0.035347308 | GO:0005737 | GO:CC | cytoplasm |
| RV5 | 4.25E-07 | 3725 | 427 | 152 | 0.355971897 | 0.040805369 | GO:0031981 | GO:CC | nuclear lumen |
| RV5 | 8.74E-07 | 9673 | 427 | 312 | 0.730679157 | 0.03225473 | GO:0043227 | GO:CC | membrane-bounded organelle |
| RV5 | 7.64E-06 | 3979 | 427 | 155 | 0.362997658 | 0.038954511 | GO:0031974 | GO:CC | membrane-enclosed lumen |
| RV5 | 7.64E-06 | 3979 | 427 | 155 | 0.362997658 | 0.038954511 | GO:0070013 | GO:CC | intracellular organelle lumen |
| RV5 | 7.64E-06 | 3979 | 427 | 155 | 0.362997658 | 0.038954511 | GO:0043233 | GO:CC | organelle lumen |
| RV5 | 1.74E-05 | 9306 | 427 | 298 | 0.697892272 | 0.032022351 | GO:0043231 | GO:CC | intracellular membrane-bounded organelle |
| RV5 | 3.36E-05 | 11014 | 427 | 338 | 0.791569087 | 0.030688215 | GO:0043226 | GO:CC | organelle |
| RV5 | 0.00019573 | 10760 | 427 | 329 | 0.770491803 | 0.030576208 | GO:0043229 | GO:CC | intracellular organelle |
| RV5 | 0.000208961 | 2627 | 427 | 108 | 0.2529274 | 0.041111534 | GO:0005829 | GO:CC | cytosol |
| RV5 | 0.000849875 | 119 | 427 | 14 | 0.032786885 | 0.117647059 | GO:0005681 | GO:CC | spliceosomal complex |
| RV5 | 0.020123925 | 6388 | 427 | 207 | 0.484777518 | 0.032404508 | GO:0005634 | GO:CC | nucleus |

| | | | | | | | | | |
|---|---|---|---|---|---|---|---|---|---|
| RV5 | 0.022198792 | 998 | 427 | 47 | 0.110070258 | 0.047094188 | GO:0005694 | GO:CC | chromosome |
| RV5 | 2.13E-07 | 12006 | 419 | 353 | 0.8424821 | 0.029401966 | GO:0005488 | GO:MF | binding |
| RV5 | 2.62E-05 | 7099 | 419 | 231 | 0.551312649 | 0.032539794 | GO:0005515 | GO:MF | protein binding |
| RV5 | 0.006185448 | 162 | 419 | 15 | 0.035799523 | 0.092592593 | GO:0003729 | GO:MF | mRNA binding |
| RV5 | 0.007437048 | 271 | 419 | 20 | 0.047732697 | 0.073800738 | GO:0061629 | GO:MF | RNA polymerase II-specific DNA-binding transcription factor binding |
| RV5 | 0.014348951 | 1570 | 419 | 65 | 0.155131265 | 0.041401274 | GO:0042802 | GO:MF | identical protein binding |
| RV5 | 0.031843856 | 56 | 419 | 8 | 0.019093079 | 0.142857143 | GO:0003724 | GO:MF | RNA helicase activity |
| RV5 | 0.036246084 | 57 | 419 | 8 | 0.019093079 | 0.140350877 | GO:0008186 | GO:MF | ATP-dependent activity, acting on RNA |
| RV5 | 0.000663771 | 2472 | 145 | 101 | 0.696551724 | 0.040857605 | HP:0012758 | HP | Neurodevelopmental delay |
| RV5 | 0.001240206 | 113 | 145 | 15 | 0.103448276 | 0.132743363 | HP:0002273 | HP | Tetraparesis |
| RV5 | 0.006294887 | 2943 | 145 | 111 | 0.765517241 | 0.037716616 | HP:0012759 | HP | Neurodevelopmental abnormality |
| RV5 | 0.007930043 | 70 | 145 | 11 | 0.075862069 | 0.157142857 | HP:0001285 | HP | Spastic tetraparesis |
| RV5 | 0.011960463 | 152 | 145 | 16 | 0.110344828 | 0.105263158 | HP:0030182 | HP | Tetraplegia/tetraparesis |
| RV5 | 0.014298555 | 1942 | 145 | 82 | 0.565517241 | 0.042224511 | HP:0001263 | HP | Global developmental delay |
| RV5 | 0.015799752 | 3781 | 145 | 130 | 0.896551724 | 0.034382439 | HP:0012638 | HP | Abnormal nervous system physiology |
| RV5 | 0.043535868 | 7 | 145 | 4 | 0.027586207 | 0.571428571 | HP:0003524 | HP | Decreased methionine synthase activity |
| RV5 | 0.049941138 | 1639 | 145 | 71 | 0.489655172 | 0.043319097 | HP:0000240 | HP | Abnormality of skull size |
| RV7 | 0.000671655 | 3856 | 323 | 113 | 0.349845201 | 0.029304979 | GO:0051179 | GO:BP | localization |
| RV7 | 0.001462253 | 1850 | 323 | 65 | 0.20123839 | 0.035135135 | GO:0035556 | GO:BP | intracellular signal transduction |
| RV7 | 0.002036114 | 2401 | 323 | 78 | 0.241486068 | 0.032486464 | GO:0023051 | GO:BP | regulation of signaling |
| RV7 | 0.007273991 | 3311 | 323 | 97 | 0.300309598 | 0.029296285 | GO:0051234 | GO:BP | establishment of localization |
| RV7 | 0.008061208 | 2403 | 323 | 76 | 0.235294118 | 0.031627133 | GO:0010646 | GO:BP | regulation of cell communication |
| RV7 | 0.010723428 | 3073 | 323 | 91 | 0.281733746 | 0.029612756 | GO:0006810 | GO:BP | transport |
| RV7 | 0.012347896 | 2094 | 323 | 68 | 0.210526316 | 0.032473734 | GO:0009966 | GO:BP | regulation of signal transduction |
| RV7 | 0.01515629 | 1859 | 323 | 62 | 0.191950464 | 0.033351264 | GO:0007166 | GO:BP | cell surface receptor signaling pathway |
| RV7 | 0.03204448 | 1225 | 323 | 45 | 0.139318885 | 0.036734694 | GO:0141124 | GO:BP | intracellular signaling cassette |

| | | | | | | | | | |
|---|---|---|---|---|---|---|---|---|---|
| RV7 | 0.037041399 | 581 | 323 | 27 | 0.083591331 | 0.046471601 | GO:0022603 | GO:BP | regulation of anatomical structure morphogenesis |
| RV7 | 0.046802328 | 1244 | 323 | 45 | 0.139318885 | 0.036173633 | GO:0042592 | GO:BP | homeostatic process |
| RV7 | 1.29E-10 | 7299 | 305 | 199 | 0.652459016 | 0.027264009 | GO:0005737 | GO:CC | cytoplasm |
| RV7 | 0.037139096 | 2627 | 305 | 74 | 0.242622951 | 0.028169014 | GO:0005829 | GO:CC | cytosol |
| RV7 | 0.046444635 | 9673 | 305 | 212 | 0.695081967 | 0.021916675 | GO:0043227 | GO:CC | membrane-bounded organelle |
| RV7 | 0.034403313 | 1570 | 334 | 53 | 0.158682635 | 0.033757962 | GO:0042802 | GO:MF | identical protein binding |
| RV7 | 0.000220515 | 238 | 110 | 20 | 0.181818182 | 0.084033613 | HP:0000944 | HP | Abnormal metaphysis morphology |
| RV7 | 0.006539155 | 59 | 110 | 9 | 0.081818182 | 0.152542373 | HP:0003015 | HP | Flared metaphysis |
| RV7 | 0.021194184 | 104 | 110 | 11 | 0.1 | 0.105769231 | HP:0003016 | HP | Metaphyseal widening |
| RV8 | 7.36E-05 | 25 | 241 | 7 | 0.029045643 | 0.28 | GO:0006084 | GO:BP | acetyl-CoA metabolic process |
| RV8 | 0.000981721 | 69 | 241 | 9 | 0.037344398 | 0.130434783 | GO:0034032 | GO:BP | purine nucleoside bisphosphate metabolic process |
| RV8 | 0.000981721 | 69 | 241 | 9 | 0.037344398 | 0.130434783 | GO:0033875 | GO:BP | ribonucleoside bisphosphate metabolic process |
| RV8 | 0.000981721 | 69 | 241 | 9 | 0.037344398 | 0.130434783 | GO:0033865 | GO:BP | nucleoside bisphosphate metabolic process |
| RV8 | 0.001157076 | 52 | 241 | 8 | 0.033195021 | 0.153846154 | GO:0035383 | GO:BP | thioester metabolic process |
| RV8 | 0.001157076 | 52 | 241 | 8 | 0.033195021 | 0.153846154 | GO:0006637 | GO:BP | acyl-CoA metabolic process |
| RV8 | 0.001655605 | 919 | 241 | 33 | 0.136929461 | 0.035908596 | GO:1901565 | GO:BP | organonitrogen compound catabolic process |
| RV8 | 0.002499407 | 1657 | 241 | 48 | 0.199170124 | 0.028968014 | GO:0009056 | GO:BP | catabolic process |
| RV8 | 0.003864578 | 1188 | 241 | 38 | 0.157676349 | 0.031986532 | GO:0044281 | GO:BP | small molecule metabolic process |
| RV8 | 0.006101721 | 30 | 241 | 6 | 0.024896266 | 0.2 | GO:0034033 | GO:BP | purine nucleoside bisphosphate biosynthetic process |
| RV8 | 0.006101721 | 30 | 241 | 6 | 0.024896266 | 0.2 | GO:0033866 | GO:BP | nucleoside bisphosphate biosynthetic process |
| RV8 | 0.006101721 | 30 | 241 | 6 | 0.024896266 | 0.2 | GO:0034030 | GO:BP | ribonucleoside bisphosphate biosynthetic process |
| RV8 | 0.009672447 | 19 | 241 | 5 | 0.020746888 | 0.263157895 | GO:0071616 | GO:BP | acyl-CoA biosynthetic process |
| RV8 | 0.009672447 | 19 | 241 | 5 | 0.020746888 | 0.263157895 | GO:0035384 | GO:BP | thioester biosynthetic process |
| RV8 | 0.017786372 | 1368 | 241 | 40 | 0.165975104 | 0.029239766 | GO:1901575 | GO:BP | organic substance catabolic process |
| RV8 | 0.043442407 | 1025 | 241 | 32 | 0.132780083 | 0.031219512 | GO:0044248 | GO:BP | cellular catabolic process |
| RV8 | 3.78E-07 | 7299 | 231 | 149 | 0.645021645 | 0.020413755 | GO:0005737 | GO:CC | cytoplasm |

| | | | | | | | | | |
|---|---|---|---|---|---|---|---|---|---|
| RV8 | 0.023585473 | 7 | 231 | 3 | 0.012987013 | 0.428571429 | GO:0005967 | GO:CC | mitochondrial pyruvate dehydrogenase complex |
| RV8 | 0.023585473 | 7 | 231 | 3 | 0.012987013 | 0.428571429 | GO:0045254 | GO:CC | pyruvate dehydrogenase complex |
| RV8 | 6.98E-07 | 554 | 256 | 30 | 0.1171875 | 0.054151625 | GO:0008289 | GO:MF | lipid binding |
| RV8 | 0.000111277 | 5159 | 256 | 117 | 0.45703125 | 0.022678814 | GO:0003824 | GO:MF | catalytic activity |
| RV8 | 0.000459437 | 339 | 256 | 19 | 0.07421875 | 0.056047198 | GO:0005543 | GO:MF | phospholipid binding |
| RV8 | 0.01887636 | 3602 | 256 | 82 | 0.3203125 | 0.02276513 | GO:0043167 | GO:MF | ion binding |
| RV8 | 0.024985911 | 3802 | 256 | 85 | 0.33203125 | 0.022356654 | GO:0036094 | GO:MF | small molecule binding |
| RV8 | 0.026465655 | 6 | 256 | 3 | 0.01171875 | 0.5 | GO:0034603 | GO:MF | pyruvate dehydrogenase [NAD(P)+] activity |
| RV8 | 0.026465655 | 6 | 256 | 3 | 0.01171875 | 0.5 | GO:0034604 | GO:MF | pyruvate dehydrogenase (NAD+) activity |
| RV8 | 0.026465655 | 6 | 256 | 3 | 0.01171875 | 0.5 | GO:0004738 | GO:MF | pyruvate dehydrogenase activity |
| RV8 | 0.027071412 | 1943 | 256 | 51 | 0.19921875 | 0.02624807 | GO:0043168 | GO:MF | anion binding |
| RV8 | 0.027498891 | 401 | 96 | 21 | 0.21875 | 0.052369077 | HP:0001941 | HP | Acidosis |
| RV10 | 0.000175418 | 221 | 237 | 16 | 0.067510549 | 0.07239819 | GO:0015980 | GO:BP | energy derivation by oxidation of organic compounds |
| RV10 | 0.000831642 | 161 | 237 | 13 | 0.054852321 | 0.080745342 | GO:0045333 | GO:BP | cellular respiration |
| RV10 | 0.001381256 | 323 | 237 | 18 | 0.075949367 | 0.055727554 | GO:0006091 | GO:BP | generation of precursor metabolites and energy |
| RV10 | 0.010715094 | 33 | 237 | 6 | 0.025316456 | 0.181818182 | GO:0072528 | GO:BP | pyrimidine-containing compound biosynthetic process |
| RV10 | 0.040840512 | 25 | 237 | 5 | 0.021097046 | 0.2 | GO:0006221 | GO:BP | pyrimidine nucleotide biosynthetic process |
| RV10 | 0.041631018 | 61 | 237 | 7 | 0.029535865 | 0.114754098 | GO:1901874 | GO:BP | negative regulation of post-translational protein modification |
| RV10 | 0.044103841 | 1188 | 237 | 35 | 0.147679325 | 0.029461279 | GO:0044281 | GO:BP | small molecule metabolic process |
| RV10 | 0.045468398 | 4430 | 237 | 91 | 0.383966245 | 0.020541761 | GO:1901564 | GO:BP | organonitrogen compound metabolic process |
| RV10 | 2.89E-16 | 1163 | 227 | 59 | 0.259911894 | 0.050730868 | GO:0005739 | GO:CC | mitochondrion |
| RV10 | 3.65E-16 | 7299 | 227 | 167 | 0.735682819 | 0.022879847 | GO:0005737 | GO:CC | cytoplasm |
| RV10 | 1.44E-08 | 11922 | 227 | 206 | 0.907488987 | 0.01727898 | GO:0005622 | GO:CC | intracellular anatomical structure |
| RV10 | 4.92E-06 | 198 | 227 | 16 | 0.070484581 | 0.080808081 | GO:0005759 | GO:CC | mitochondrial matrix |
| RV10 | 6.28E-06 | 9306 | 227 | 170 | 0.748898678 | 0.018267784 | GO:0043231 | GO:CC | intracellular membrane-bounded organelle |
| RV10 | 1.12E-05 | 268 | 227 | 18 | 0.079295154 | 0.067164179 | GO:0005743 | GO:CC | mitochondrial inner membrane |

| | | | | | | | | | |
|---|---|---|---|---|---|---|---|---|---|
| RV10 | 1.94E-05 | 1292 | 227 | 43 | 0.189427313 | 0.033281734 | GO:1902494 | GO:CC | catalytic complex |
| RV10 | 2.82E-05 | 254 | 227 | 17 | 0.074889868 | 0.066929134 | GO:0098798 | GO:CC | mitochondrial protein-containing complex |
| RV10 | 5.00E-05 | 296 | 227 | 18 | 0.079295154 | 0.060810811 | GO:0019866 | GO:CC | organelle inner membrane |
| RV10 | 5.08E-05 | 9673 | 227 | 172 | 0.757709251 | 0.017781454 | GO:0043227 | GO:CC | membrane-bounded organelle |
| RV10 | 5.62E-05 | 436 | 227 | 22 | 0.0969163 | 0.050458716 | GO:0005740 | GO:CC | mitochondrial envelope |
| RV10 | 6.32E-05 | 403 | 227 | 21 | 0.092511013 | 0.052109181 | GO:0031966 | GO:CC | mitochondrial membrane |
| RV10 | 0.000145253 | 10760 | 227 | 184 | 0.810572687 | 0.017100372 | GO:0043229 | GO:CC | intracellular organelle |
| RV10 | 0.000328495 | 11014 | 227 | 186 | 0.81938326 | 0.016887598 | GO:0043226 | GO:CC | organelle |
| RV10 | 0.000769499 | 109 | 227 | 10 | 0.044052863 | 0.091743119 | GO:1990204 | GO:CC | oxidoreductase complex |
| RV10 | 0.001090199 | 1604 | 227 | 45 | 0.198237885 | 0.028054863 | GO:0031090 | GO:CC | organelle membrane |
| RV10 | 0.001718773 | 2627 | 227 | 63 | 0.27753304 | 0.023981728 | GO:0005829 | GO:CC | cytosol |
| RV10 | 0.008330592 | 728 | 227 | 25 | 0.110132159 | 0.034340659 | GO:0031967 | GO:CC | organelle envelope |
| RV10 | 0.008330592 | 728 | 227 | 25 | 0.110132159 | 0.034340659 | GO:0031975 | GO:CC | envelope |
| RV10 | 0.026555763 | 3979 | 227 | 81 | 0.356828194 | 0.020356874 | GO:0031974 | GO:CC | membrane-enclosed lumen |
| RV10 | 0.026555763 | 3979 | 227 | 81 | 0.356828194 | 0.020356874 | GO:0070013 | GO:CC | intracellular organelle lumen |
| RV10 | 0.026555763 | 3979 | 227 | 81 | 0.356828194 | 0.020356874 | GO:0043233 | GO:CC | organelle lumen |
| RV10 | 0.029385964 | 55 | 227 | 6 | 0.026431718 | 0.109090909 | GO:0031902 | GO:CC | late endosome membrane |
| RV10 | 9.32E-05 | 5159 | 219 | 103 | 0.470319635 | 0.01996511 | GO:0003824 | GO:MF | catalytic activity |
| RV10 | 0.000583846 | 1775 | 219 | 47 | 0.214611872 | 0.026478873 | GO:0000166 | GO:MF | nucleotide binding |
| RV10 | 0.000583846 | 1775 | 219 | 47 | 0.214611872 | 0.026478873 | GO:1901265 | GO:MF | nucleoside phosphate binding |
| RV10 | 0.00108786 | 1872 | 219 | 48 | 0.219178082 | 0.025641026 | GO:1901363 | GO:MF | heterocyclic compound binding |
| RV10 | 0.018290148 | 706 | 219 | 23 | 0.105022831 | 0.032577904 | GO:0016491 | GO:MF | oxidoreductase activity |
| RV10 | 0.039421159 | 37 | 219 | 5 | 0.02283105 | 0.135135135 | GO:0019205 | GO:MF | nucleobase-containing compound kinase activity |
| RV10 | 0.000258465 | 401 | 87 | 23 | 0.264367816 | 0.057356608 | HP:0001941 | HP | Acidosis |
| RV10 | 0.00090417 | 2089 | 87 | 59 | 0.67816092 | 0.028243179 | HP:0002060 | HP | Abnormal cerebral morphology |
| RV10 | 0.000918263 | 430 | 87 | 23 | 0.264367816 | 0.053488372 | HP:0004360 | HP | Abnormality of acid-base homeostasis |

| | | | | | | | | | |
|---|---|---|---|---|---|---|---|---|---|
| RV10 | 0.001638636 | 2120 | 87 | 59 | 0.67816092 | 0.027830189 | HP:0100547 | HP | Abnormal forebrain morphology |
| RV10 | 0.003261069 | 181 | 87 | 14 | 0.16091954 | 0.077348066 | HP:0003128 | HP | Lactic acidosis |
| RV10 | 0.00658438 | 192 | 87 | 14 | 0.16091954 | 0.072916667 | HP:0003287 | HP | Abnormality of mitochondrial metabolism |
| RV10 | 0.015864269 | 207 | 87 | 14 | 0.16091954 | 0.06763285 | HP:0002151 | HP | Increased serum lactate |
| RV10 | 0.02453357 | 215 | 87 | 14 | 0.16091954 | 0.065116279 | HP:0012103 | HP | Abnormality of the mitochondrion |
| RV10 | 0.029907124 | 1942 | 87 | 53 | 0.609195402 | 0.027291452 | HP:0001263 | HP | Global developmental delay |
| RV15 | 0.005490754 | 7299 | 104 | 68 | 0.653846154 | 0.009316345 | GO:0005737 | GO:CC | cytoplasm |
| RV15 | 0.012904039 | 150 | 104 | 7 | 0.067307692 | 0.046666667 | GO:0000139 | GO:CC | Golgi membrane |
| RV15 | 0.019598006 | 2533 | 104 | 32 | 0.307692308 | 0.012633241 | GO:0012505 | GO:CC | endomembrane system |
| RV15 | 0.015969253 | 5159 | 110 | 54 | 0.490909091 | 0.010467145 | GO:0003824 | GO:MF | catalytic activity |
| RV15 | 0.026458604 | 2001 | 110 | 28 | 0.254545455 | 0.013993003 | GO:0016740 | GO:MF | transferase activity |
| RV25 | 2.49E-06 | 458 | 75 | 15 | 0.2 | 0.032751092 | GO:0030036 | GO:BP | actin cytoskeleton organization |
| RV25 | 7.99E-06 | 960 | 75 | 20 | 0.266666667 | 0.020833333 | GO:0007010 | GO:BP | cytoskeleton organization |
| RV25 | 1.54E-05 | 524 | 75 | 15 | 0.2 | 0.028625954 | GO:0030029 | GO:BP | actin filament-based process |
| RV25 | 1.65E-05 | 4205 | 75 | 42 | 0.56 | 0.009988109 | GO:0048856 | GO:BP | anatomical structure development |
| RV25 | 3.52E-05 | 4492 | 75 | 43 | 0.573333333 | 0.009572573 | GO:0032502 | GO:BP | developmental process |
| RV25 | 0.00027496 | 4050 | 75 | 39 | 0.52 | 0.00962963 | GO:0048522 | GO:BP | positive regulation of cellular process |
| RV25 | 0.001035234 | 287 | 75 | 10 | 0.133333333 | 0.034843206 | GO:0007015 | GO:BP | actin filament organization |
| RV25 | 0.00113676 | 4447 | 75 | 40 | 0.533333333 | 0.008994828 | GO:0048518 | GO:BP | positive regulation of biological process |
| RV25 | 0.002540281 | 3317 | 75 | 33 | 0.44 | 0.009948749 | GO:0007275 | GO:BP | multicellular organism development |
| RV25 | 0.003067121 | 1785 | 75 | 23 | 0.306666667 | 0.012885154 | GO:0050793 | GO:BP | regulation of developmental process |
| RV25 | 0.003670848 | 500 | 75 | 12 | 0.16 | 0.024 | GO:0097435 | GO:BP | supramolecular fiber organization |
| RV25 | 0.005244362 | 2754 | 75 | 29 | 0.386666667 | 0.010530138 | GO:0048731 | GO:BP | system development |
| RV25 | 0.01338898 | 2401 | 75 | 26 | 0.346666667 | 0.010828821 | GO:0023051 | GO:BP | regulation of signaling |
| RV25 | 0.013399042 | 1256 | 75 | 18 | 0.24 | 0.01433121 | GO:0006915 | GO:BP | apoptotic process |
| RV25 | 0.017433489 | 1977 | 75 | 23 | 0.306666667 | 0.011633789 | GO:0009653 | GO:BP | anatomical structure morphogenesis |

| | | | | | | | | | |
|---|---|---|---|---|---|---|---|---|---|
| RV25 | 0.017599677 | 2758 | 75 | 28 | 0.373333333 | 0.010152284 | GO:0048583 | GO:BP | regulation of response to stimulus |
| RV25 | 0.019399592 | 2140 | 75 | 24 | 0.32 | 0.011214953 | GO:0051239 | GO:BP | regulation of multicellular organismal process |
| RV25 | 0.025986493 | 1318 | 75 | 18 | 0.24 | 0.013657056 | GO:0012501 | GO:BP | programmed cell death |
| RV25 | 0.025986493 | 1318 | 75 | 18 | 0.24 | 0.013657056 | GO:0008219 | GO:BP | cell death |
| RV25 | 0.035051619 | 338 | 75 | 9 | 0.12 | 0.026627219 | GO:0043065 | GO:BP | positive regulation of apoptotic process |
| RV25 | 0.043966556 | 2403 | 75 | 25 | 0.333333333 | 0.010403662 | GO:0010646 | GO:BP | regulation of cell communication |
| RV25 | 0.044859385 | 2094 | 75 | 23 | 0.306666667 | 0.010983763 | GO:0009966 | GO:BP | regulation of signal transduction |
| RV25 | 0.048512644 | 5110 | 75 | 40 | 0.533333333 | 0.007827789 | GO:0032501 | GO:BP | multicellular organismal process |
| RV25 | 0.000794513 | 135 | 77 | 7 | 0.090909091 | 0.051851852 | GO:0030496 | GO:CC | midbody |
| RV25 | 0.015061033 | 3 | 77 | 2 | 0.025974026 | 0.666666667 | GO:0035189 | GO:CC | Rb-E2F complex |
| RV25 | 0.017076917 | 3202 | 77 | 30 | 0.38961039 | 0.009369144 | GO:0071944 | GO:CC | cell periphery |
| RV25 | 0.025055096 | 20 | 77 | 3 | 0.038961039 | 0.15 | GO:0005834 | GO:CC | heterotrimeric G-protein complex |
| RV25 | 0.025055096 | 20 | 77 | 3 | 0.038961039 | 0.15 | GO:1905360 | GO:CC | GTPase complex |
| RV25 | 0.034707712 | 57 | 77 | 4 | 0.051948052 | 0.070175439 | GO:0019898 | GO:CC | extrinsic component of membrane |
| RV25 | 3.20E-06 | 281 | 79 | 12 | 0.151898734 | 0.042704626 | GO:0003779 | GO:MF | actin binding |
| RV25 | 5.14E-05 | 694 | 79 | 16 | 0.202531646 | 0.023054755 | GO:0008092 | GO:MF | cytoskeletal protein binding |
| RV25 | 0.000107332 | 7099 | 79 | 56 | 0.708860759 | 0.007888435 | GO:0005515 | GO:MF | protein binding |
| RV32 | 0.026967734 | 691 | 47 | 10 | 0.212765957 | 0.01447178 | GO:0007167 | GO:BP | enzyme-linked receptor protein signaling pathway |
| RV32 | 0.034555213 | 427 | 47 | 8 | 0.170212766 | 0.018735363 | GO:0007169 | GO:BP | transmembrane receptor protein tyrosine kinase signaling pathway |
| RV32 | 0.016422123 | 440 | 50 | 8 | 0.16 | 0.018181818 | GO:0008134 | GO:MF | transcription factor binding |
| RV37 | 1.64E-09 | 24 | 27 | 6 | 0.222222222 | 0.25 | GO:0003009 | GO:BP | skeletal muscle contraction |
| RV37 | 2.35E-08 | 36 | 27 | 6 | 0.222222222 | 0.166666667 | GO:0050881 | GO:BP | musculoskeletal movement |
| RV37 | 2.80E-08 | 37 | 27 | 6 | 0.222222222 | 0.162162162 | GO:0050879 | GO:BP | multicellular organismal movement |
| RV37 | 7.39E-07 | 197 | 27 | 8 | 0.296296296 | 0.040609137 | GO:0006936 | GO:BP | muscle contraction |
| RV37 | 4.86E-06 | 250 | 27 | 8 | 0.296296296 | 0.032 | GO:0003012 | GO:BP | muscle system process |
| RV37 | 3.28E-05 | 116 | 27 | 6 | 0.222222222 | 0.051724138 | GO:0006941 | GO:BP | striated muscle contraction |

| | | | | | | | | | |
|---|---|---|---|---|---|---|---|---|---|
| RV37 | 8.50E-05 | 136 | 27 | 6 | 0.222222222 | 0.044117647 | GO:0050905 | GO:BP | neuromuscular process |
| RV37 | 0.00078787 | 104 | 27 | 5 | 0.185185185 | 0.048076923 | GO:0006937 | GO:BP | regulation of muscle contraction |
| RV37 | 0.000805973 | 11 | 27 | 3 | 0.111111111 | 0.272727273 | GO:0014819 | GO:BP | regulation of skeletal muscle contraction |
| RV37 | 0.00332719 | 2 | 27 | 2 | 0.074074074 | 1 | GO:0031446 | GO:BP | regulation of fast-twitch skeletal muscle fiber contraction |
| RV37 | 0.00332719 | 2 | 27 | 2 | 0.074074074 | 1 | GO:0031448 | GO:BP | positive regulation of fast-twitch skeletal muscle fiber contraction |
| RV37 | 0.003557331 | 141 | 27 | 5 | 0.185185185 | 0.035460993 | GO:0090257 | GO:BP | regulation of muscle system process |
| RV37 | 0.003911654 | 63 | 27 | 4 | 0.148148148 | 0.063492063 | GO:0006942 | GO:BP | regulation of striated muscle contraction |
| RV37 | 0.009971716 | 3 | 27 | 2 | 0.074074074 | 0.666666667 | GO:0014724 | GO:BP | regulation of twitch skeletal muscle contraction |
| RV37 | 0.009971716 | 3 | 27 | 2 | 0.074074074 | 0.666666667 | GO:0031443 | GO:BP | fast-twitch skeletal muscle fiber contraction |
| RV37 | 0.019923746 | 4 | 27 | 2 | 0.074074074 | 0.5 | GO:0014721 | GO:BP | twitch skeletal muscle contraction |
| RV37 | 0.019923746 | 4 | 27 | 2 | 0.074074074 | 0.5 | GO:0003010 | GO:BP | voluntary skeletal muscle contraction |
| RV37 | 6.30E-06 | 131 | 25 | 6 | 0.24 | 0.045801527 | GO:0030016 | GO:CC | myofibril |
| RV37 | 7.88E-06 | 136 | 25 | 6 | 0.24 | 0.044117647 | GO:0043292 | GO:CC | contractile fiber |
| RV37 | 2.94E-05 | 8 | 25 | 3 | 0.12 | 0.375 | GO:0005861 | GO:CC | troponin complex |
| RV37 | 0.000116408 | 113 | 25 | 5 | 0.2 | 0.044247788 | GO:0030017 | GO:CC | sarcomere |
| RV37 | 0.000149274 | 13 | 25 | 3 | 0.12 | 0.230769231 | GO:0005865 | GO:CC | striated muscle thin filament |
| RV37 | 0.000291393 | 16 | 25 | 3 | 0.12 | 0.1875 | GO:0036379 | GO:CC | myofilament |
| RV37 | 0.001617872 | 7299 | 25 | 22 | 0.88 | 0.003014112 | GO:0005737 | GO:CC | cytoplasm |
| RV37 | 0.0036404 | 36 | 25 | 3 | 0.12 | 0.083333333 | GO:0016529 | GO:CC | sarcoplasmic reticulum |
| RV37 | 0.010457008 | 51 | 25 | 3 | 0.12 | 0.058823529 | GO:0016528 | GO:CC | sarcoplasm |
| RV37 | 0.018701448 | 523 | 25 | 6 | 0.24 | 0.011472275 | GO:0099512 | GO:CC | supramolecular fiber |
| RV37 | 0.019910339 | 529 | 25 | 6 | 0.24 | 0.011342155 | GO:0099081 | GO:CC | supramolecular polymer |
| RV37 | 0.038550085 | 15 | 25 | 2 | 0.08 | 0.133333333 | GO:0033017 | GO:CC | sarcoplasmic reticulum membrane |
| RV37 | 0.000995385 | 582 | 28 | 8 | 0.285714286 | 0.013745704 | GO:0005509 | GO:MF | calcium ion binding |
| RV37 | 0.027045062 | 281 | 28 | 5 | 0.178571429 | 0.017793594 | GO:0003779 | GO:MF | actin binding |
| RV37 | 0.032970138 | 694 | 28 | 7 | 0.25 | 0.010086455 | GO:0008092 | GO:MF | cytoskeletal protein binding |

| | | | | | | | | | |
|---|---|---|---|---|---|---|---|---|---|
| RV37 | 0.000389501 | 6 | 14 | 3 | 0.214285714 | 0.5 | HP:0100830 | HP | Round ear |
| RV37 | 0.00045859 | 809 | 14 | 11 | 0.785714286 | 0.013597033 | HP:0001371 | HP | Flexion contracture |
| RV37 | 0.000600827 | 830 | 14 | 11 | 0.785714286 | 0.013253012 | HP:0034392 | HP | Joint contracture |
| RV37 | 0.000751281 | 67 | 14 | 5 | 0.357142857 | 0.074626866 | HP:0001838 | HP | Rocker bottom foot |
| RV37 | 0.000871627 | 69 | 14 | 5 | 0.357142857 | 0.072463768 | HP:0008365 | HP | Abnormal talus morphology |
| RV37 | 0.000883652 | 861 | 14 | 11 | 0.785714286 | 0.012775842 | HP:0100261 | HP | Abnormal tendon morphology |
| RV37 | 0.00110591 | 229 | 14 | 7 | 0.5 | 0.030567686 | HP:0000160 | HP | Narrow mouth |
| RV37 | 0.002136174 | 35 | 14 | 4 | 0.285714286 | 0.114285714 | HP:0010557 | HP | Overlapping fingers |
| RV37 | 0.004092126 | 41 | 14 | 4 | 0.285714286 | 0.097560976 | HP:0009465 | HP | Ulnar deviation of finger |
| RV37 | 0.004515643 | 42 | 14 | 4 | 0.285714286 | 0.095238095 | HP:0003327 | HP | Axial muscle weakness |
| RV37 | 0.007683323 | 1345 | 14 | 12 | 0.857142857 | 0.008921933 | HP:0034430 | HP | Abnormal joint physiology |
| RV37 | 0.007683323 | 1345 | 14 | 12 | 0.857142857 | 0.008921933 | HP:0011729 | HP | Abnormality of joint mobility |
| RV37 | 0.009043251 | 456 | 14 | 8 | 0.571428571 | 0.01754386 | HP:0003121 | HP | Limb joint contracture |
| RV37 | 0.010185794 | 113 | 14 | 5 | 0.357142857 | 0.044247788 | HP:0001181 | HP | Adducted thumb |
| RV37 | 0.010185794 | 113 | 14 | 5 | 0.357142857 | 0.044247788 | HP:0003557 | HP | Increased variability in muscle fiber diameter |
| RV37 | 0.010728128 | 16 | 14 | 3 | 0.214285714 | 0.1875 | HP:0005272 | HP | Prominent nasolabial fold |
| RV37 | 0.01260063 | 118 | 14 | 5 | 0.357142857 | 0.042372881 | HP:0012084 | HP | Abnormality of skeletal muscle fiber size |
| RV37 | 0.016044913 | 341 | 14 | 7 | 0.5 | 0.020527859 | HP:0100360 | HP | Upper-limb joint contracture |
| RV37 | 0.016683427 | 58 | 14 | 4 | 0.285714286 | 0.068965517 | HP:0003273 | HP | Hip contracture |
| RV37 | 0.018472093 | 19 | 14 | 3 | 0.214285714 | 0.157894737 | HP:0003049 | HP | Ulnar deviation of the wrist |
| RV37 | 0.018472093 | 19 | 14 | 3 | 0.214285714 | 0.157894737 | HP:0005289 | HP | Abnormality of the nasolabial region |
| RV37 | 0.01949381 | 129 | 14 | 5 | 0.357142857 | 0.03875969 | HP:0001850 | HP | Abnormality of the tarsal bones |
| RV37 | 0.020017989 | 227 | 14 | 6 | 0.428571429 | 0.026431718 | HP:0005750 | HP | Lower-limb joint contracture |
| RV37 | 0.026325846 | 65 | 14 | 4 | 0.285714286 | 0.061538462 | HP:0003803 | HP | Type 1 muscle fiber predominance |
| RV37 | 0.02871778 | 1881 | 14 | 13 | 0.928571429 | 0.006911217 | HP:0011805 | HP | Abnormal skeletal muscle morphology |
| RV37 | 0.029708135 | 67 | 14 | 4 | 0.285714286 | 0.059701493 | HP:0001193 | HP | Ulnar deviation of the hand or of fingers of the hand |

| | | | | | | | | | |
|---|---|---|---|---|---|---|---|---|---|
| RV37 | 0.033231153 | 248 | 14 | 6 | 0.428571429 | 0.024193548 | HP:0003701 | HP | Proximal muscle weakness |
| RV37 | 0.035613237 | 1540 | 14 | 12 | 0.857142857 | 0.007792208 | HP:0003549 | HP | Abnormality of connective tissue |
| RV37 | 0.036118791 | 1232 | 14 | 11 | 0.785714286 | 0.008928571 | HP:0001367 | HP | Abnormal joint morphology |
| RV37 | 0.03771539 | 552 | 14 | 8 | 0.571428571 | 0.014492754 | HP:0003272 | HP | Abnormal hip bone morphology |
| RV37 | 0.039556375 | 72 | 14 | 4 | 0.285714286 | 0.055555556 | HP:0100295 | HP | Muscle fiber atrophy |
| RV37 | 0.04255923 | 259 | 14 | 6 | 0.428571429 | 0.023166023 | HP:0004303 | HP | Abnormal muscle fiber morphology |
| RV37 | 0.048137729 | 403 | 14 | 7 | 0.5 | 0.017369727 | HP:0001762 | HP | Talipes equinovarus |
| RV37 | 0.048996246 | 26 | 14 | 3 | 0.214285714 | 0.115384615 | HP:0008368 | HP | Tarsal synostosis |

**Table S13.** Genes for module RV03 that overlap with significant modules from Velotta et al. 2018. Parentheses indicate gene ID differences between the P. maniculatus v1.0 annotation used in Velotta et al. 2018 and the most recent annotation used here (v.2.1.3). We found 19 genes in our module RV03 that overlapped with modules M7 and M8 from Velotta et al. 2018.

| Gene | Velotta et al. 2018 Module ID | Module ID this study |
|---|---|---|
| *Arl15* | M8 | RV03 |
| *Cntrob* | M7 | RV03 |
| *Cpped1* | M7 | RV03 |
| *Dock6* | M8 | RV03 |
| *Dtna* | M7 | RV03 |
| *Hspb2* | M7 | RV03 |
| *Il17rc* | M8 | RV03 |
| ***Irf7*** | M7 | RV03 |
| *Kctd12* | M8 | RV03 |
| *Kiaa1217* | M7 | RV03 |
| *LOC102906896* | M7 | RV03 |
| *LOC102907615* | M7 | RV03 |
| *Lrrc3b* | M7 | RV03 |
| *Pkd2l2* | M7 | RV03 |
| *Tsc2* | M7 | RV03 |
| *Ccbl1 (Kyat1)* | M7 | RV03 |
| *Ftsj2 (Mrm2)* | M7 | RV03 |
| *LOC102909699 (Trim34)* | M7 | RV03 |
| *LOC102928752 (Tfpi)* | M8 | RV03 |
